# A meta-analysis of the gut microbiome in inflammatory bowel disease patients identifies disease-associated small molecules

**DOI:** 10.1101/2024.02.07.579278

**Authors:** Moamen M. Elmassry, Kohei Sugihara, Pranatchareeya Chankhamjon, Francine R. Camacho, Shuo Wang, Yuki Sugimoto, Seema Chatterjee, Lea Ann Chen, Nobuhiko Kamada, Mohamed S. Donia

## Abstract

Changes in the gut microbiome have been associated with several human diseases, but the molecular and functional details underlying these associations remain largely unknown. Here, we performed a multi-cohort analysis of small molecule biosynthetic gene clusters (BGCs) in 5,306 metagenomic samples of the gut microbiome from 2,033 Inflammatory Bowel Disease (IBD) patients and 833 matched healthy subjects and identified a group of Clostridia-derived BGCs that are significantly associated with IBD. Using synthetic biology, we discovered and solved the structures of six fatty acid amides as the products of the IBD-enriched BGCs. Using two mouse models of colitis, we show that the discovered small molecules disrupt gut permeability and exacerbate inflammation in chemically and genetically susceptible mice. These findings suggest that microbiome-derived small molecules may play a role in the etiology of IBD and represent a generalizable approach for discovering molecular mediators of microbiome-host interactions in the context of microbiome-associated diseases.

## Introduction

Several inflammatory (e.g., inflammatory bowel diseases), metabolic (e.g., type 2 diabetes and obesity), and other (e.g., colorectal cancer) diseases have been associated with changes in the gut microbiome’s composition and function (Fan and Pedersen, 2021; Karlsson et al., 2013; Lloyd-Price et al., 2019; Mills et al., 2022; Turnbaugh et al., 2006; Wirbel et al., 2019; Yachida et al., 2019; Zhang et al., 2022). Some of these changes are thought to be causal of disease, but the exact mechanisms by which the gut microbiome may alter host physiology and impact disease progression are still not fully understood. Thus, there is a dire need to identify molecular mediators of microbiome-disease associations, *en route* towards disentangling cause and effect and migrating from an observational to a mechanistic state in microbiome studies (Fischbach, 2018; Maruvada et al., 2017).

Inflammatory Bowel Diseases (IBD), comprising mainly Ulcerative Colitis (UC) and Crohn’s Disease (CD), are complex disorders that have intertwined genetic and environmental origins (Imhann et al., 2018). The hallmark of these diseases is chronic inflammation of the gastrointestinal tract due to a hyperactive immune response (Gao et al., 2018). Several studies have documented notable differences in the human gut microbiome’s composition between IBD patients and healthy individuals (Franzosa et al., 2019; Gevers et al., 2014; Halfvarson et al., 2017; Lloyd-Price et al., 2019; Morgan et al., 2012; Pascal et al., 2017). Such differences are thought to be a consequence of the oxidative environment during IBD-associated inflammation (Morgan et al., 2012), but they have also been shown to directly cause an aberrant immune activity in genetically susceptible animals (Atarashi et al., 2017; Garrett et al., 2007; Nagao-Kitamoto et al., 2016). Beyond taxonomic composition, recent metagenomic, metatranscriptomic, and metabolomic studies have revealed significant differences in several functional pathways encoded and expressed by the IBD microbiome (Chen et al., 2020; Lloyd-Price et al., 2019; Schirmer et al., 2018; Zhang et al., 2022). Moreover, these functional and compositional signatures of the IBD microbiome are important in enhancing or predicting response to both microbiome-targeted (e.g., fecal microbial transplants and antibiotics) and immune-targeted (e.g., biologics) treatments (Ananthakrishnan et al., 2017; Delannoy-Bruno et al., 2021; Federici et al., 2022; Haifer et al., 2022; He et al., 2017; Lee et al., 2021; Lewis et al., 2015; Suskind et al., 2015).

Notably, a few microbiome-derived molecules were shown to be positively or negatively associated with IBD. In one case, a pro-inflammatory polysaccharide produced by *Ruminococcus gnavus* was shown to be associated with disease flares in CD patients (Henke et al., 2019). This polysaccharide-associated inflammation is dependent on toll-like receptor 4. In another case, *Bacteroides*-derived sphingolipids, which are decreased in IBD patients, were found to be important in maintaining homeostasis and their absence caused intestinal inflammation in gnotobiotic mice (Brown et al., 2019). In a third case, the tryptophan derivative indole acrylic acid, produced by several *Peptostreptococcus* sp., was shown to improve intestinal barrier function and mediate anti-inflammatory effects. Interestingly, the biosynthetic genes responsible for the production of this molecule are depleted in the gut microbiome of IBD patients (Wlodarska et al., 2017). These promising examples suggest that bioactive molecules from the human gut microbiome may explain mechanistic details of microbiome-host interactions in IBD, with potential impacts on disease progression and treatment. At the same time, the very few examples elucidated to date highlight the need for more comprehensive and systematic explorations of the role of microbiome-derived bioactive molecules in IBD.

## Results

### A computational pipeline for discovering disease-associated BGCs in human cohorts

We started by devising a computational approach for quantifying the small molecule-coding potential of the gut microbiome in thousands of metagenomic samples and computing statistical enrichment in disease sub-cohorts. While various computational methods have been previously developed by us and others for the identification and quantification of small molecule biosynthetic gene clusters (BGCs) in metagenomic samples of the human microbiome (Pascal Andreu et al., 2021; Sugimoto et al., 2019), complications with the quantification of BGCs hinder the accurate computation of differential enrichment. BGCs are multi-gene elements, where defining accurate boundaries is almost impossible without manual curation or functional characterization. Moreover, BGCs do not only encode biosynthetic enzymes, but they also encode transcriptional regulators, transporters, resistance genes, and mobility determinants – all of which confound accurate quantification in metagenomic data because of their tendency to frequently appear in genomes and metagenomes, even in the absence of the occasionally co-encoded BGC.

To circumvent this problem, we reasoned that quantification based solely on genes encoding biosynthetic enzymes is a valid solution for differential enrichment analysis of BGCs. To this end, we devised the following pipeline. First, from assembled metagenomic sequencing data of each subject’s gut microbiome in all cohorts (scaffolds ≥ 5 Kbps), BGCs are identified using antiSMASH, an unbiased tool for discovering and annotating diverse small molecule BGCs (Blin et al., 2019). Second, BGCs are de-replicated to produce a non-redundant set of BGCs. Third, from all BGCs, a non-redundant set of open reading frames (ORFs) encoding core biosynthetic enzymes are extracted (CB-ORFs). Finally, the extracted ORFs are used as a database for read mapping and the abundance of CB-ORFs is used as a proxy to evaluate BGC enrichment. Enrichment can be in either prevalence (i.e., a statistically significant difference in the percentage of samples harboring the BGC of interest between two sub-cohorts), abundance (i.e., a statistically significant difference in the per-sample abundance of the BGC of interest between two sub-cohorts), or both (**Figure 1**).

**Figure 1.**
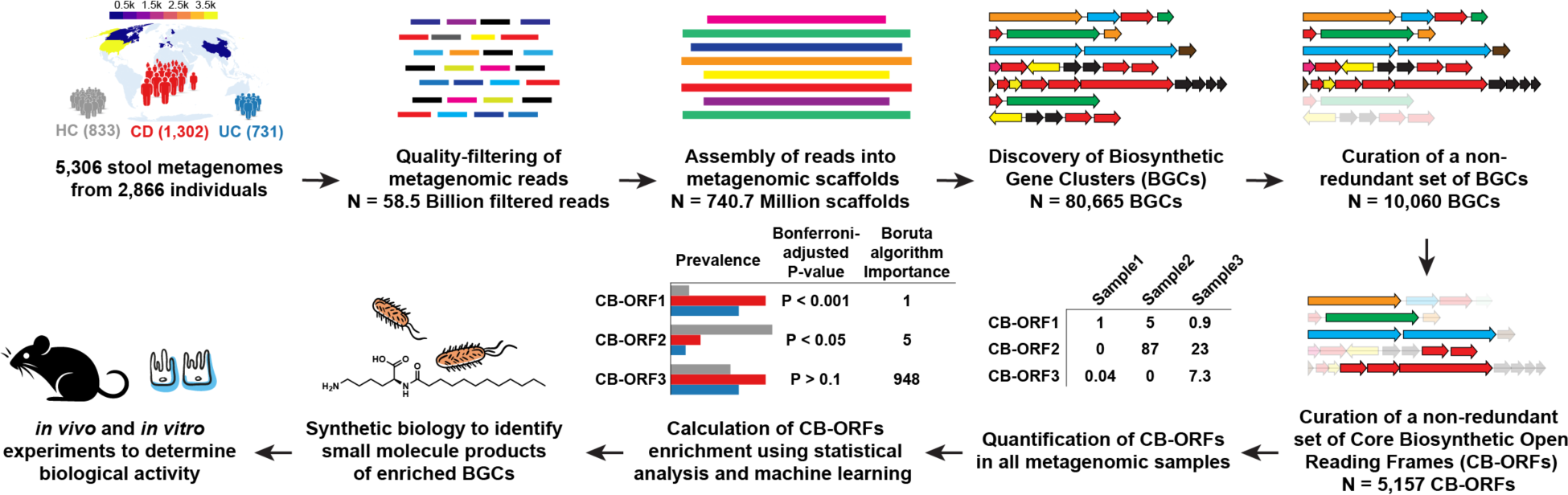
A systematic analysis workflow to identify disease-associated small molecules. A meta-analysis of stool metagenomes from IBD cohorts originating from several geographical regions. The heatmap indicates the number of samples per country. Numbers per IBD sub-cohort (Healthy controls, HC; Crohn’s Disease, CD; or Ulcerative Colitis, UC) indicate the number of subjects included in the analysis. Details of each step are described in the main text and Methods.

### Systematic analysis of small molecule BGCs in the IBD microbiome

We applied this computational strategy to metagenomic sequencing data from the gut microbiome of IBD patients (UC and CD) and matching healthy controls (HC) (**Figure 1**). Since individual studies are subject to technical batch effects and demographical or geographical differences, we decided to employ a multi-cohort analysis and focus on signals that are strongly detected across cohorts. From 18 published studies, including large cohorts such as Integrative Human Microbiome Project (iHMP-IBD) and Metagenomics of the Human Intestinal Tract study (MetaHIT) and two initiatives (IBD Plexus SPARC and RISK), we collated 5,306 samples (1,323 HC, 2,476 CD, and 1,507 UC) from 2,866 subjects (833 HC, 1,302 CD, and 731 UC), originating mainly from four countries (United States, Spain, Netherlands, and China) (**Figure 1**, **Table 1, Data S1, Methods**) (Ananthakrishnan et al., 2017; Connors et al., 2020; Gevers et al., 2014; Hall et al., 2017; He et al., 2017; Lee et al., 2021; Lewis et al., 2015; Li et al., 2014; Lloyd-Price et al., 2019; Muniz Pedrogo et al., 2019; Nielsen et al., 2014; Qin et al., 2010; Schirmer et al., 2018; Turner et al., 2020; Vaughn et al., 2016; Weng et al., 2019). As described above, we quality-filtered the metagenomic reads, assembled them into metagenomic scaffolds, identified the encoded BGCs, then used our ORF-targeted approach to compute the abundance of each BGC in each sample. Overall, we identified 10,060 BGCs (**Data S2**), harboring 199,333 ORFs, of which 5,157 are unique CB-ORFs (**Data S1, Data S3**). BGCs ranged in prevalence from 0.04% to 92.8% of all subjects, with an average prevalence of 10%. In this overall prevalence calculation and all subsequent quantitative analyses, we only used samples with high sequencing depths (≥10 million high-quality reads), which reduced the number of analyzed samples to 3,496 (885 HC, 1,677 CD, and 934 UC) from 2,274 subjects (567 HC, 1,082 CD, and 625 UC).

**Table 1.**
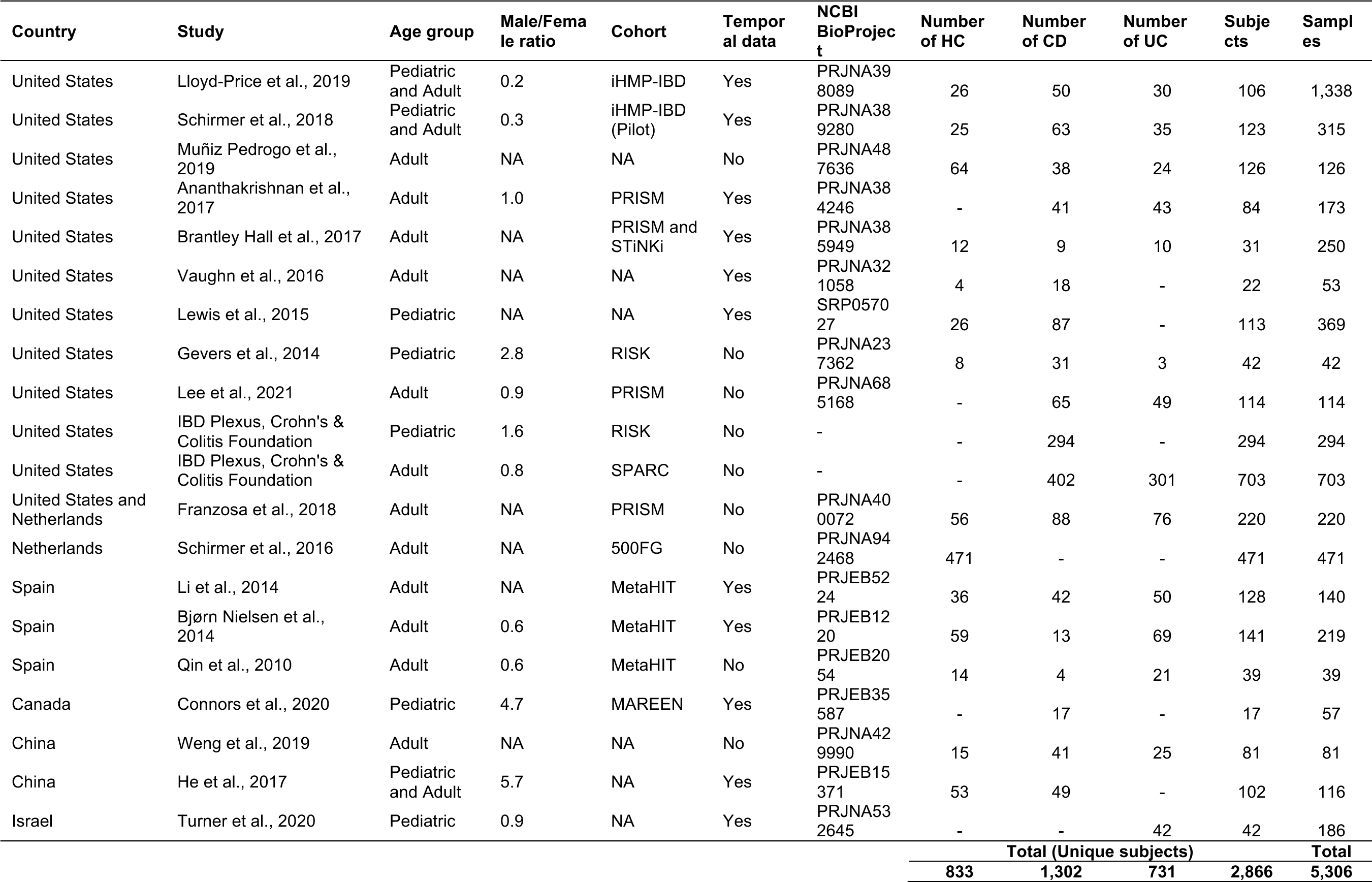
Description of the metagenomic datasets included in the meta-analysis.

Next, we compared the total number of BGCs detected in metagenomic samples of each sub-cohort (i.e., HC, CD, and UC). We found that the median number of BGCs detected in HC was significantly larger than that detected in CD (347 versus 243, P = 3.3×10^-97^, Bonferroni-corrected Dunn’s test following a Kruskal–Wallis test of P = 3.4×10^-98^) or UC (347 versus 282, P = 2.8×10^-28^) (**Figure 2A–B**). This is not surprising, given that the overall diversity of the microbiome is known to decrease in IBD (Gevers et al., 2014; Lloyd-Price et al., 2019; Manichanh et al., 2006; Schirmer et al., 2018). The identified BGCs belonged to almost all known small molecule biosynthetic classes, including Non-Ribosomal Peptide Synthetase (NRPS), Polyketide Synthase (PKS), Ribosomally synthesized and Post-translationally modified Peptide (RiPP), saccharide, terpene, and siderophore BGCs, as well as hybrids thereof (**Figure 2B–C, Data S1**). Notably, when mapping the metagenomically identified BGCs to the NCBI RefSeq Genome and Nucleotide collection databases (Haft et al., 2024; Sayers et al., 2024), only 62% of the BGCs mapped to sequenced genomes, suggesting that a sizable portion of gut microbes capable of producing small molecules have not yet been isolated and/or sequenced. The successfully mapped BGCs are harbored by members of all major phyla in the gut microbiome, namely Firmicutes, Bacteroidetes, Proteobacteria, Actinobacteria, and several other phyla in smaller numbers (e.g., Fusobacteria, Verrucomicrobia, Euryarchaeota, and Ascomycota) (**Figure 2C**). Taken together, these results reveal a chemically and phylogenetically diverse repertoire of small molecule BGCs, which are contributed by both sequenced and not yet sequenced members of the gut microbiome.

**Figure 2.**
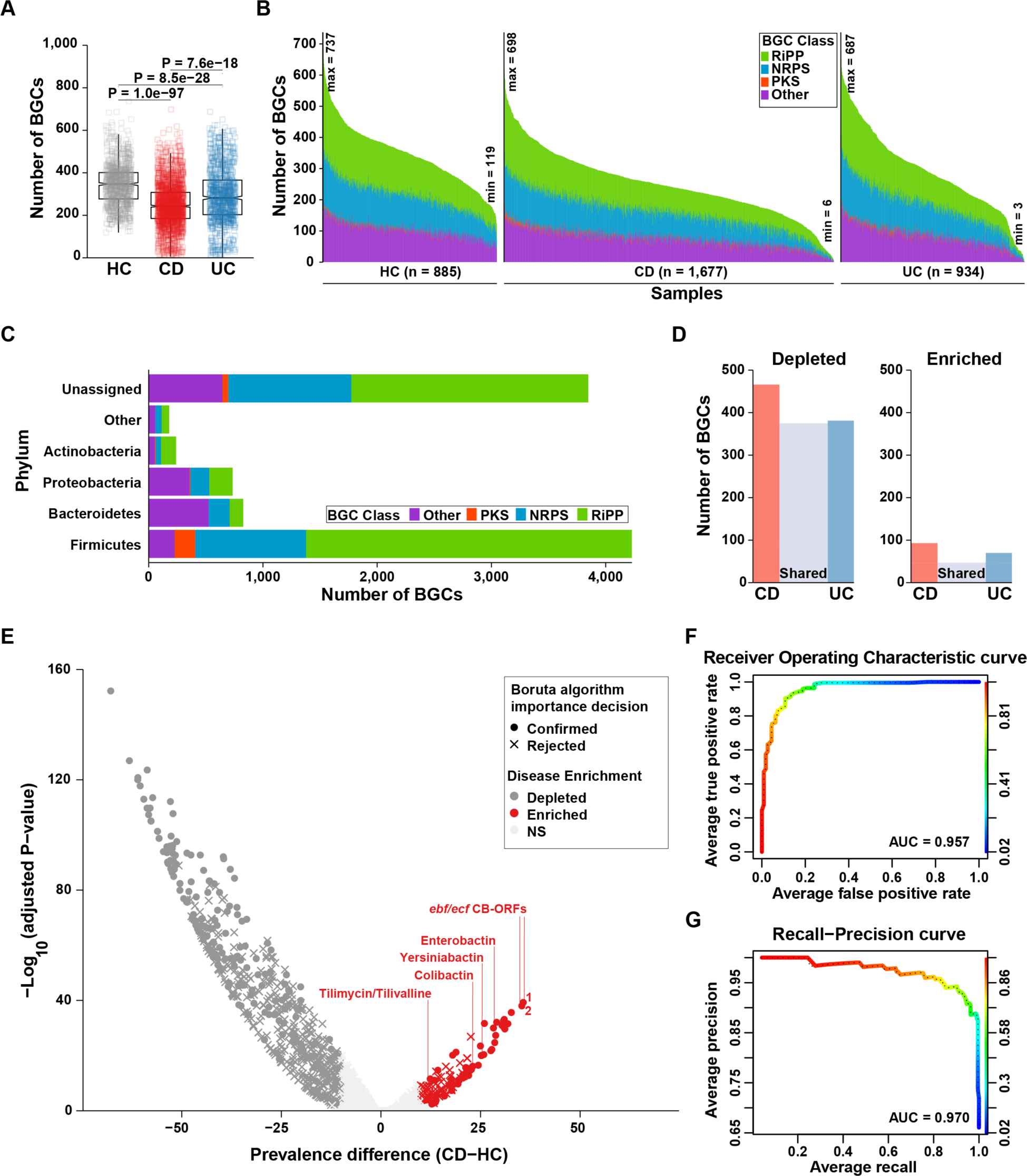
Discovery of IBD-associated small molecule BGCs. **(A)** Number of detected small molecule BGCs in gut metagenomes from healthy and diseased subjects. Statistical significance was determined using Kruskal-Wallis test followed by Dunn’s multiple comparison test and Bonferroni correction. **(B)** A stacked bar plot showing the distribution of all identified small molecule BGCs (grouped by chemical class) in deeply sequenced samples, faceted by health and disease states. **(C)** A stacked bar plot showing the distribution of all identified small molecule BGCs based on the taxonomy of their organism of origin (displayed at the Phylum level) and grouped by their respective chemical class. The category “Other” includes the following Phyla: Fusobacteria, Verrucomicrobia, Euryarchaeota, and Ascomycota (as well as phages). BGCs that did not match to any reference genome in our analysis are labeled as “Unassigned”. **(D)** A Bar plot showing the number of BGCs that are enriched or depleted between healthy and disease states. Two-samples proportion *z*-test followed by Bonferroni correction was used for calculating statistical significance, with P ≤ 0.01 and absolute prevalence difference ≥ 10 as cutoffs. The middle bar labeled as “Shared” indicates the number of BGCs that are commonly detected as statistically significant in both IBD subtypes. **(E)** A volcano plot of CB-ORFs and their prevalence enrichment statistics between HC and CD. Two-samples proportion *z*-test followed by Bonferroni correction was used to calculate statistical significance. CB-ORFs with P ≤ 0.01 and absolute prevalence difference ≥ 10 are highlighted (red if enriched in CD and dark grey if depleted in CD, i.e., enriched in HC). The shape of the points indicates whether a CB-ORF was determined to be important for classification in the machine learning model, based on Boruta algorithm importance. Top enriched CB-ORFs belonging to *ebf* and *ecf* are labelled and the red numbers shown to their right indicate their rank based on Boruta algorithm importance among CB-ORFs enriched in CD. A few of the CD-enriched CB-ORFs belonged to previously characterized BGCs. A single CB-ORF of those BGCs is labeled with their corresponding small molecule products. **(F–G)** A random forest machine learning algorithm trained using CB-ORF abundance profiles of 80% of the samples (456 HC and 862 CD), then tested on never-seen 20% of the samples (112 HC and 218 CD) is able to classify CD and HC samples with high performance (**Methods**). Receiver operating characteristic (ROC) and precision-recall (PR) curves are plotted, and their area under the curve (AUC) values are shown (0.957 and 0.970, respectively). See **Data S1** for the results of a five-fold cross-validation method used to evaluate the performance of the classifier model. All analyses were performed at the level of CB-ORFs, except for panels A, B, and D: due to the fact that a CB-ORF can be present in several BGCs, and to avoid inflating the number of detected or enriched BGCs, we computed representative BGCs per CB-ORFs for the purposes of plotting and calculating enrichment in these figures (and relevant text) (**Methods**).

Next, we sought to examine disease-specific BGC enrichment. To achieve this goal, we compared the prevalence of each CB-ORF (and correspondingly, their parent BGC) at the subject level in a pairwise manner, i.e., HC versus CD and HC versus UC, using two-samples proportion *z*-test followed by Bonferroni correction (**Figure 2D, Data S1**). Using an adjusted P cutoff of ≤ 0.01 and an absolute prevalence difference of ≥ 10, we found 93 and 70 BGCs significantly enriched in CD and UC when compared to HC, respectively (**Figure 2D**). In contrast, 466 and 380 BGCs are significantly depleted in CD and UC relative to HC, respectively. Importantly, 47 and 374 BGCs are enriched and depleted, respectively, in both types of IBD vs. HC (**Figure 2D**). Overall, the majority of depleted BGCs in either CD or UC are depleted in both IBD subtypes, 81% and 98%, respectively. In addition, 51% and 67% of enriched BGCs in either CD or UC are enriched in both, respectively (**Figure 2D**). These results suggest that some small molecule BGCs are enriched generally in IBD, while others are specific to one of the two disease subtypes.

With CD- and UC-enriched BGCs determined, we wondered which members of the microbiome harbor them and what types of molecules they encode. Because Enterobacterales (mainly *Escherichia coli*) are known to be enriched in IBD (Atarashi et al., 2017; Gevers et al., 2014; Schirmer et al., 2018), we expected to identify BGCs derived from this order in our analysis. Indeed, 10 (8%) of the CD-enriched BGCs are assigned to *E. coli*, while only one *E. coli*-derived BGC is enriched in HC. CD-enriched, *E. coli*-derived BGCs include ones that encode for the production of the siderophores yersiniabactin and enterobactin (Laird and Young, 1980; Schubert et al., 2002), as well as the genotoxin colibactin (**Figure 2E**) (Nougayrede et al., 2006), while the HC-enriched, *E. coli*-derived BGC encodes for the production of the antibacterial peptide, microcin J25 (Solbiati et al., 1996). Furthermore, we found that BGCs encoding for the production of the *Klebsiella oxytoca* enterotoxins tilimycin and tilivalline are also enriched in CD (**Figure 2E**) (Dornisch et al., 2017; Schneditz et al., 2014; Unterhauser et al., 2019). Unexpectedly though, we found 55 Firmicutes-derived BGCs that are also enriched in CD, 22 of which derive from members of the class Clostridia, which is generally known to be depleted in CD (Gevers et al., 2014; Pascal et al., 2017). Among the CD-enriched, Clostridia-derived BGCs is a group of two related NRPSs (sharing ∼95% identical CB-ORFs) that are derived from *Clostridium* sp. (*Clostridium clostridioforme* and *Clostridium bolteae*, reclassified now as *Enterocloster clostridioformis* and *Enterocloster bolteae*, respectively) (Haas and Blanchard, 2020). Interestingly, these two BGCs belong to a larger group of NRPSs that have been recently shown to encode diverse fatty acid amides (FAAs) (Chang et al., 2021); therefore, we tentatively named the *E. clostridioformis* and *E. bolteae* FAA BGCs as *ecf* and *ebf*, respectively (**Figure 2E**). UC-enriched BGCs include similar ones to those enriched in CD, i.e., BGCs encoding colibactin, enterobactin, yersiniabactin, and Clostridia-derived NRPSs (*ecf* from *E. clostridioformis* and *ebf* from *E. bolteae*). Taken together, these results indicate that there exist microbiome-derived BGCs that are enriched in CD or UC or both, derived from taxa that are typically known to be enriched in IBD (e.g., *E. coli*), as well as from unexpected taxa known previously to be depleted (e.g., *Clostridium* sp.).

Next, we wondered whether the metagenomic profile of CB-ORFs alone can accurately classify subjects into HC vs. CD or HC vs. UC. To answer this question, we evaluated the ability of a random forest machine learning model trained solely on the CB-ORF’s profiles of known samples to classify unknown samples (i.e., 20% of the samples were not part of the training set, their unique CB-ORFs were not included in the model, and only one sample per subject was used) into UC and HC, or CD and HC (**Methods**). The classifier performed robustly when attempting to separate UC and HC samples: area under the curve (AUC) of the receiver operating characteristic (ROC) curve = 0.932, AUC of the precision-recall (PR) curve = 0.939, and out-of-bag (OOB) estimate of error rate = 13.25% (**Figure S1A–B**). Similarly, a high-performance classifier was trained to separate HC and CD samples: AUC of ROC curve = 0.957, AUC of PR curve = 0.970, and OOB estimate of error rate = 8.88% (**Figure 2F–G**). Satisfyingly, using a five-fold cross-validation method, where we iteratively removed 20% of the samples at random from the training dataset and tested the trained model on the removed samples, we continued to obtain robust classification for both subtypes IBD against HC (AUC of ROC curves ranged from 0.937–0.975 and AUC of PR curves ranged from 0.929–0.986) (**Data S1**). Note that during the cross-validation test, as in the initial model, the removed samples were not part of the training set, their unique CB-ORFs were not included in the model, and only one sample per subject was used. When analyzing the features that best separate CD and HC samples using Boruta algorithm (Kursa and Rudnicki, 2010), 252 (4.9%) CB-ORFs were confirmed to be important features; while in the model separating UC and HC samples, 185 (3.6%) CB-ORFs were confirmed to be important features (**Data S1**). Taken together, these results indicate that the repertoire of microbiome-derived small molecule BGCs is significantly different between IBD and HC, which can be used to successfully classify samples into their respective group.

### Prioritization of disease-enriched BGCs for experimental characterization

Due to the relatively large number of enriched BGCs in CD and UC, we thought to further prioritize BGCs before characterizing their products and biological activity. To achieve this goal, we decided to select BGCs that show significant enrichment in either CD or UC versus HC and are at the same time confirmed to be important and highly ranked by the machine learning classifier. 44 and 16 of the CD-and UC-enriched BGCs met this criterion, respectively (**Figure 2E, Data S1**). Interestingly, the top CD-enriched BGCs are the two related, Clostridia-derived *ebf* and *ecf* mentioned above (**Data S4**): CB-ORFs from *ebf* and *ecf* are the top two CD-enriched CB-ORFs with adjusted P = 5.39×10^-40^ and 9.48×10^-39^, respectively, and are also the top two important CD-enriched features ranked by Boruta algorithm (**Figure 2E**). Moreover, CB-ORFs from *ebf* and *ecf* are among the top three UC-enriched CB-ORFs with adjusted P = 2.01×10^-20^ and 5.68×10^-23^; respectively, and are ranked 9^th^ and 15^th^ among the UC-enriched features by Boruta algorithm (**Data S1**).

Next, we sought to examine the enrichment of *ebf* and *ecf* using a different method. We mapped metagenomic reads from each sample to the entire sequence of the two BGCs and computed two metrics per sample and per subject: a presence or absence status for either of the two BGCs, and an abundance value in Reads Per Kbps per Millions of sequenced reads (RPKM), for each of the two BGCs; the two BGCs share 92% identity on their DNA level and are expected to produce the same products (**Methods**, **Data S1**, **Data S4**). Using this method, we found that while 48% (272 out of 567) of HC subjects harbor *ebf* or *ecf*, 83.7% of CD (906 out of 1,082, P = 8.68×10^-52^, two-samples proportion *z*-test, followed by Bonferroni correction, in comparison to HC) and 78.6% of UC subjects (491 out of 625, P = 2.54×10^-27^, in comparison to HC) do (**Figure 3A, Data S1**). Moreover, the abundance of *ebf* is significantly higher in CD and UC subjects in comparison to HC (6.27×10^-69^ and 3.2×10^-21^, respectively, Bonferroni-corrected Dunn’s multiple comparison test that followed Kruskal-Wallis test P of 3.02×10^-68^) (**Figure 3B**).

**Figure 3.**
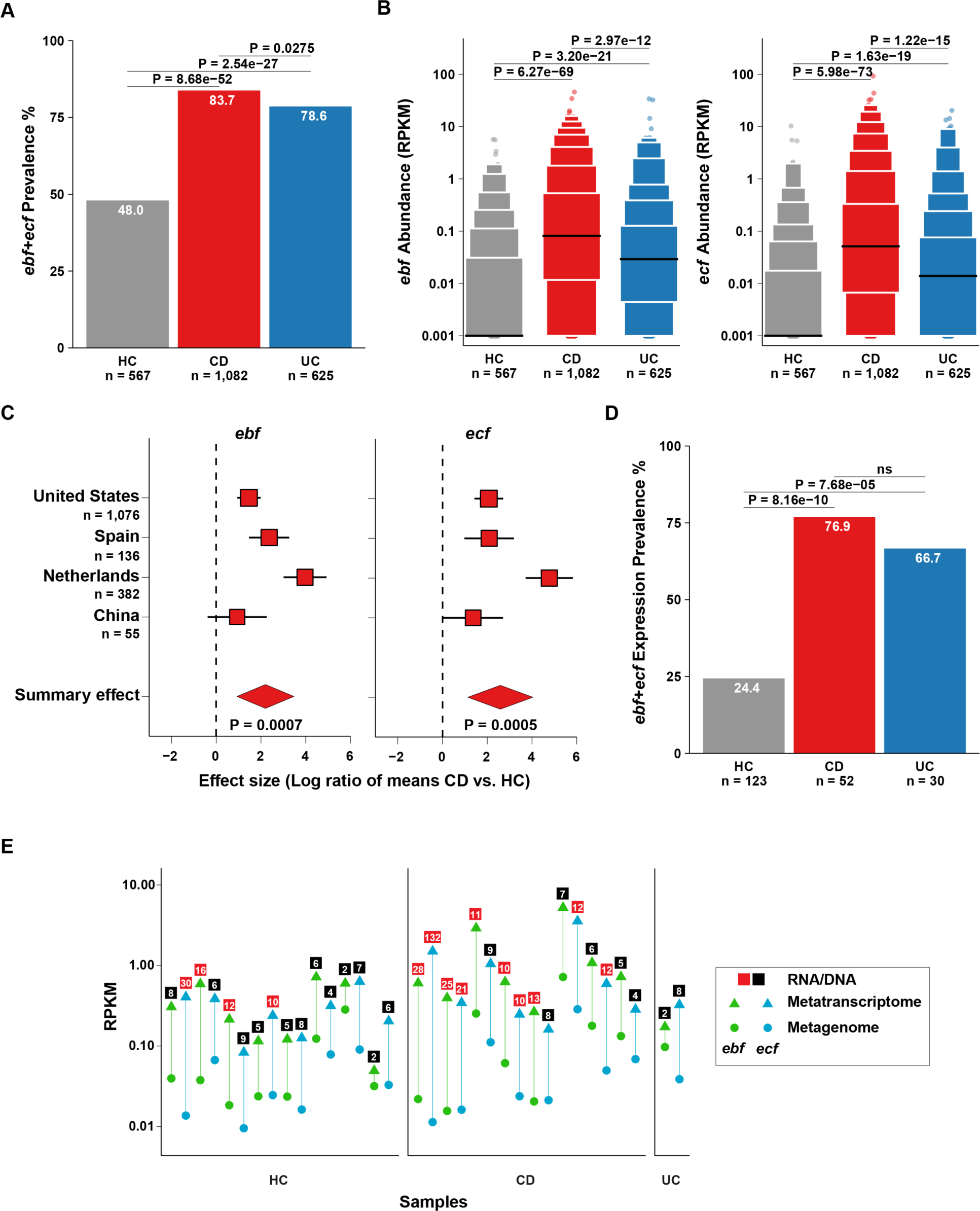
Clostridia-derived *ebf* and *ecf* are enriched and frequently transcribed in the gut microbiome of IBD patients. **(A)** Prevalence of Clostridia-derived *ebf* and *ecf* across HC and IBD (CD and UC) patients. In cases where multiple samples per subject are available, a given subject was deemed “positive” for *ebf* or *ecf* presence if either of the two BGCs was detected in any metagenomic sample from the same subject. Two-samples proportion *z*-test was used to determine statistical significance, followed by Bonferroni correction. **(B)** Abundance of *ebf* and *ecf* in HC, CD and UC patients. Data are presented as boxen (aka letter-value) plot to show distribution with black lines representing the median. Kruskal-Wallis test followed by Dunn’s multiple comparison test and further corrected by Bonferroni method was used to determine statistical significance. In cases where multiple samples per subject are available, the average abundance across samples from the same subject is shown. A pseudo-count of 0.001 was added to all values before plotting. **(C)** Rainforest plot showing effect sizes with confidence intervals of *ebf* and *ecf* enrichment per country. In cases where multiple samples per subject are available, the average abundance across samples from the same subject was used in this analysis. Effect sizes were calculated using a meta-analysis of log ratio of means. A positive shift in effect size indicates enrichment in CD patients, while a negative shift indicates enrichment in HC (or depletion in CD). Random effects model was applied because of study heterogeneity for *ebf* (P = 0.0007, *I*^2^ = 88.4%, *τ*^2^ = 1.5 Cochran’s *Q* = 23.5) and *ecf* (P = 0.0005, *I*^2^ = 88.3%, *τ*^2^ = 1.9, Cochran’s *Q* = 22.8). **(D)** Prevalence of *ebf* and *ecf* transcription across HC, CD, and UC subjects from samples that had paired metagenomic and metatranscriptomic sequencing data available. In cases where multiple samples per subject are available, a given subject was deemed “positive” for *ebf* or *ecf* transcription if either of the two BGCs was detected in any metatranscriptomic sample from the same subject. Two-samples proportion *z*-test was used to determine statistical significance, followed by Bonferroni correction. **(E)** Abundance of *ebf* and *ecf* in paired metagenomic and metatranscriptomic samples. Only samples where RNA/DNA ratio ≥ 5 (for either *ebf* or *ecf*) are shown from the three sub-cohorts, and RNA/DNA ratios ≥ 10 are indicated by red squares (most of them exist in CD). A complete list of samples with associated DNA and RNA RPKM abundances is shown in **Data S1**.

Similarly, the abundance of *ecf* is significantly higher in CD and UC subjects in comparison to HC (P = 5.98×10^-73^ and 1.63×10^-19^, respectively, Bonferroni-corrected Dunn’s multiple comparison test that followed Kruskal-Wallis test P of 8.61×10^-73^) (**Figure 3B**). Finally, to further test enrichment across all cohorts based on their geographical region, we performed a meta-analysis using a random effects model on *ebf* and *ecf* abundance, which revealed a consistent enrichment of *ebf* and *ecf* in CD vs. HC across all countries with a mean effect size estimate of 2.2 (95% confidence interval: 0.9–3.5) and 2.6 (95% confidence interval: 1.1–4.0), and P = 0.0007 and 0.0005, respectively (**Figure 3C**).

While it is clear that *ebf* and *ecf* are enriched in CD and UC patients on the metagenomic level, their *in vivo* relevance depends on whether or not they are being actively expressed under host colonization conditions. To answer this question, we analyzed fecal metatranscriptomic datasets originating from samples where both metagenomic and metatranscriptomic data are available (205 subjects, 123 HC, 52 CD, 30 UC) (**Methods**) (Abu-Ali et al., 2018; Lloyd-Price et al., 2019; Schirmer et al., 2018). Using the same whole-BGC quantification approach described above, we determined that significantly more IBD patients than healthy subjects show active expression of *ebf* or *ecf* in their gut microbiome: 24.4% (30 out of 123) HC subjects, 76.9% (40 out of 52) CD subjects (P = 8.16×10^-10^ and, two-samples proportion *z*-test, followed by Bonferroni correction), and 66.7% (20 out of 30) of UC subjects (P = 7.68×10^-5^, compared to HC) (**Figure 3D**). Intriguingly, there exist paired samples where *ebf* or *ecf* are at low abundance on the metagenomic level while at the same time showing high abundance on the metatranscriptomic level, indicating a robust expression profile even at low abundance of the organism encoding them. This observation was found in samples from all three sub-cohorts, but was more visible in CD (resulting in RNA/DNA RPKM ratios of more than 10 and even more than 100 in one sample) (**Figure 3E**). Consistent with this finding, *ebf*-harboring species, *E. bolteae*, was previously identified as one of a few species with high transcriptional activity (i.e., transcript abundance relative to genomic abundance) in IBD patients with microbiome dysbiosis (Lloyd-Price et al., 2019). These results indicate that *ebf* and *ecf* are indeed expressed under host colonization conditions, at higher frequency in IBD patients, and that their expression levels can vary between samples independently of the abundance of the harboring organism.

### A synthetic biology strategy to access IBD-enriched small molecules

The clear enrichment of *ebf* and *ecf* in IBD motivated us to further explore this association on a functional level. The first step towards this goal is to characterize the small molecule products of *ebf* and *ecf*. Computationally, *ebf* and *ecf* encode six proteins (**Figure 4A**): 1) a 4’-phosphopantetheinyl transferase (PPTase, an enzyme known to activate a peptide carrier protein by transferring a 4’-phosphopantetheine group from coenzyme A to a conserved serine on the carrier protein) (Quadri et al., 1998), 2) a protein with similarity to mammalian lipid transport proteins (Szyperski et al., 1993), 3) a predicted hydrolase, and three stand-alone NRPS domains: 4) a condensation domain (**C**, responsible for amide bond formation), 5) a thiolation domain (**T**, which acts as an acyl/peptide carrier protein for the growing chain), and 6) an adenylation domain (**A**, which activates specific amino acids to be added to the growing chain) (Fischbach and Walsh, 2006) (**Figure 4A**). Interestingly, a recent elegant study undertook a comprehensive *in vitro* biochemical approach to characterize related gut Clostridia-derived NRPS BGCs and showed that their products are fatty acid amides (FAAs) with diverse activities against G-protein coupled receptors (GPCRs) (Chang et al., 2021). Notably, other GPCR-active FAAs, albeit biosynthesized through an unrelated pathway, have also been discovered from more diverse members of the human gut and oral microbiome (Cohen et al., 2017; Cohen et al., 2015). Thus, while the exact small molecule products of *ecf* and *ebf* have not yet been identified, we predicted that they would also be FAAs that are similar in structure to the products of other members of this class. These molecules typically comprise a single amino acid (or a biogenic amine) condensed to a long-chain fatty acid, with the preference to both substrates varying widely depending on the substrate specificity of the A and C domains in the pathway (Chang et al., 2021).

**Figure 4.**
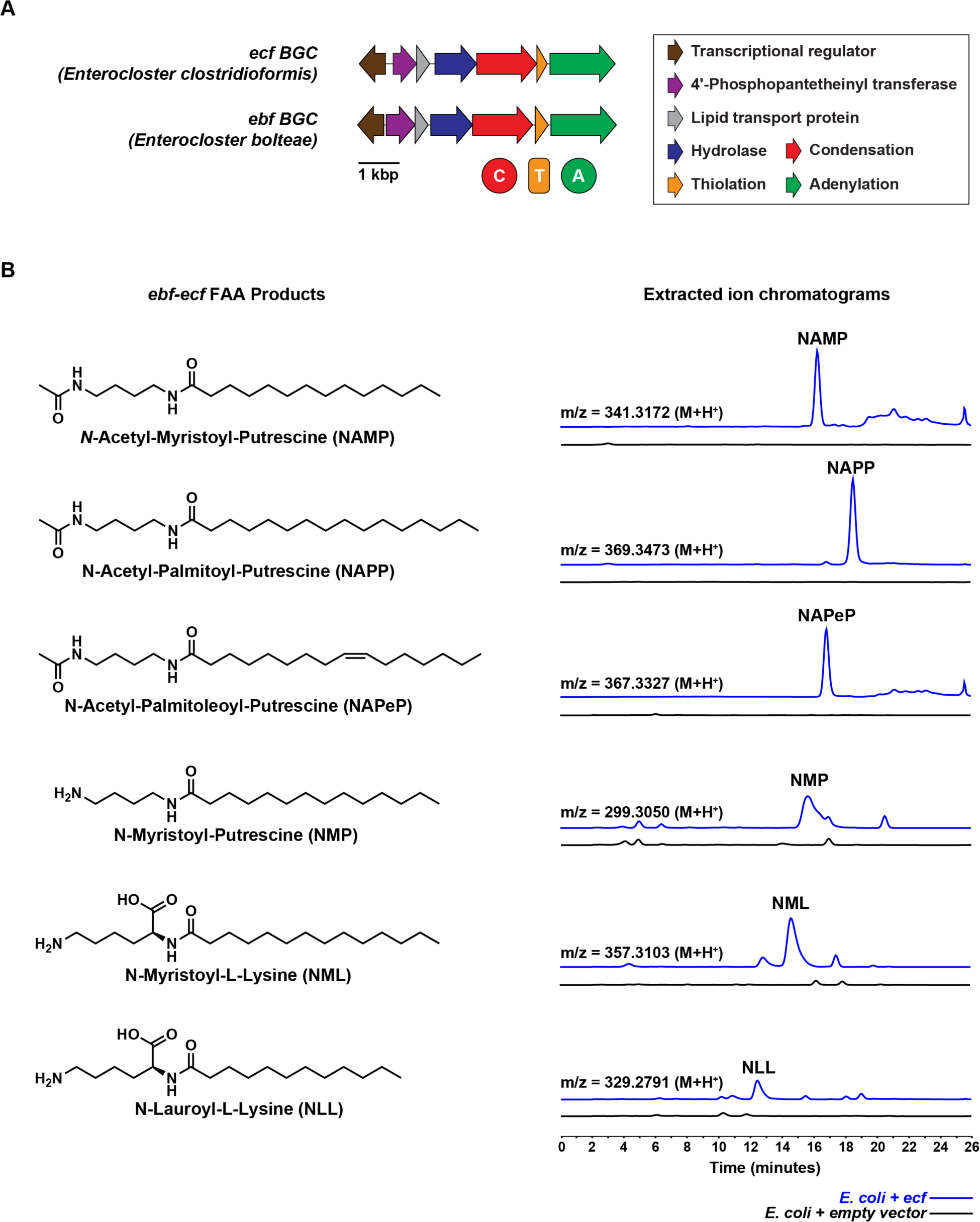
Functional characterization of *ebf* and *ecf* and discovery of their small molecule products. **(A)** Gene and domain architecture of *ecf* (*E. clostridioformis*) and *ebf* (*E. bolteae*). **(B)** Molecular structures of *ebf*-*ecf*-FAAs, the small molecule products of *ebf* and *ecf*. Extracted ion chromatograms (HPLC-HR-MS) for the indicated m/z, obtained from chemical extracts of *E. coli* expressing *ecf* (blue) or a control *E. coli* strain harboring an empty vector (black). MS peaks corresponding to the six discovered FAAs are present in the *ecf* expression line and not the empty vector control.

Owing to the inherent difficulty in genetically manipulating Clostridia species, we decided to undertake a synthetic biology strategy that is based on the heterologous expression of *ebf* and *ecf* in *E. coli*. We cloned the native sequence of *ebf* from *E. bolteae* ATCC BAA-613, and a fully synthetic, codon-optimized version of *ecf* (sequence derived from the MetaHIT metagenomes and closely matching to that of *E. clostridioformis*) into *E. coli* expression vectors under the control of a strong promoter (**Methods**). *E. coli* BL21 strains harboring these two expression constructs with and without an additional PPTase from *Bacillus subtilis* (*sfp*) (Quadri et al., 1998), as well as matching controls harboring empty vectors were cultivated for three days to test whether they produce pathway-specific metabolites. High Performance Liquid Chromatography, coupled with High Resolution Tandem Mass Spectrometry (HPLC-HR-

MS/MS) analysis of chemical extracts from the expression lines and empty vector controls revealed six novel peaks to be commonly produced by both *ebf* and *ecf* in small quantities (**Figure 4B**). These results were corroborated in triplicated experiments. To improve the titer of the produced molecules, we explored several culturing conditions including different media, temperatures, cultivation periods, and oxygenation parameters (**Figure S2**). Using our most optimized conditions, we cultured the *ecf* expression line in large scale and purified all six molecules in pure form (**Methods**).

With the produced molecules in hand, we successfully elucidated their chemical structures using a combination of HR-MS/MS, Nuclear Magnetic Resonance (NMR), and chemical synthesis (**Methods**, **Data S5 and Figures S3**–**5**). As predicted bioinformatically, the products of *ebf* and *ecf* are indeed long-chain FAAs (**Figure 4B**). We determined the structure of the first molecule to be *N*-Acetyl-Myristoyl-Putrescine (NAMP) and the second to be *N*-Acetyl-Palmitoyl-Putrescine (NAPP), and further confirmed them by chemical synthesis; both are novel molecules differing only by two carbons in the fatty acid tail. HR-MS/MS and NMR analyses revealed that the third molecule harbors the same acetyl amide head group but differs than NAPP in having one unsaturation in the fatty acid moiety, thus establishing an *N*-Acetyl-Palmitoleoyl-Putrescine (NAPeP) structure (**Data S5**). The structure of the fourth molecule was elucidated using HR-MS/MS to be *N*-Lauroyl-L-Lysine (NLL), which was confirmed by comparison with a commercially available authentic standard (**Figure S3**). Structures of the fifth and sixth compounds were elucidated using HR-MS/MS to be *N*-myristoyl-putrescine (NMP) and *N*-myristoyl-L-Lysine (NML) and confirmed by chemical synthesis (**Data S5 and Figures S3**–**5**). Taken together, we discovered and solved the structure of six FAAs as products of *ebf* and *ecf* (hereafter *ebf*-*ecf*-FAAs), five of which are novel natural products, while the sixth (NMP) was previously discovered in a functional metagenomic screen of soil-derived environmental DNA, where it was shown to be biosynthesized by an unrelated, non-NRPS pathway (Craig et al., 2010).

Next, we wondered whether the discovered *ebf*-*ecf*-FAAs represent the real products of *ebf* and *ecf* (i.e., match their counterparts produced in the native isolates) or are an artifact of heterologous expression. To answer this question, we cultivated two Clostridia species that have been isolated from human feces: *E. clostridioformis* WAL-7855 (harboring *ecf*) and *E. bolteae* ATCC BAA-613 (harboring *ebf*) in seven different media, then analyzed their chemical extracts using HPLC-HR-MS/MS in comparison to our authentic standards. Indeed, we were able to detect three molecules (NMP, NML, and NLL) from *E. bolteae* in several media, and two molecules (NAMP and NML) from *E. clostridioformis* in one medium each (**Figure S6**). These results confirm that the *ebf*-*ecf*-FAAs obtained using our synthetic biology strategy match ones produced in the native organisms and solidify the link between the produced molecules and their corresponding BGCs without the need for tedious genetic manipulations in Clostridia species.

### A specific subset of FAA-producing BGCs is enriched in the gut microbiome of IBD patients

Because other Clostridia-derived FAA-encoding BGCs similar to *ebf* and *ecf* have been recently characterized (Chang et al., 2021) but not associated with disease, we wondered if they are also enriched in IBD patients. To answer this question, we sought to comprehensively identify BGCs similar to *ebf* and *ecf* in bacterial isolates of the class Clostridia and examine their enrichment or depletion state in CD and UC. First, we downloaded all 8,427 publicly available genome assemblies of Clostridia strains from the RefSeq database (Haft et al., 2024). Then, we generated a phylogenetic tree of Clostridia genomes based on 400 universal marker genes using PhyloPhlAn (Asnicar et al., 2020) (**Figure 5**). Next, we used tblastn to identify genomes harboring closely related homologs to two core biosynthetic genes (C and A) of *ebf* and *ecf* (**Methods**). We found 713 genomes (8.5% of all genomes in the dataset) that harbor NRPS BGCs similar to *ebf* and *ecf*, the majority of which (95.4%) belonged to *Clostridium* cluster XIVa (**Figure 5, Data S1**). Satisfyingly, not only does this list include all of the recently characterized FAA-encoding NRPSs, but it also includes hundreds of uncharacterized ones (Chang et al., 2021). While it is likely that all of these related NRPS BGCs produce FAAs of different flavors, we refrain from such assumption given how promiscuous NRPS enzymes are known to be.

**Figure 5.**
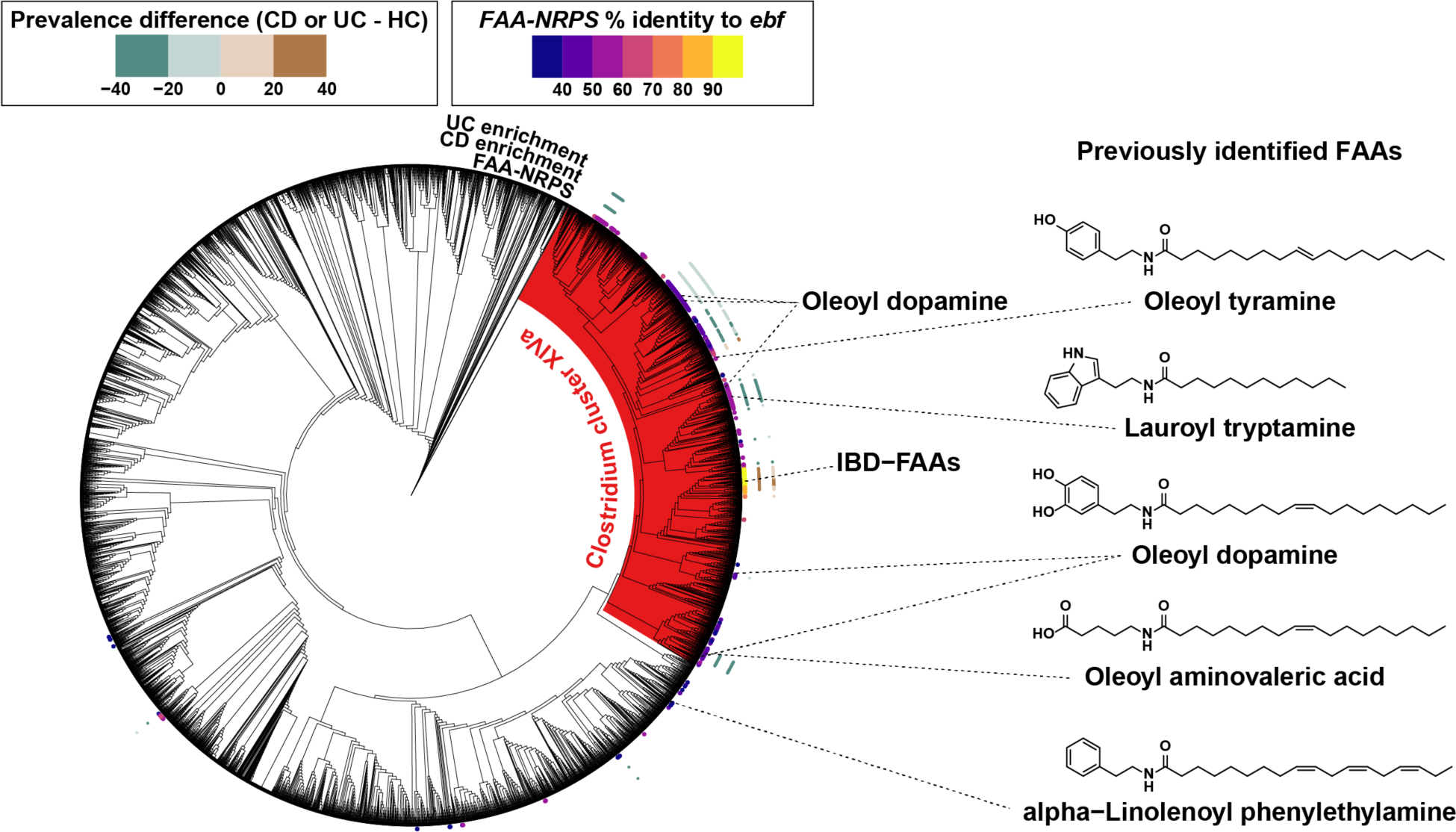
Only specific FAA BGCs are enriched in IBD. A phylogenetic tree of 8,427 strains from the class Clostridia, with publicly available genomes from the RefSeq database. The tree was constructed using PhyloPhlAn, based on a set of 400 universal marker genes. *Clostridium* cluster XIVa clade is highlighted in red. Genomes with BGCs that are homologous to *ebf* and *ecf* are marked in the innermost layer of points, labeled as FAA-NRPSs, and colored by their percent identity to *ebf* as indicated in the color key at the top right. The two outermost layers indicate disease enrichment in CD and UC, colored by prevalence difference (CD or UC - HC) as indicated in the color key at the top left. BGCs with previously identified FAA products are connected to the name and molecular structure of their cognate FAA (Chang et al., 2021).

Next, we leveraged our BGC quantification method to compute the enrichment status of all identified *ebf* and *ecf* homologs in IBD. First, we annotated NRPS BGCs from all positive Clostridia genomes using antiSMASH, then used tblastn to match the identified BGCs to CB-ORFs in the set that we curated from our IBD multi-cohort analysis (≥ 90% protein sequence identity, breadth of coverage ≥ 50%, and E-value < 0.001). Using this criterion, 592 of the 713 NRPS BGCs had representative CB-ORFs in our dataset. Finally, we inferred the enrichment of the matched NRPS BGCs based on the enrichment statistics already computed in our global analysis. Interestingly, 503 of the matched BGCs (85%) were differentially prevalent in IBD (CD or UC) vs. HC: while the majority (402, 67.9%) were depleted in CD and UC, only a small subset of NRPSs (101, 17.1%) were enriched in CD or UC, including *ebf* and *ecf* (**Figure 5**). Since the products of several FAA-encoding NRPSs in our dataset have been recently identified (Chang et al., 2021), we took this analysis one step further and annotated NRPSs, when applicable, with their corresponding small molecules (**Figure 5**). *ecf* and *ebf* are the only IBD-enriched BGCs with characterized products. Interestingly, several known FAA-encoding NRPSs were consistently depleted from both UC and CD in comparison to HC: a BGC from [*Eubacterium*] *rectale* ATCC 33656 that produces lauroyl tryptamine, a BGC from *Coprococcus eutactus* ATCC 27759 that produces oleoyl aminovaleric acid, and three BGCs that produce oleoyl dopamine, from *Blautia wexlerae* AGR2146, *Blautia* sp. Marseille-P2398, and *Clostridium* sp. L2-50. It is important to note that most of the Clostridia-encoded *ebf* and *ecf* homologs identified in our analysis still await characterization, some of which share relatively high nucleotide sequence identity to *ebf* (and *ecf*) and may potentially produce similar products (e.g., NRPSs from *Enterocloster citroniae*, 84.6% identity to *ebf*, and from *Enterocloster aldenensis*, 87.7% identity, the latter of which is enriched in both CD and UC). On the other hand, other BGCs are quite dissimilar (but still related) to *ebf*, and are likely to produce divergent molecules (e.g., *Blautia producta*, 68.1% identity to *ebf*, enriched in CD, and *Mediterraneibacter faecis*, 55.9% identity, depleted in both CD and UC).

### *ebf*-*ecf*-FAAs exacerbate disease in mouse models of colitis

Motivated by the enrichment of *ebf* and *ecf* in IBD, we wondered whether their small molecule products have a role in disease. To answer this question, we sought to test the effects of *ebf*-*ecf*-FAAs in two mouse models of colitis: the dextran sulfate sodium (DSS)-induced colitis model and the IL-10-deficient (IL-10^-/-^) colitis model. For the first model, we treated C57BL/6J mice with a cocktail of antibiotics then colonized them with either an *E. coli* strain expressing *ecf* (*ecf*+) or an isogenic strain harboring an empty vector control (*ecf*-). DSS was administered daily (until day 6) to induce colitis, then mice were sacrificed on day 7 to determine the extent of disease (**Figure 6A**). Despite similar colonization levels of the introduced strains (**Figure S7**), we found that *ecf*+ mice had significantly decreased colon length and increased colon density (weight divided by length) when compared to *ecf*-mice (**Figure 6B, 6C**), both of which are hallmarks of increased inflammation.

**Figure 6.**
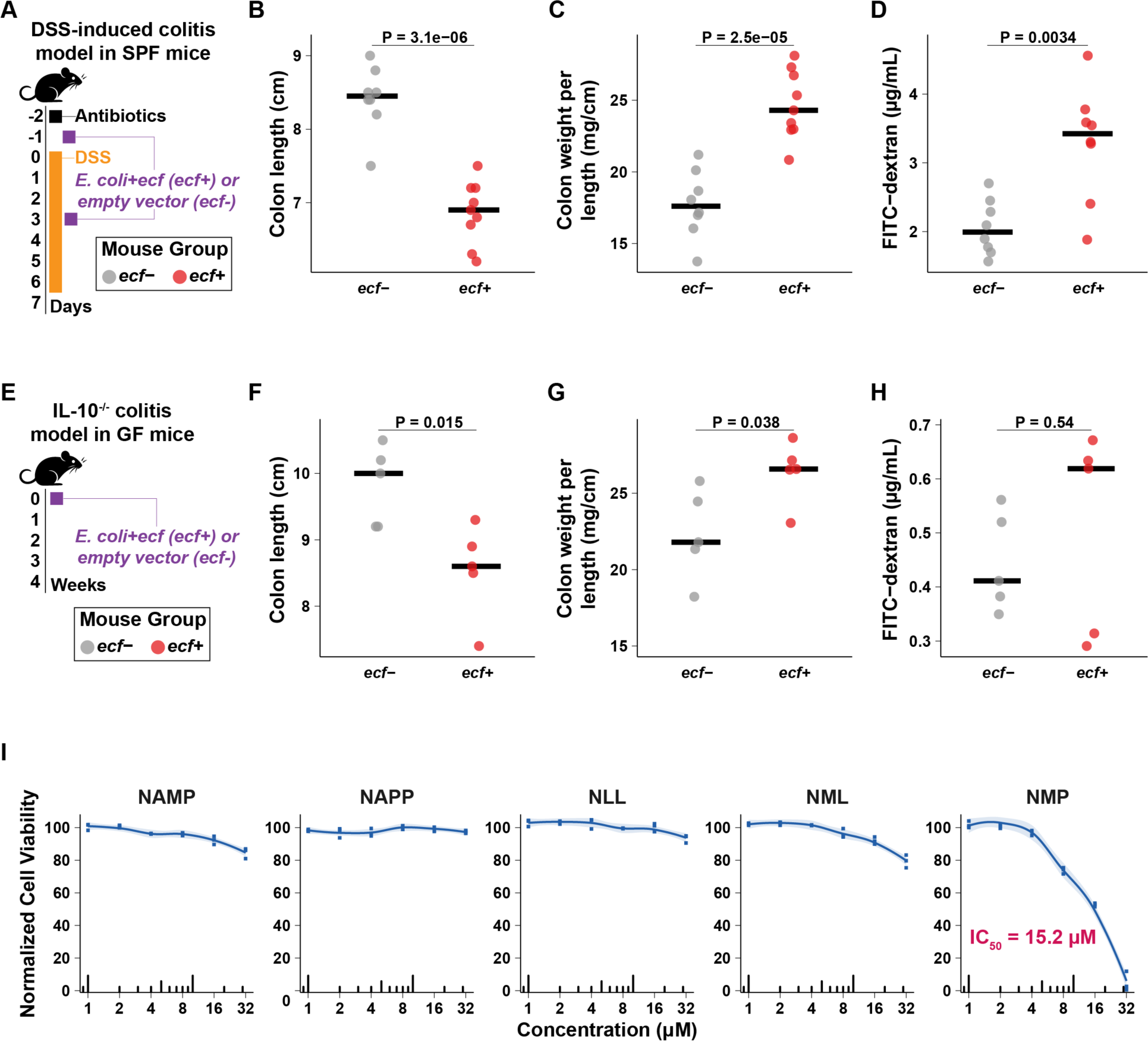
ebf-ecf-FAAs exacerbate disease in mouse models of colitis. **(A)** Timeline for the DSS-induced colitis mouse model experiment. *ecf*+ indicates the group of mice colonized with *E. coli* expressing *ecf*, and *ecf-* indicates the group of mice colonized with *E. coli* harboring an empty vector control. **(B–D)** Comparison between *ecf*+ and *ecf*-DSS-treated mice in: **(B)** colon length, **(C)** colon weight per length, **(D)** Intestinal permeability using FITC-dextran. **(E)** Timeline for the IL-10^-/-^ germ-free mouse model experiment. **(F–H)** Comparison between *ecf*+ and *ecf*-IL- 10^-/-^ gnotobiotic mice in: **(F)** colon length, **(G)** colon weight per length, and **(H)** Intestinal permeability measured using FITC-dextran. Data are presented as individual points, and the median is presented as a horizontal line. Data in the DSS-induced colitis model are collected from two independent experiments. Two-sided Student’s *t*-test was used to determine statistical significance in all comparisons (ns: not statistically significant). **(I)** Cytotoxicity of *ebf*-*ecf*-FAAs was evaluated by measuring relative cell viability of Caco-2 cells after 24 hours of incubation with each of five *ebf*-*ecf*-FAAs (at concentrations 1–32 µM, with each concentration measured in triplicates). Values are normalized to untreated cells. Data are represented as dose-response curves (x-axis is log-scaled) with individual data points shown and curve fitted using loess regression. IC_50_, the half-maximal inhibitory concentration, was calculated for NMP using regression curves fitted to NMP concentration and normalized cell viability, according to the Hill Equation.

Next, we measured intestinal permeability in colonized mice using fluorescein isothiocyanate dextran (FITC-dextran) administration. Interestingly, we found that *ecf*+ mice showed significantly higher intestinal permeability than *ecf*- mice (**Figure 6D**), indicating a disturbed gut barrier integrity – or a “leaky gut”, as commonly observed in human IBD (Mehandru and Colombel, 2021). It is important to note that our results do not determine whether the increased permeability is a cause or a consequence of the exacerbated colitis in *ecf*+ mice. Furthermore, no statistically significant differences in histology scores or cytokine levels were observed between the two groups except for colonic TNF-alpha (P < 0.05), which was significantly increased in *ecf*+ mice (**Figure S7**).

To verify these results in a second model and to determine if they are dependent on the resident microbiome, we colonized two sets of IL-10^-/-^ germ-free mice with either *E. coli* expressing *ecf* or the empty vector control strain (**Figure 6E**). Satisfyingly, we obtained very similar results to those obtained in the DSS colitis model after four weeks of colonization (**Figure S7**): *ecf*+ mice showed a significantly decreased colon length (**Figure 6F**), a significantly increased colon density (**Figure 6G**), and a trend towards increased intestinal permeability (albeit not reaching statistical significance) (**Figure 6H**) when compared to *ecf*- mice. Finally, to determine whether this effect on intestinal permeability may be related to direct toxicity on epithelial cells, as in the case with other enterotoxins (Dornisch et al., 2017; Tse et al., 2017), we measured the viability of cultured colonic epithelial cells (Caco-2) in the presence of various concentrations of each of five *ebf*-*ecf*-FAAs (see **Methods**). Interestingly, NMP showed strong cytotoxicity against the colonic cell line at 32 µM concentration (IC_50_ 15.2 µM) (**Figure 6I**). These cell-based results, combined with the exacerbated colitis phenotype we observed in the two mouse models described above, suggest a potential role of *ebf*-*ecf*-FAAs in IBD pathophysiology that may be dependent on their direct cytotoxicity against epithelial cells and independent of the resident microbiome.

## Discussion

The gut microbiome has been long thought, and in a few cases shown to contribute to human diseases, yet the molecular mediators of these effects are not completely understood. This is especially true for IBD, where strong experimental and epidemiological evidence support a role for the microbiome in the etiology of the disease. Using a systematic approach, we report a series of microbiome-derived small molecules that are strongly associated with IBD. We show that the genes encoding these molecules are significantly enriched and highly expressed in the microbiome of IBD patients (especially in CD samples), and that bacterial strains expressing the same molecules exacerbate disease in mouse models of colitis, likely through an epithelial cytotoxicity mechanism. While our approach is generally unbiased when it comes to the BGCs and molecules identified, broadly applicable to other microbiome-associated diseases, and functionally instead of taxonomically driven, it still suffers from several limitations.

First, we relied exclusively on profiling small molecule BGCs that are identifiable in assembled metagenomic data by well-established BGC-finding algorithms. Small molecules that are produced by yet-unknown classes of biosynthetic pathways, single enzymes, or encoded by BGCs that were not assembled in any of the samples in the analyzed cohort would be missed in our analysis. Nevertheless, we detected thousands of BGCs from highly diverse structural classes and bacterial taxa, many of which do not map to any sequenced organism. As our ability to *de novo* assemble metagenomic datasets gets better, and as new biosynthetic classes are characterized, the comprehensiveness of our approach will increase. Second, an enriched small molecule BGC does not necessitate a causative role in disease, but could also indicate a selection process as a consequence of the disease, where the produced molecule provides a survival or competitive advantage to the producer (Marcelino et al., 2023; Watson et al., 2023). For example, it is often suggested that the expansion of facultative anaerobes in the CD microbiome is a consequence of disease-associated inflammation and oxidative stress (Morgan et al., 2012). Therefore, following the initial computational identification of BGC enrichment with functional experiments in relevant models of the respective disease, as we did in this study, is necessary. Whether the discovered small molecules contribute to the pathogenesis of the disease, or the response to it, or both, studying their effects provides molecular and mechanistic insights into disease-specific, microbiome-host interactions.

The human microbiome is a vast resource for novel small molecules, many of which could explain mechanistic details of microbe-host and microbe-microbe interactions. With thousands of yet-uncharacterized small molecule BGCs encoded by the human microbiome (Cao et al., 2022; Chang et al., 2021; Donia et al., 2014; Guo et al., 2017; Sugimoto et al., 2019), there is a dire need for systematic prioritization of candidates for functional studies. Our approach described here – combining quantitative metagenomic analysis, synthetic biology, and mouse colonization experiments – holds a great promise for both the prioritization and functional characterization of microbiome-derived small molecules that are relevant to human diseases. Although we have used IBD here as a proof-of-principle example, our disease-centric approach is applicable to any microbiome-associated or microbiome-correlated human disease where metagenomic sequencing data is available.

## Star Methods

Detailed methods are provided in the online version of this paper and include the following: KEY RESOURCES TABLE, CONTACT FOR REAGENT AND RESOURCE SHARING, METHOD DETAILS, EXPERIMENTAL DESIGN, AND DATA AND SOFTWARE AVAILABILITY.

## Supplemental Information

Supplemental Information includes seven figures and five data files.

## Author Contributions

M.S.D., M.M.E., F.R.C., P.C. and N.K. designed the study. M.M.E., F.R.C. and S.W. performed the computational analysis. M.M.E., P.C., Y.S., and S.C. performed synthetic biology and molecular biology experiments, heterologous expression, structural elucidation, chemical synthesis and cytotoxicity assays. S.K. performed mouse studies. M.S.D., M.M.E., F.R.C., P.C., S.W., S.K., L.A.C., and N.K. analyzed the data and wrote the manuscript.

## Acknowledgments

We would like to thank Matthew Cahn for assistance with our metagenomic pipeline configuration, Marco Colonna and Luisa Cervantes-Barragan for help with designing mouse experiments, and members of the Donia lab for useful discussions. We would like to acknowledge that the work reported here was substantially performed using the Princeton Research Computing resources at Princeton University. We would like to thank the various research groups who developed software used in this study and maintained them over the years, as well as research groups and centers who deposited their metagenomic data and associated metadata in public repositories, allowing us to use them in this study. Funding for this project has been provided by a Princeton University Intellectual Property Accelerator Fund, an Innovation Award, a Breakthrough Award, and a Phase III Award by the Kenneth Rainin Foundation to M.S.D., National Institute of Health awards DP2AI124441 and R01AI172144 to M.S.D, DK108901 to N.K., DK119544 to L.A.C., and an IBD-Plexus data access grant from the Crohn’s and Colitis Foundation to L.A.C. and M.S.D. The results published here are in whole or part based on data obtained from the IBD Plexus program of the Crohn’s & Colitis Foundation. M.M.E. is supported by the New Jersey Commission on Cancer Research (NJCCR) Postdoctoral Fellowship and the Writing in Science and Engineering Teaching Fellowship.

## Declaration of Interests

M.S.D. is a Scientific Co-Founder and CSO at Pragma Biosciences.

