## Supplemental Figures for "A meta-analysis of the gut microbiome in inflammatory bowel disease patients identifies disease-associated small molecules"

**Table of contents:**

**Figure S1. Evaluation metrics for a machine learning model that classifies UC and HC samples.**
**Related to Figure 2 and Data S1.**

**Figure S2. Optimization of fatty acid amide production by *E. coli*::*ecf* by varying cultivation conditions.**
**Related to Figure 4 and Data S5.**

**Figure S3. Comparison of isolated and synthetic fatty acid amides.**
**Related to Figure 4 and Data S5.**

**Figure S4. Comparison of isolated and synthetic NLL and NML.**
**Related to Figure 4 and Data S5.**

**Figure S5. Synthetic routes for fatty acid amides.**
**Related to Figure 4 and Data S5.**

**Figure S6. Production of fatty acid amides by native isolates of *Clostridium* sp.**
**Related to Figure 4 and Data S5.**

**Figure S7. Effects of *ebf*-*ecf*-FAAs in mouse models of colitis.**
**Related to Figure 6.**

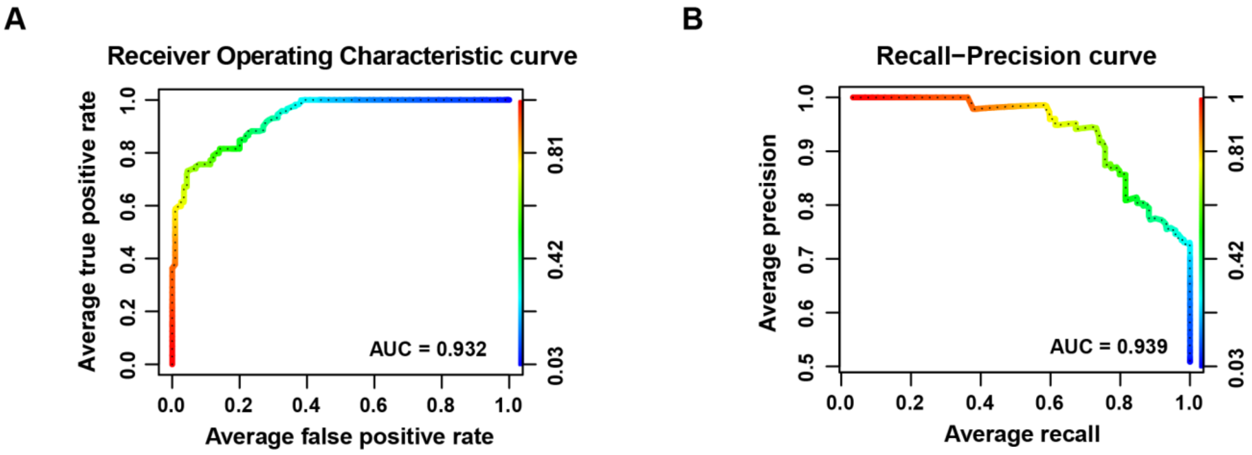


**Figure S1. Evaluation metrics for a machine learning model that classifies UC and HC samples.** A random forest machine learning algorithm trained using CB-ORF abundance profiles of 80% of the samples, then tested on never seen 20% of samples is able to classify UC and HC samples with high performance (**Methods**). Receiver operating characteristic (ROC) and precision-recall (PR) curves are plotted, and their area under the curve (AUC) values are shown (0.932 and 0.939, respectively). See **Data S1** for the results of a five-fold cross-validation method used to evaluate the performance of the classifier model.

**Related to Figure 2 and Data S1.**

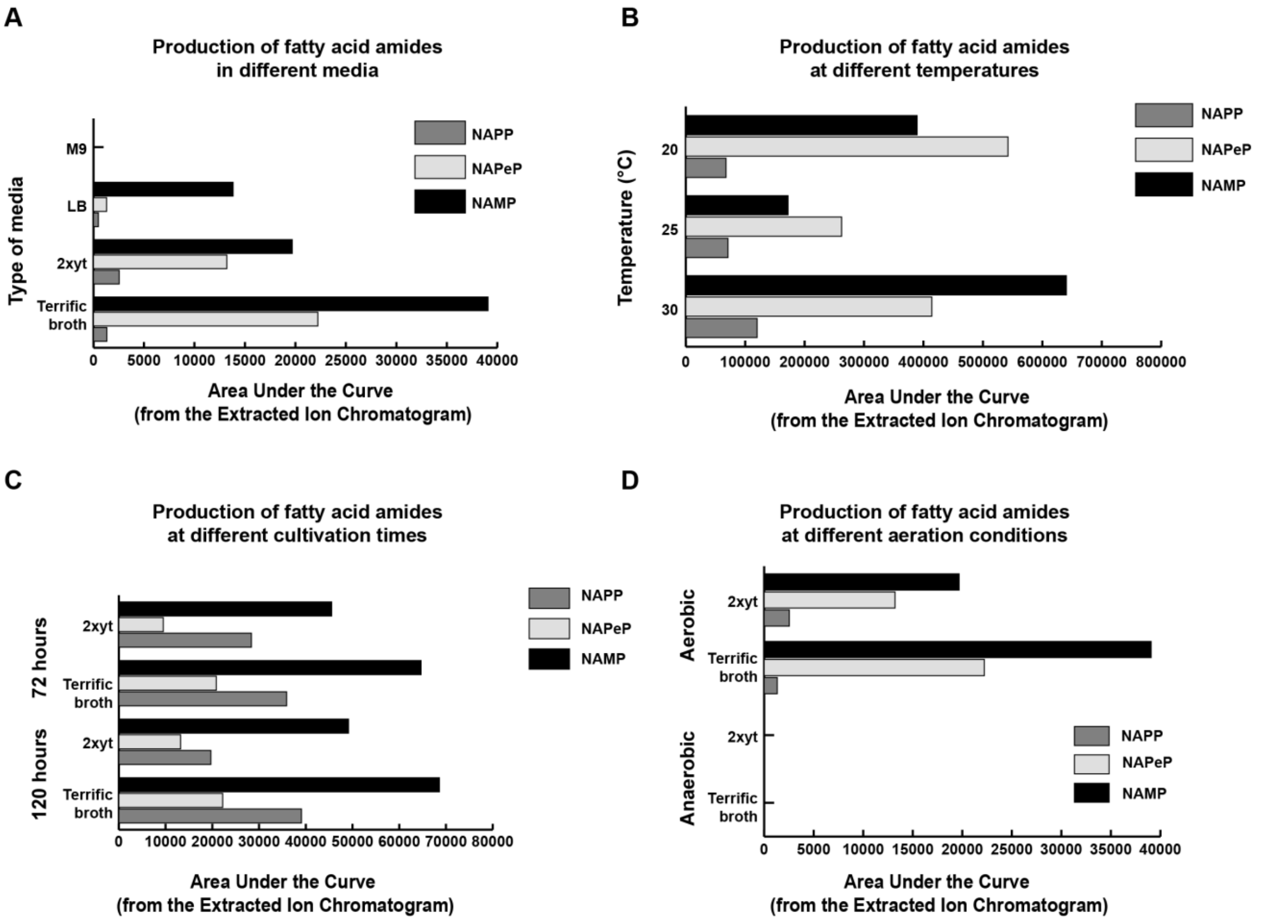


**Figure S2. Optimization of fatty acid amide production by *E. coli*::*ecf* by varying cultivation conditions.** The panels represent the following types of cultivation conditions: (A) Different media. (B) Different temperatures. (C) Different cultivation times. (D) Different aeration conditions.
**Related to Figure 4 and Data S5.**

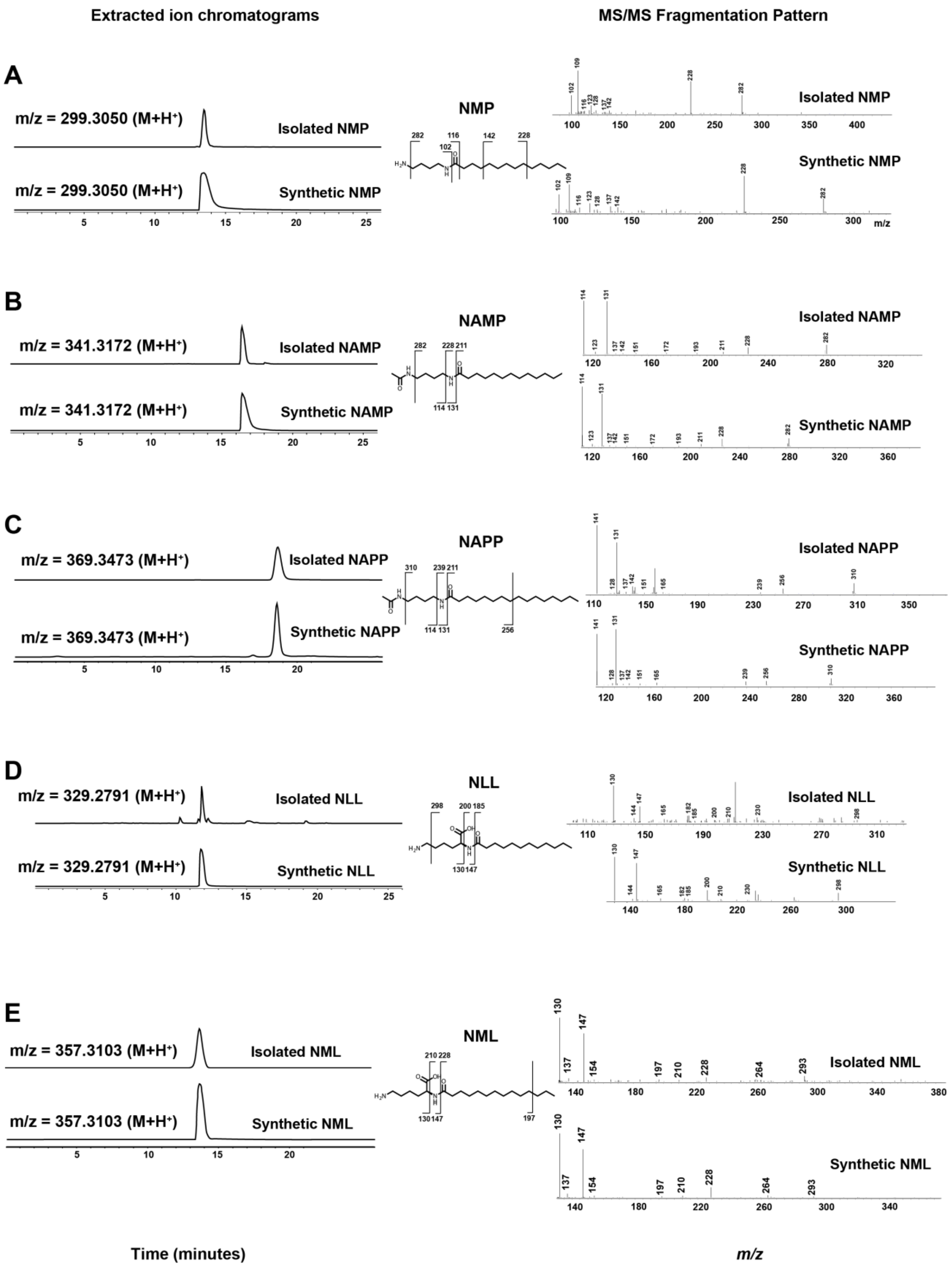
Extracted ion chromatograms
MS/MS Fragmentation Pattern
A
m/z = 299.3050 (M+H+)
Isolated NMP
m/z = 299.3050 (M+H+)
Synthetic NMP
NMP
Isolated NMP
Synthetic NMP
B
m/z = 341.3172 (M+H+)
Isolated NAMP
m/z = 341.3172 (M+H+)
Synthetic NAMP
NAMP
Isolated NAMP
Synthetic NAMP
C
m/z = 369.3473 (M+H+)
Isolated NAPP
m/z = 369.3473 (M+H+)
Synthetic NAPP
NAPP
Isolated NAPP
Synthetic NAPP
D
m/z = 329.2791 (M+H+)
Isolated NLL
m/z = 329.2791 (M+H+)
Synthetic NLL
NLL
Isolated NLL
Synthetic NLL
E
m/z = 357.3103 (M+H+)
Isolated NML
m/z = 357.3103 (M+H+)
Synthetic NML
NML
Isolated NML
Synthetic NML
Time (minutes)
m/z

**Figure S3**. **Comparison of isolated and synthetic fatty acid amides**. Retention times (in extracted ion chromatograms of the indicated m/z, left) and HPLC-HR-MS/MS fragmentation patterns (right) for isolated FAAs (top) and their synthetic standards (bottom) are shown. Predicted fragments are indicated for each molecule. (A) NMP. (B) NAMP. (C) NAPP. (D) NLL. (E) NML.
**Related to Figure 4 and Data S5.**

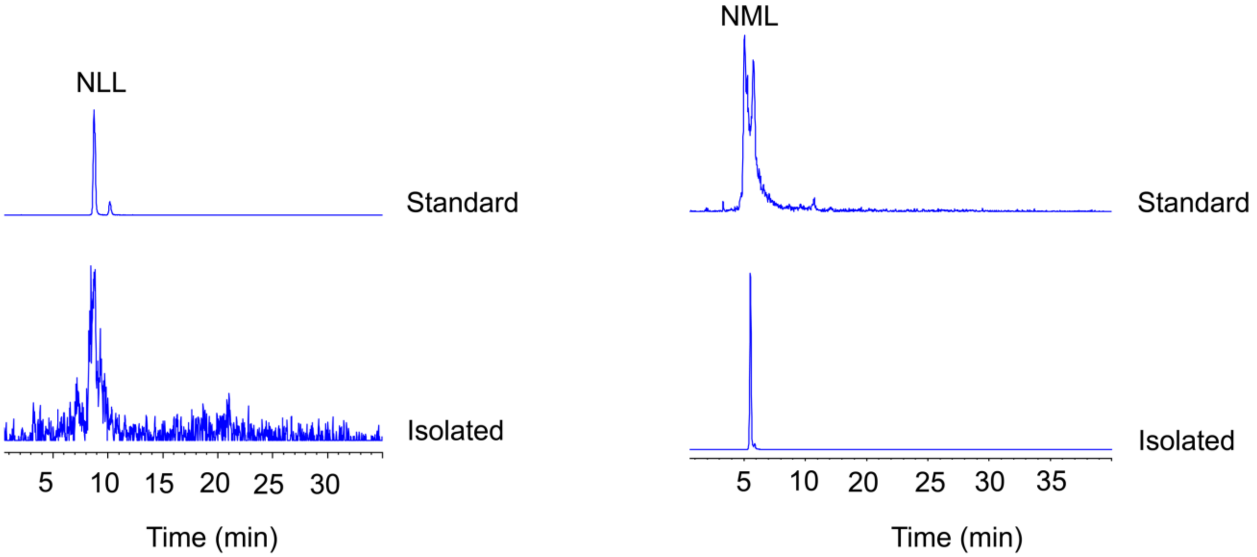


**Figure S4. Comparison of isolated and synthetic NLL and NML.** Retention times (in extracted ion chromatograms) on a chiral column are shown for isolated NLL (left) and NML (right) and their corresponding synthetic standards (top).
**Related to Figure 4 and Data S5.**

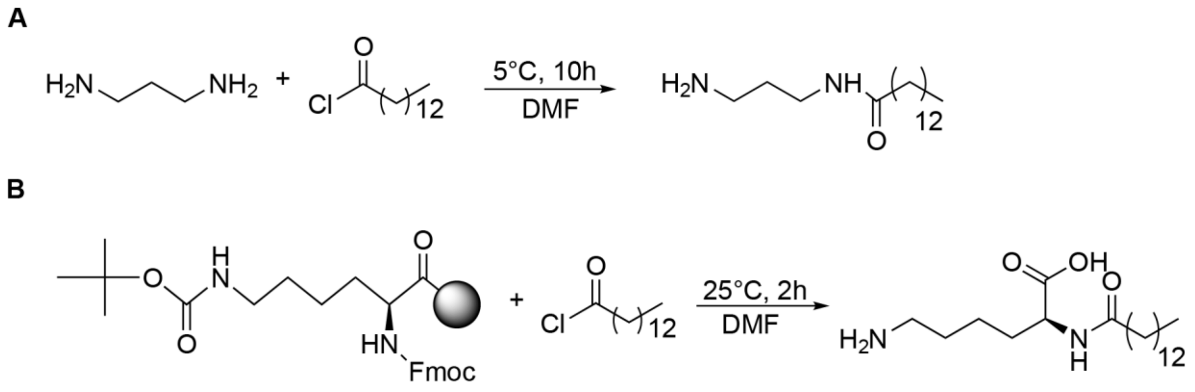


**Figure S5. Synthetic routes for fatty acid amides.** The indicated synthetic schemes were used for the chemical synthesis of: (A) NMP, NAMP, and NAPP. (B) NML. **Related to Figure 4 and Data S5.**

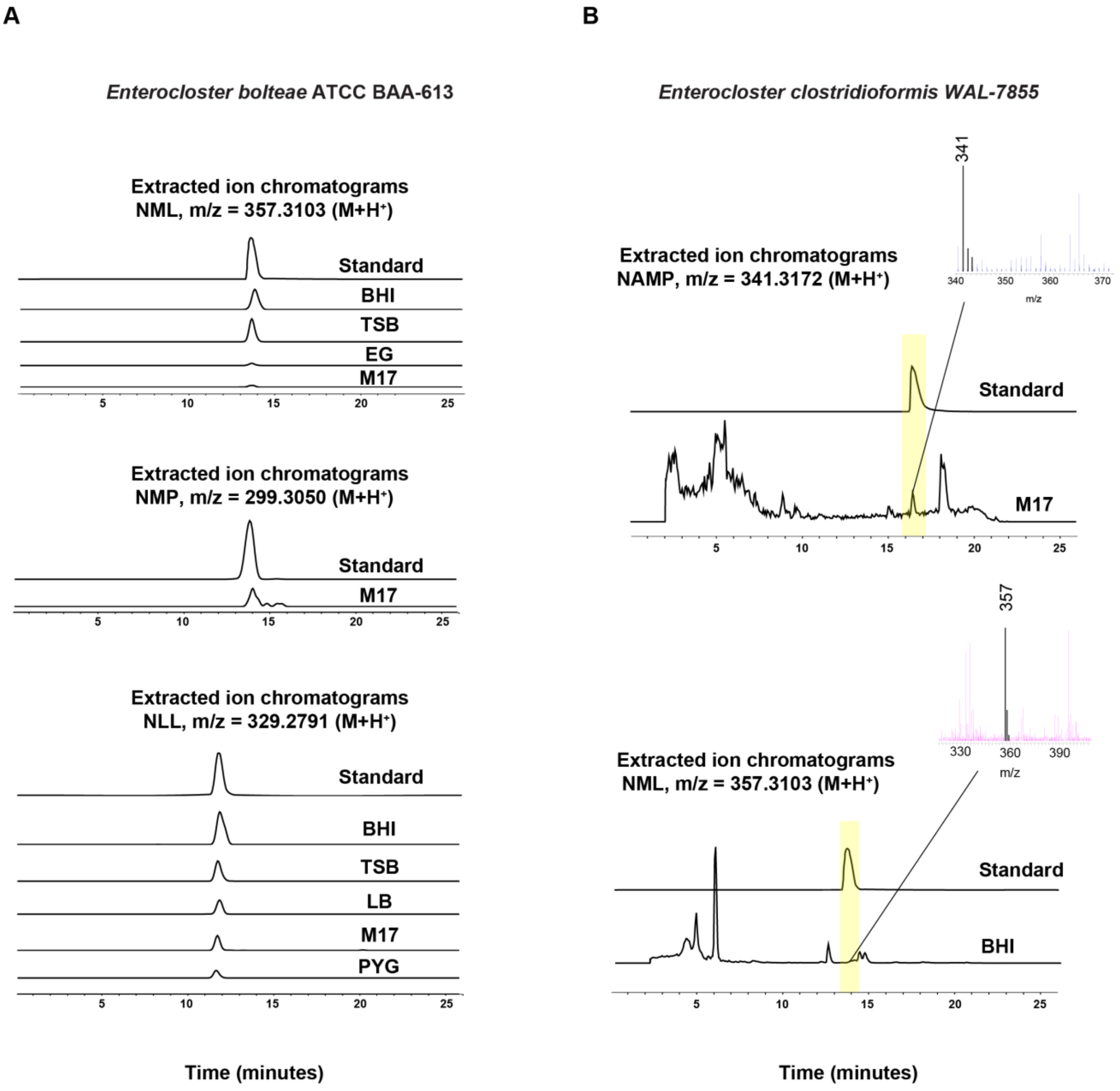


**Figure S6. Production of fatty acid amides by native isolates of *Clostridium* sp.**
(A) Extracted ion chromatograms (HPLC-HR-MS) of NML, NMP, and NLL in cell pellet extracts of *Enterocloster bolteae* ATCC BAA-613 grown in different media are shown. Extracted ion chromatograms of their corresponding synthetic standards are shown on top. (B) Extracted ion chromatograms (HPLC-HR-MS) of NML and NAMP in cell pellet extracts of *Enterocloster clostridioformis* WAL-7855 grown in different media are shown. Extracted ion chromatograms of their corresponding synthetic standards are shown on top.
**Related to Figure 4 and Data S5.**

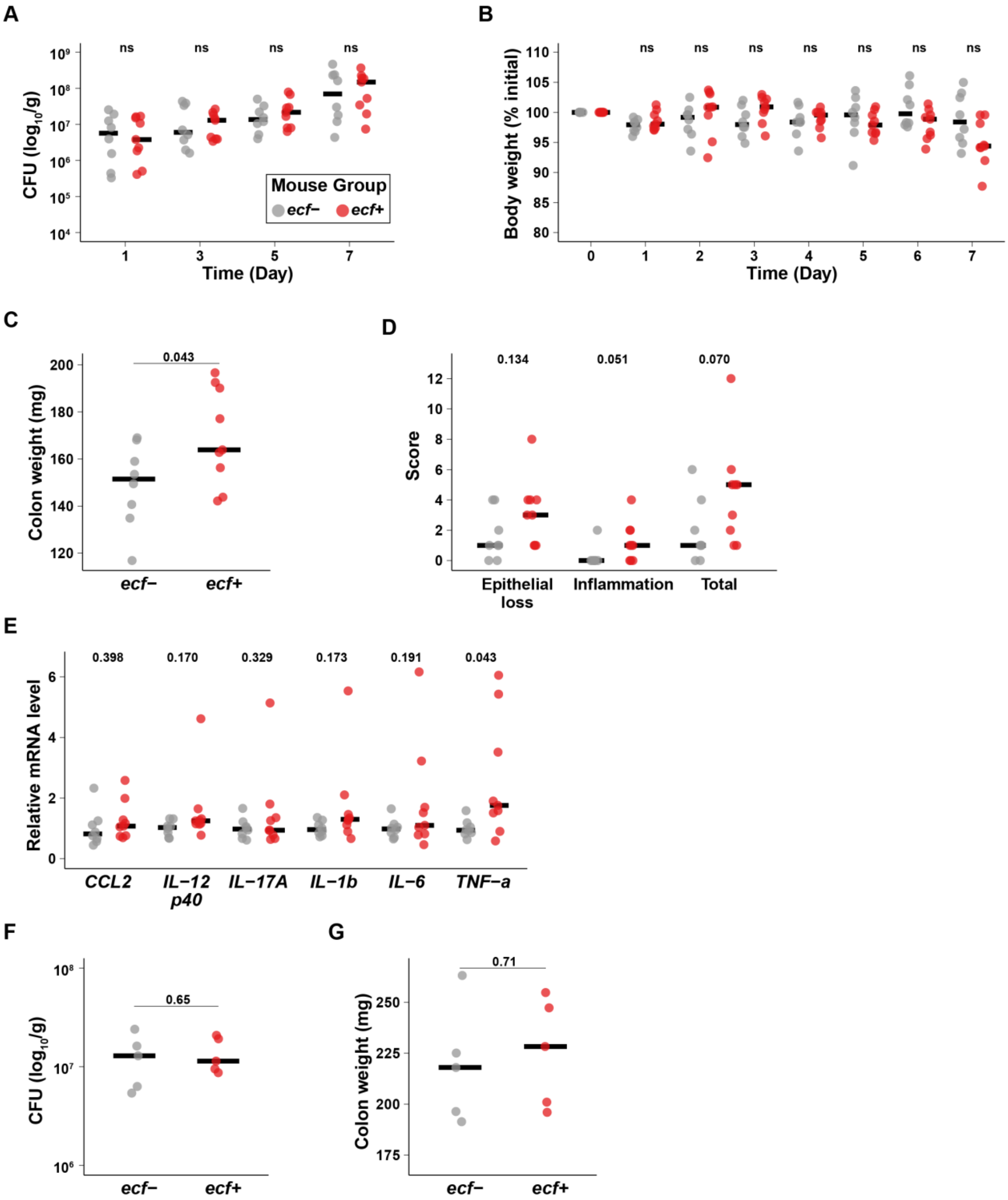
A
ns
ns
ns
ns
CFU (log10/g)
Mouse Group
ecf−
ecf+
Time (Day)
B
Body weight (% initial)
Time (Day)
C
0.043
Colon weight (mg)
ecf−
ecf+
D
0.134
0.051
0.070
Score
Epithelial loss
Inflammation
Total
E
0.398
0.170
0.329
0.173
0.191
0.043
Relative mRNA level
CCL2
IL−12 p40
IL−17A
IL−1b
IL−6
TNF−a
F
0.65
CFU (log10/g)
ecf−
ecf+
G
0.71
Colon weight (mg)
ecf−
ecf+

**Figure S7. Effects of *ebf-ecf*-FAAs in mouse models of colitis. (A–E)** DSS-induced colitis mouse model experiment. *ecf*+ indicates the group of mice colonized with *E. coli* expressing *ecf*, and *ecf*- indicates the group of mice colonized with *E. coli* harboring an empty vector control. Comparison between *ecf*+ and *ecf*- DSS-treated mice in: **(A)** colony forming units (CFUs) of the colonizing *E. coli*, **(B)** body weight, **(C)** colon weight, **(D)** histology scores (epithelial loss, inflammation, and total scores) **(E)** colonic expression of cytokines. **(F–G)** IL-$10^{-/-}$ germ-free mouse model experiment. Comparison between *ecf*+ and *ecf*- IL-$10^{-/-}$ gnotobiotic mice in: **(F)** CFU of the colonizing *E. coli* at week 4, **(G)** colon weight. Data are presented as individual points, and the median is presented as a horizontal line. Data in the DSS-induced colitis model are collected from two independent experiments. Two-sided Student's *t*-test was used to determine statistical significance in all comparisons (ns: not statistically significant).
**Related to Figure 6.**
