## Supplementary material for "A meta-analysis of the gut microbiome in inflammatory bowel disease patients identifies disease-associated small molecules": Data S5

**Table a.** The molecular formulae of FAAs predicted by HR-MS

| Compounds | predicted molecular formula | Calculate m/z $[M+H]^+$ | Observed m/z: $[M+H]^+$ |
|---|---|---|---|
| NMP | $C_{18}H_{39}N_2O$ | 299.3062 | 299.3050 |
| NAMP | $C_{20}H_{41}N_2O_2$ | 341.3168 | 341.3172 |
| NAPeP | $C_{22}H_{43}N_2O_2$ | 367.3325 | 367.3327 |
| NAPP | $C_{22}H_{45}N_2O_2$ | 369.3481 | 369.3480 |
| NLL | $C_{18}H_{37}N_2O_3$ | 329.2804 | 329.2804 |
| NML | $C_{20}H_{41}N_2O_3$ | 357.3117 | 357.3114 |

### 1. NMR spectra of isolated molecules

#### 1.1 NMR spectra of isolated *N*-Acetyl-Myristoyl- Putrescine (NAMP)

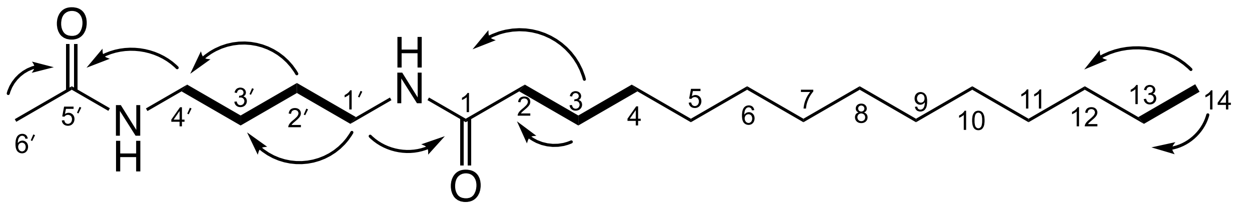


**NAMP**

**Figure a1.** Structure of **NAMP** and key HMBC ⟶ and COSY ▬ correlations.

**Table b.** NMR data for **NAMP** (500 MHz in MeOD)

| Position | δH, mult (*J* in Hz) | δC | HMBC | COSY |
|---|---|---|---|---|
| **1** | | 174.9 | | |
| **2** | 2.19, t | 35.8 | 25.7, 29.0,174.9 | H-3 |
| **3** | 1.62, p | 25.7 | 29.1, 35.8, 174.9 | H-2, H-4 |
| **4** | 1.32 | 29.0 | | H-3 |
| **5** | 1.32 | 29.1 | | |
| **6** | 1.32 | 28.9 | | |
| **7** | 1.32 | 29.0 | | |
| **8** | 1.32 | 29.2 | | |
| **9** | 1.32 | 29.3 | | |
| **10** | 1.32 | 29.4 | | |
| **11** | 1.32 | 29.4 | | |
| **12** | 1.32 | 31.7 | | |

| | | | | |
|---|---|---|---|---|
| **13** | 1.33, br, m | 22.4 | | H-14 |
| **14** | 0.93, t | 13.1 | 22.4, 31.7 | H-13 |
| **1′** | 3.2, m | 38.6 | 26.4,174.9 | H-2′ |
| **2′** | 1.5, p | 26.3 | 38.7 | H-1′, H-3′ |
| **3′** | 1.5, p | 26.4 | 38.6 | H-2′, H-4′ |
| **4′** | 3.2, m | 38.7 | 26.3,171.8 | H-3′ |
| **5′** | | 171.8 | | |
| **6′** | 1.95, s | 21.1 | 171.8 | |

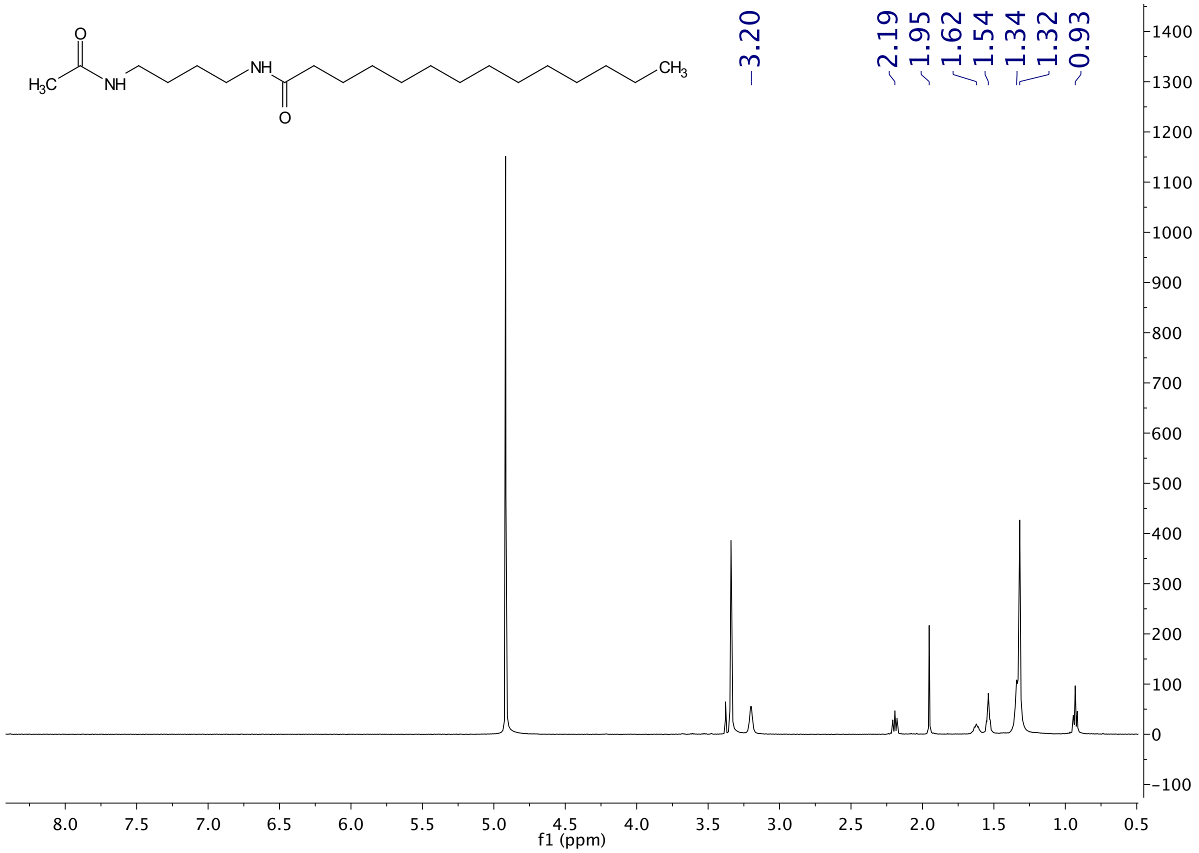


**Figure a2.** $^1$H NMR spectrum of isolated **NAMP** in MeOD

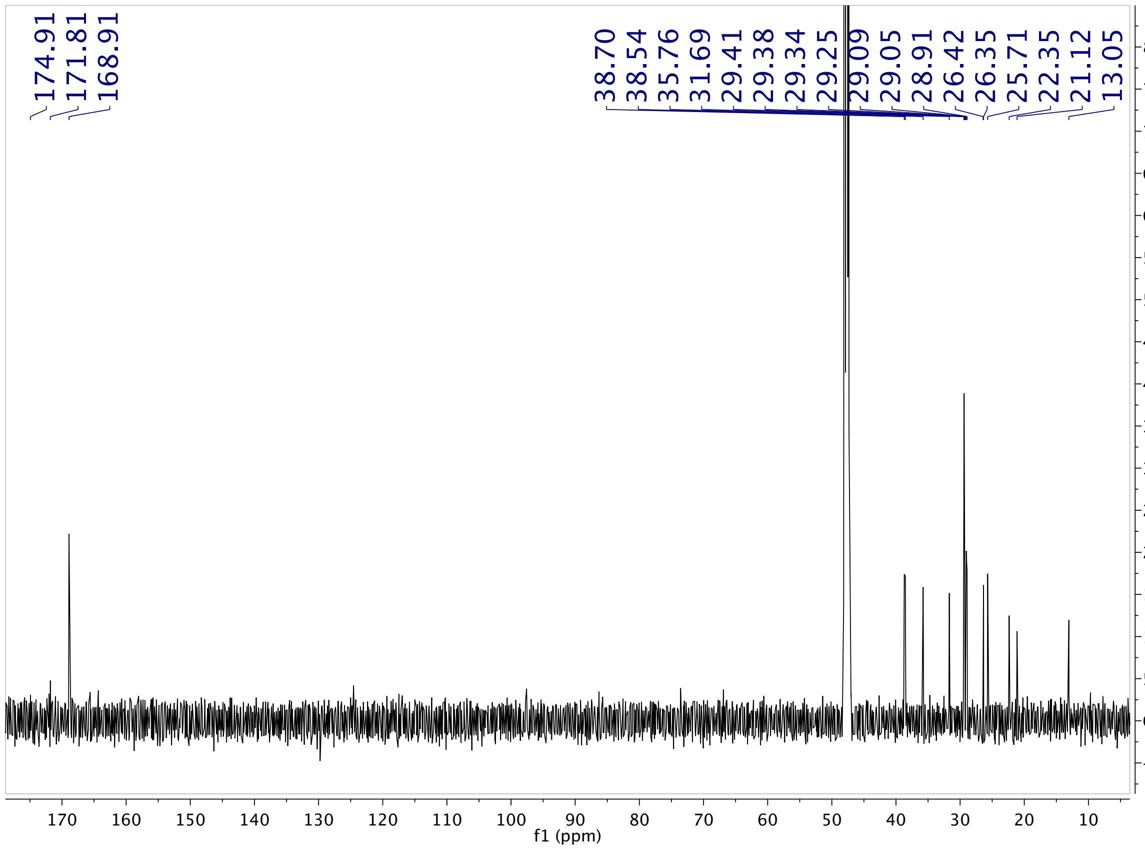


**Figure a3.** $^{13}C$ NMR spectrum of isolated **NAMP** in MeOD

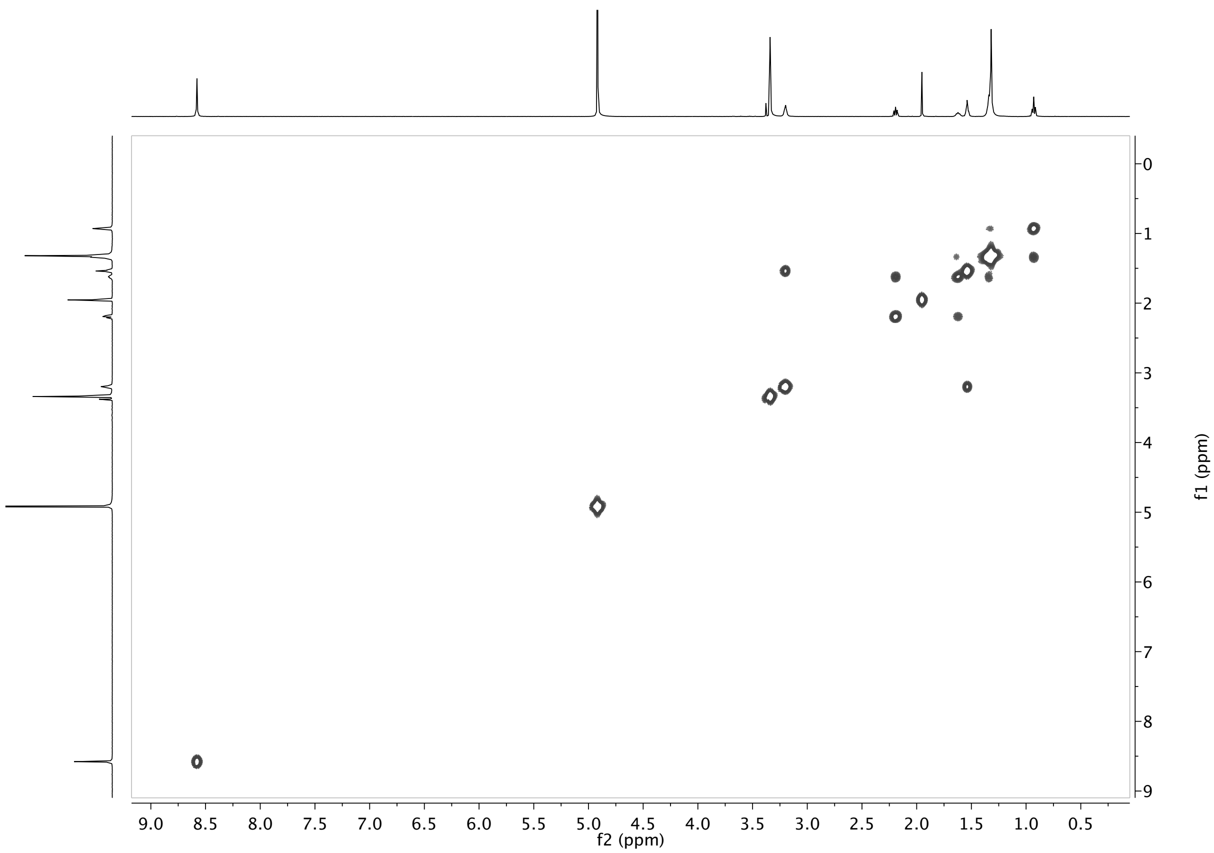


**Figure a4.** COSY spectrum of isolated **NAMP** in MeOD

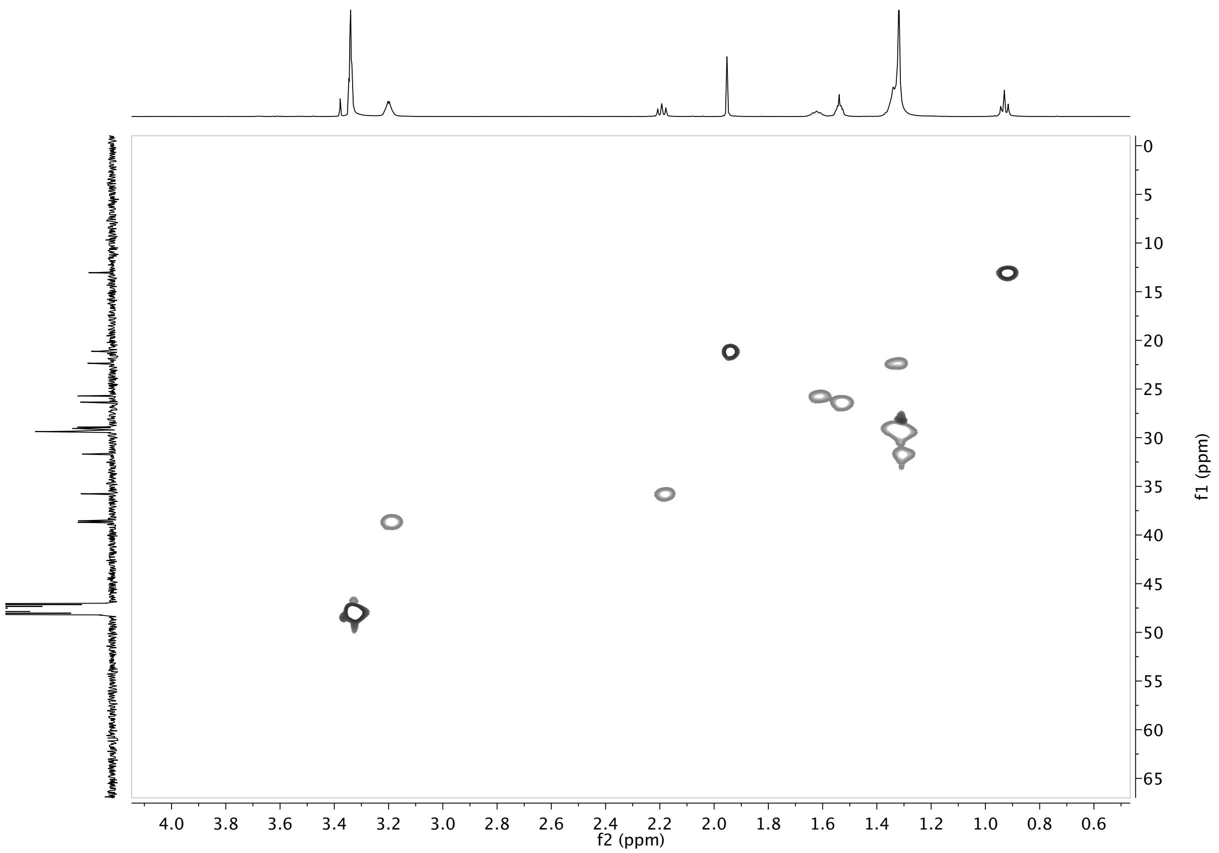


**Figure a5.** HSQC spectrum of isolated **NAMP** in MeOD

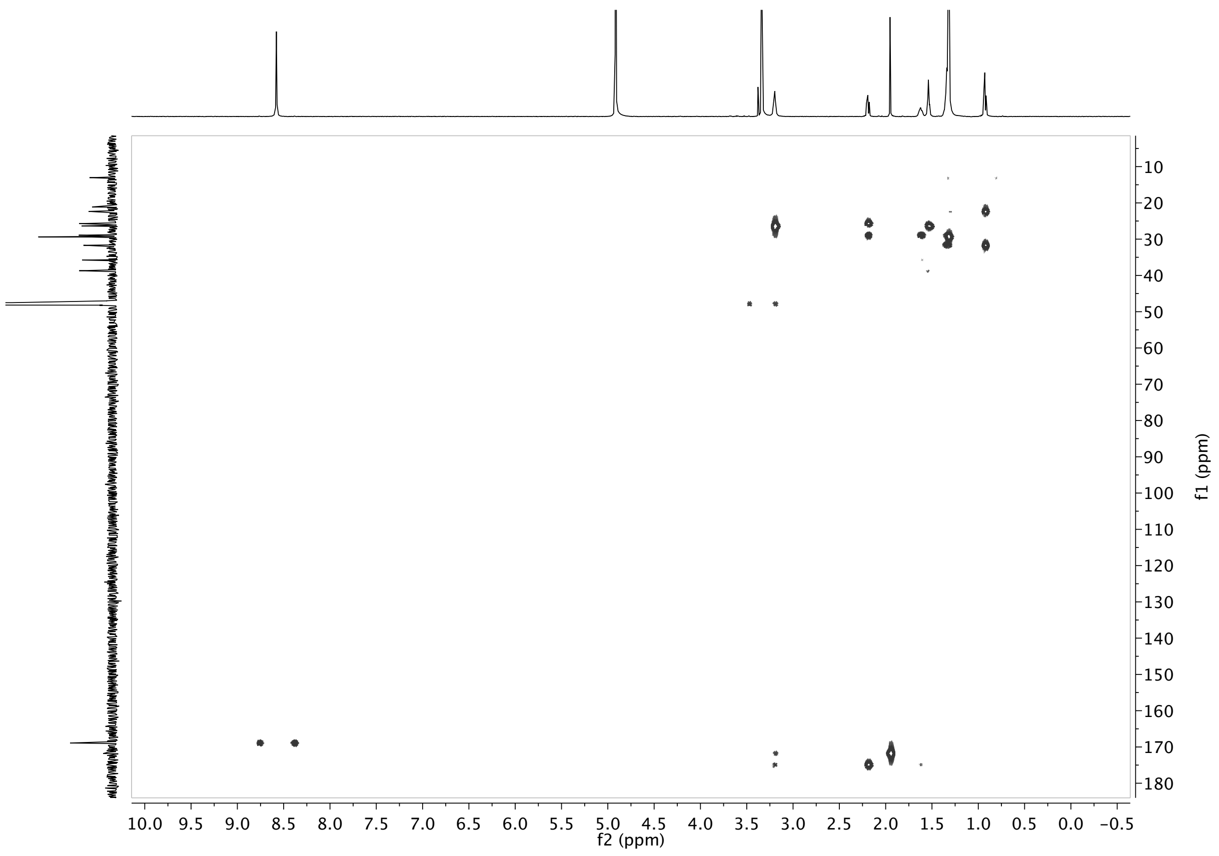


**Figure a6.** HMBC spectrum of isolated **NAMP** in MeOD

### 1.2 NMR spectrum of isolated N-Acetyl-Palmitoyl- Putrescine (NAPP)

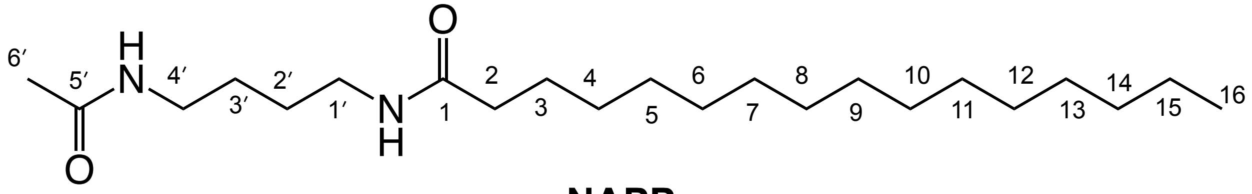


**Figure b1.** Structure of **NAPP**.

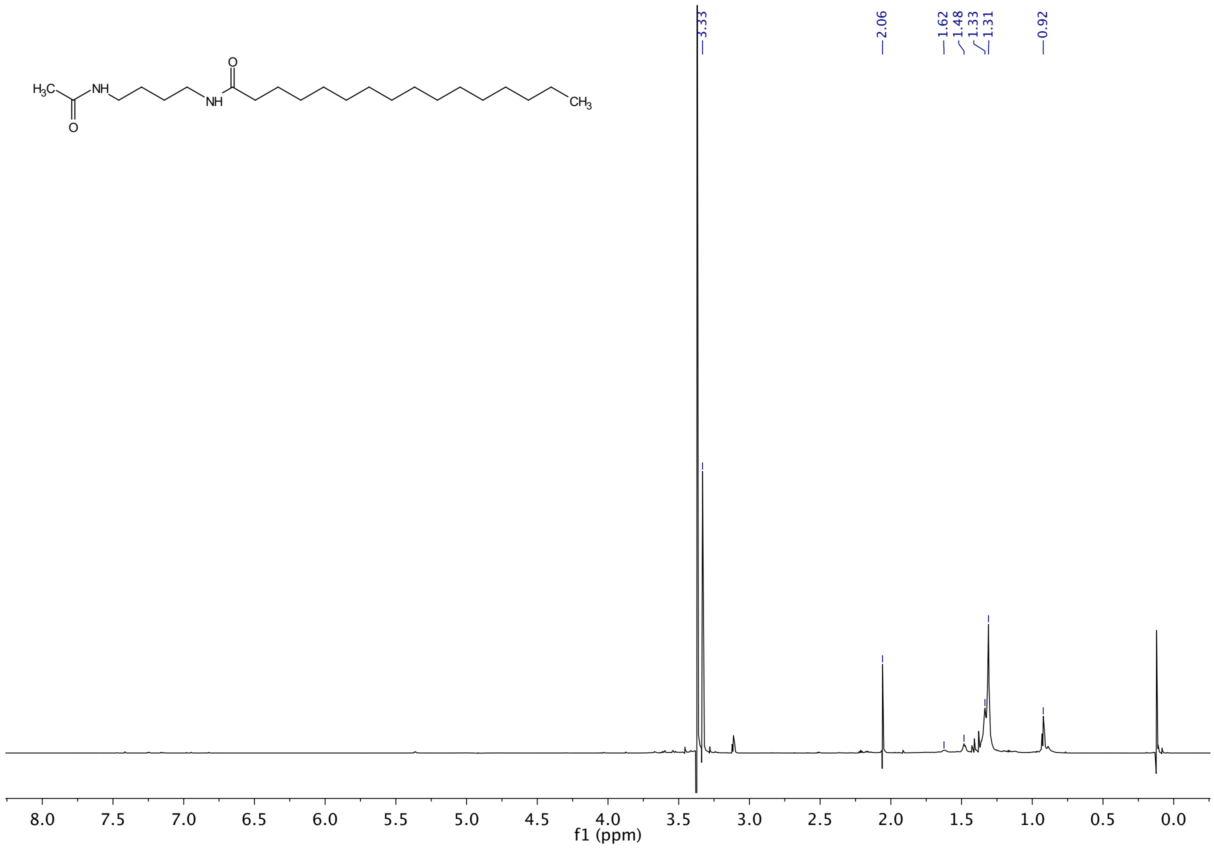


**Figure b2.** ¹H NMR spectrum of isolated **NAPP** in MeOD

**1.3 NMR spectra of isolated *N*-Acetyl-Palmitoleoyl-Putrescine (NAPeP)**

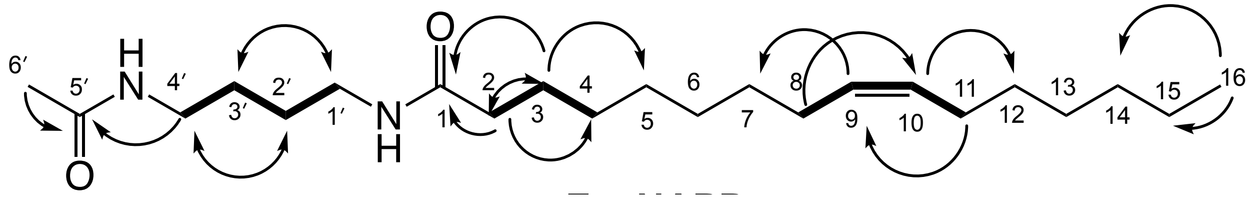


**Figure c1.** Structure of **NAPeP** and key HMBC ⟶ and COSY ▬ correlations.

**Table c.** NMR data for isolated **NAPeP** (800 MHz in MeOD)

| Position | δH, mult (*J* in Hz) | δC | HMBC | COSY |
|---|---|---|---|---|
| **1** | | 174.9 | | |
| **2** | 2.19, t (7.41, 7.59) | 35.8 | 25.7, 29.0,174.9 | H-3 |
| **3** | 1.62, p (5.9, 7.28, 7.0, 6.25) | 25.7 | 29.1, 35.8, 174.9 | H-2, H-4 |
| **4** | 1.32,m | 29.0 | | H-3 |
| **5** | 1.32,m | 29.1 | | |
| **6** | 1.32,m | 29.4 | | |
| **7** | 1.32,m | 29.0 | | |
| **8** | 2.07,m | 26.7 | 129.4, 29.4 | H-9 |
| **9** | 5.37,m | 129.4 | 26.7 | H-10,H-8 |
| **10** | 5.37,m | 129.4 | 26.7 | H-11, H-9 |
| **11** | 2.07,m | 26.7 | 129.4, 29.5 | H-10 |
| **12** | 1.32,m | 28.9 | | |
| **13** | 1.32,m | 29.5 | | |
| **14** | 1.32,m | 31.5 | | |
| **15** | 1.34, br, m | 22.4 | | H-14 |
| **16** | 0.93, t (6.64, 7.07) | 13.0 | 22.4, 31.5 | H-13 |
| **1′** | 3.2, m | 38.6 | 26.4,174.9 | H-2′ |
| **2′** | 1.54, p (3.36, 3.32, 3.42, 3.27) | 26.3 | 38.7 | H-1′, H-3′ |
| **3′** | 1.54, p (3.36, 3.32, 3.42, 3.27) | 26.4 | 38.6 | H-2′, H-4′ |
| **4′** | 3.2, m | 38.7 | 26.3,171.8 | H-3′ |
| **5′** | | 171.8 | | |
| **6′** | 1.95, s | 21.1 | 171.8 | |

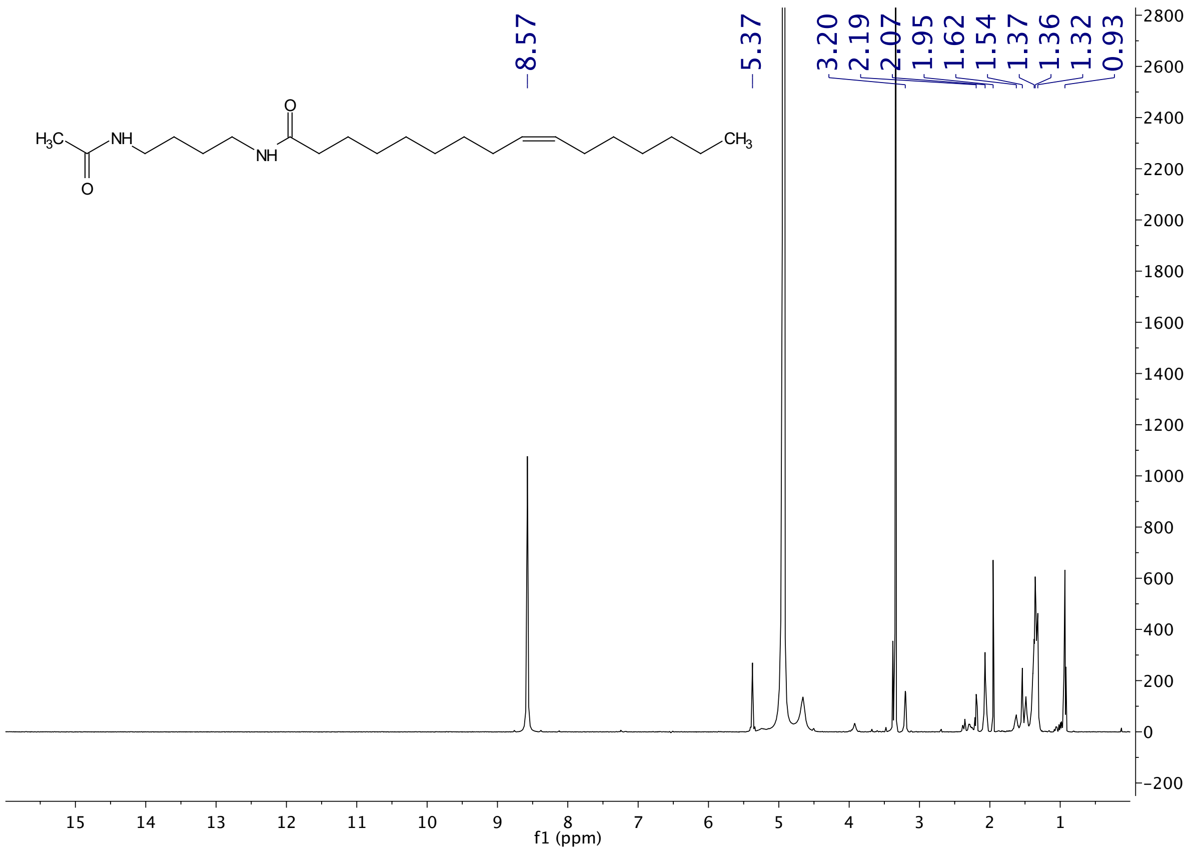


**Figure c2.** $^{1}H$ NMR spectrum of isolated **NAPeP** in MeOD

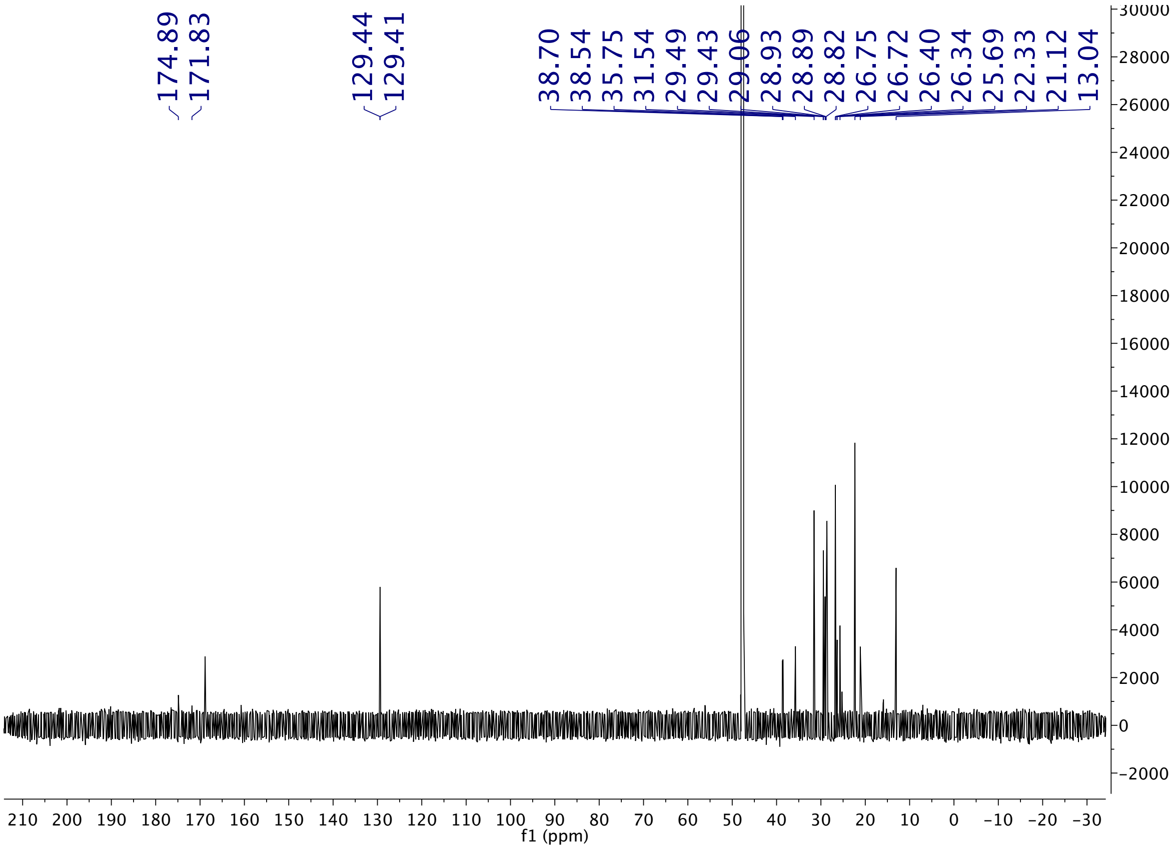


**Figure c3.** $^{13}C$ NMR spectrum of isolated **NAPeP** in MeOD

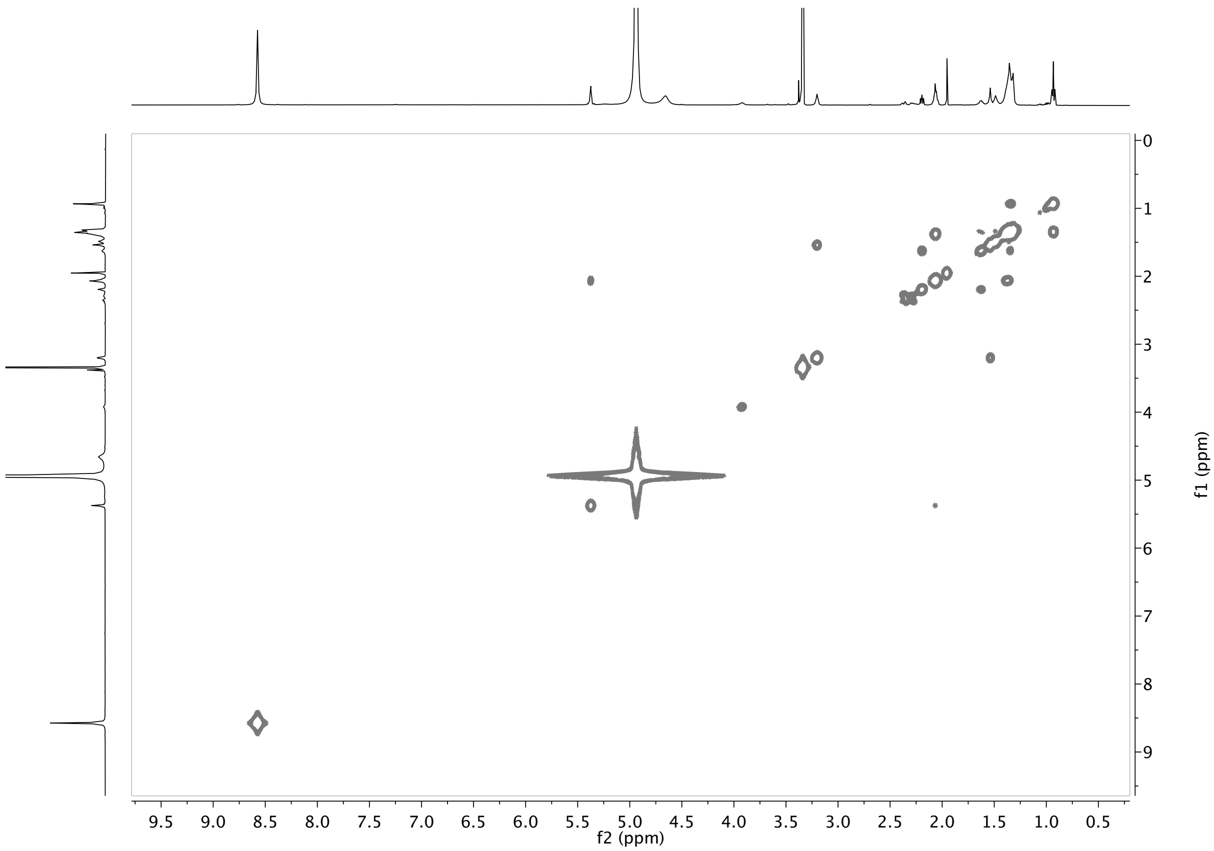


**Figure c4.** COSY NMR spectrum of isolated **NAPeP** in MeOD

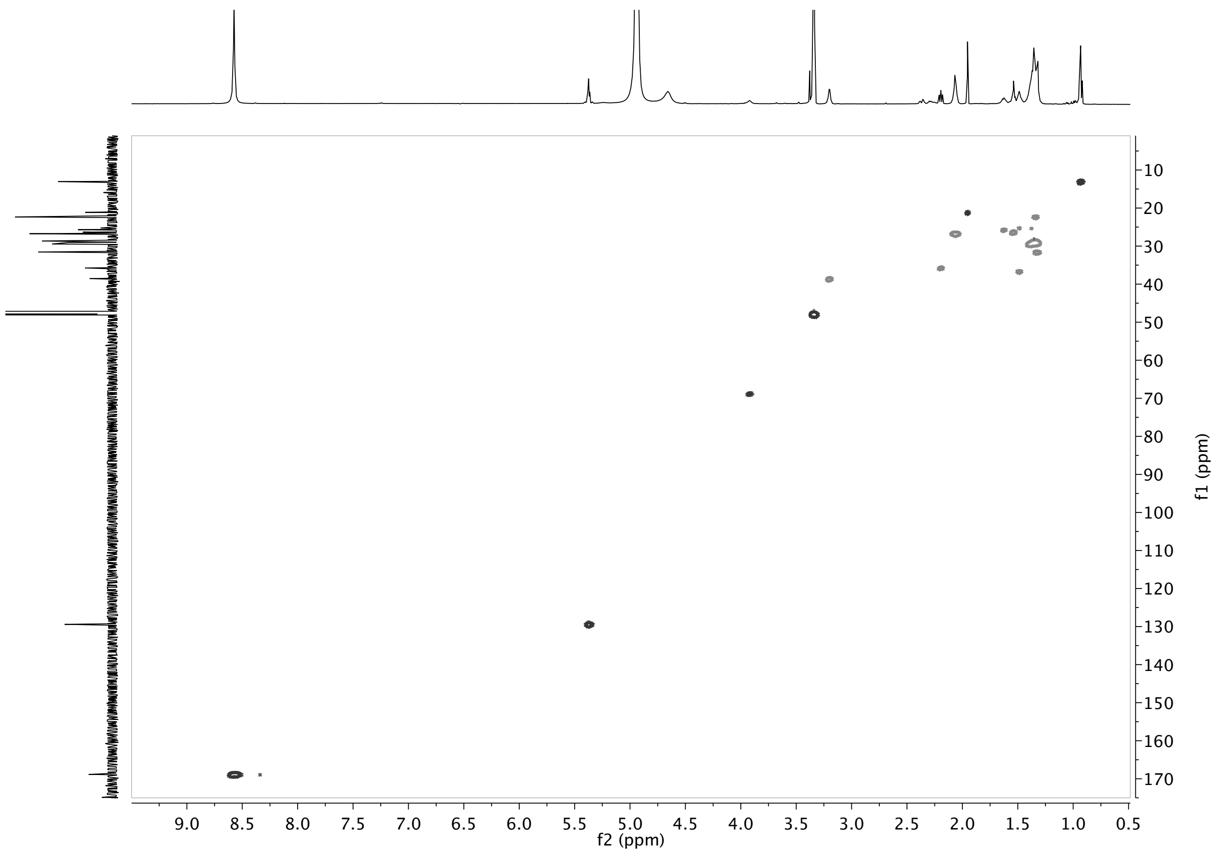


**Figure c5.** HSQC NMR spectrum of isolated **NAPeP** in MeOD

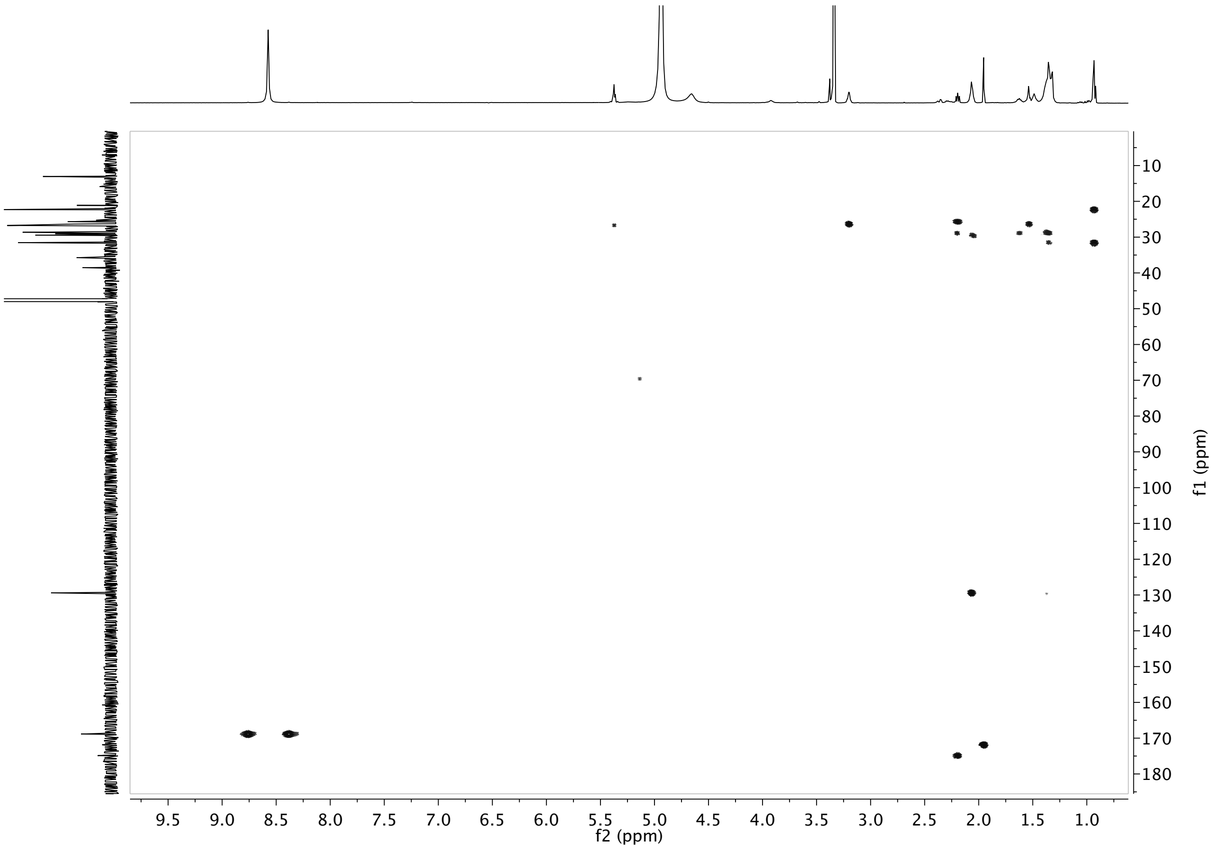


**Figure c6.** HMBC NMR spectrum of isolated **NAPeP** in MeOD

### 2. NMR spectra of synthetic molecules

#### 2.1 NMR spectra of synthetic *N*-Myristoyl-Putrescine (NMP)

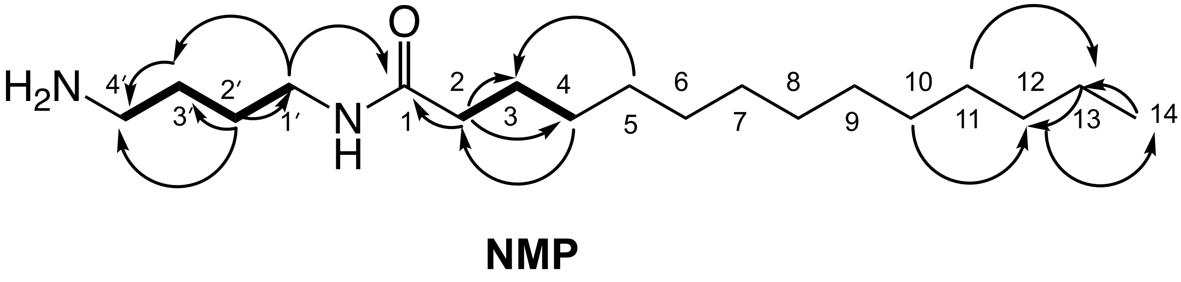


**Figure d1.** Structure of synthetic **NMP** and key HMBC ⟶ and COSY ▬ correlations.

**Table d.** NMR data for synthetic **NMP** (500 MHz in MeOD)

| Position | δH, mult (*J* in Hz) | δC | HMBC | COSY |
|---|---|---|---|---|
| **1** | | 175.0 | | |
| **2** | 2.20, t (7.51,7.51) | 35.8 | 25.7, 29.0,175.0 | H-3 |
| **3** | 1.61, m | 25.7 | 29.0, 35.8, 174.0 | H-2, H-4 |
| **4** | 1.32 | 29.0 | 35.7 | H-3 |
| **5** | 1.32 | 29.1 | 25.7 | |
| **6** | 1.32 | 29.1 | | |
| **7** | 1.32 | 29.3 | | |
| **8** | 1.32 | 29.4 | | |
| **9** | 1.32 | 29.4 | | |
| **10** | 1.32 | 29.4 | 31.7 | |
| **11** | 1.32 | 29.4 | 22.4 | |
| **12** | 1.32 | 31.7 | 13.1 | |
| **13** | 1.33, br, m | 22.4 | 31.7 | H-14 |
| **14** | 0.93, t (6.77, 7.11) | 13.1 | 22.4, 31.7 | H-13 |
| **1′** | 3.23, t (6.93,6.77) | 38.1 | 24.6,175.0 | H-2′ |
| **2′** | 1.61, br, m | 26.1 | 38.9,24.6,38.1 | H-1′, H-3′ |
| **3′** | 1.69, m | 24.6 | 38.9,26.1,38.1 | H-2′, H-4′ |
| **4′** | 2.96, t (7.33,7.64) | 38.9 | 26.1,24.6 | H-3′ |

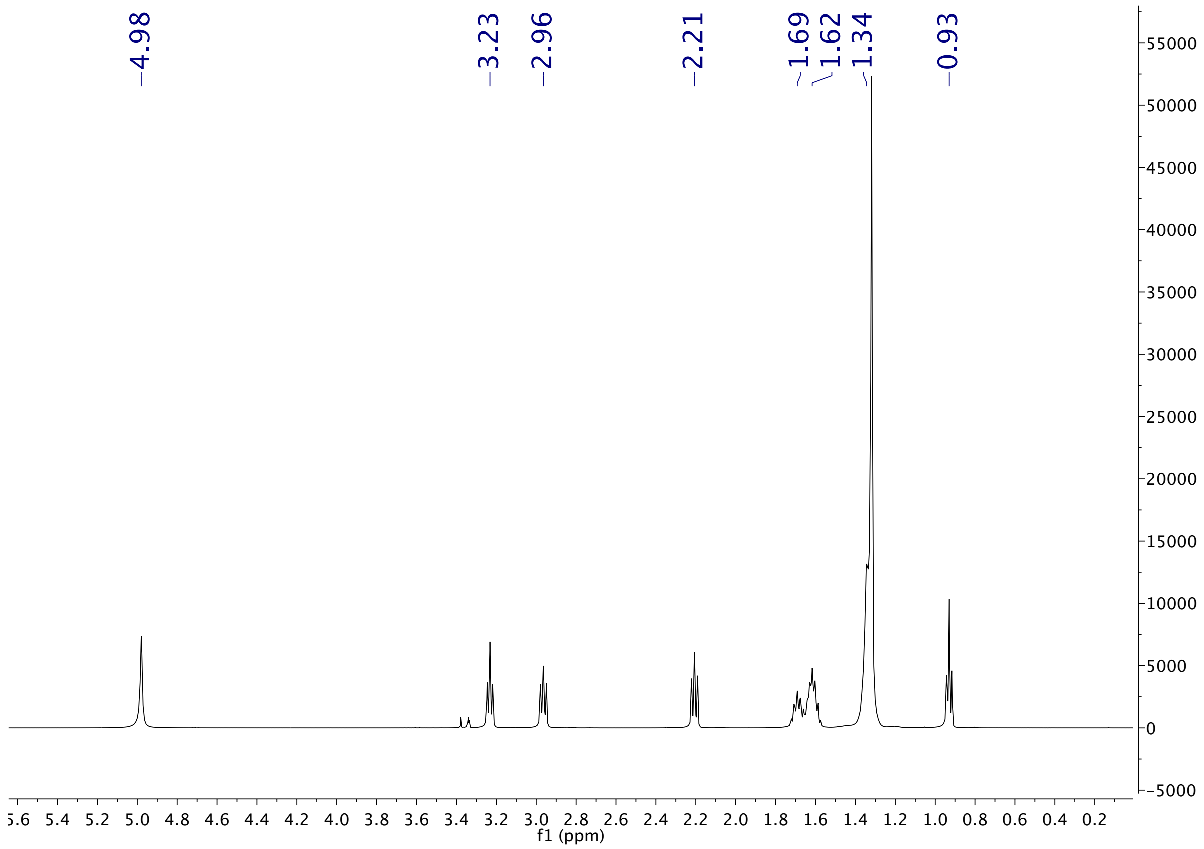


**Figure d2.** $^{1}H$ NMR spectrum of synthetic **NMP** in MeOD

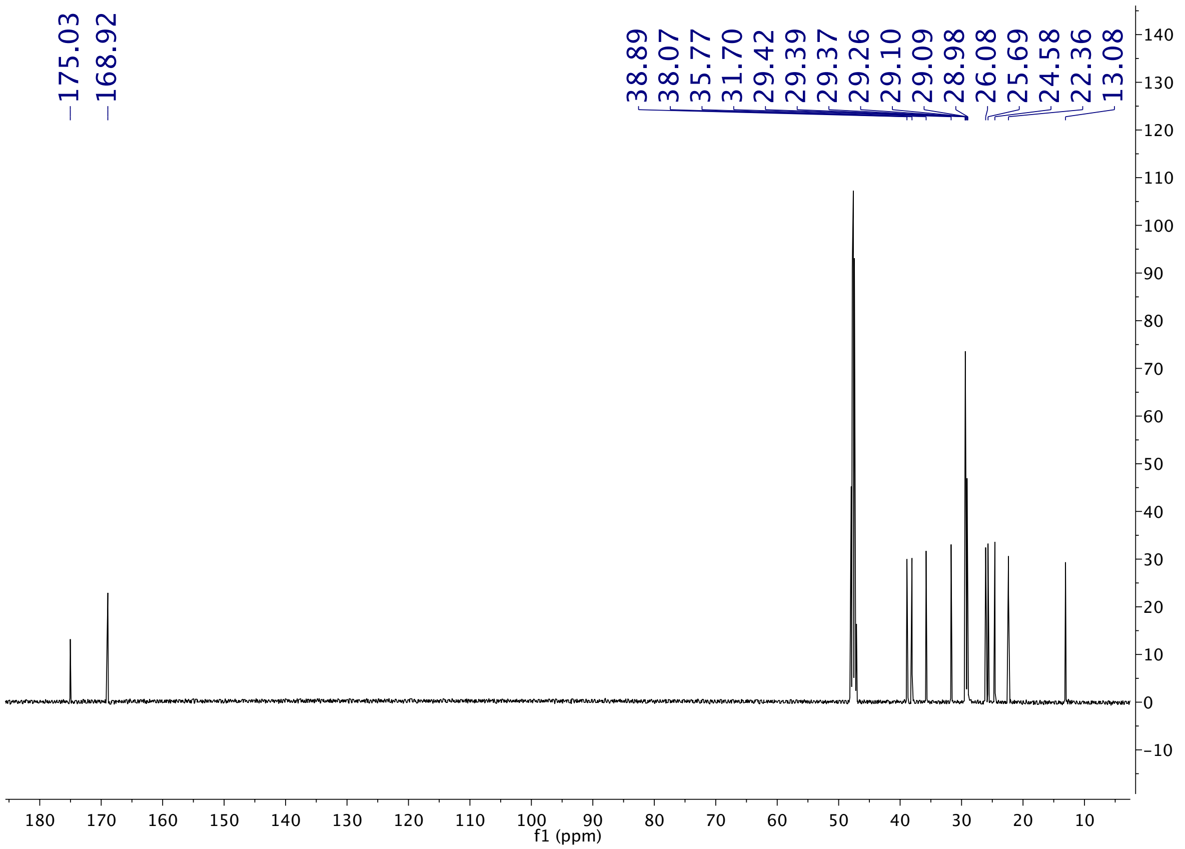


**Figure d3.** $^{13}C$ NMR spectrum of synthetic **NMP** in MeOD

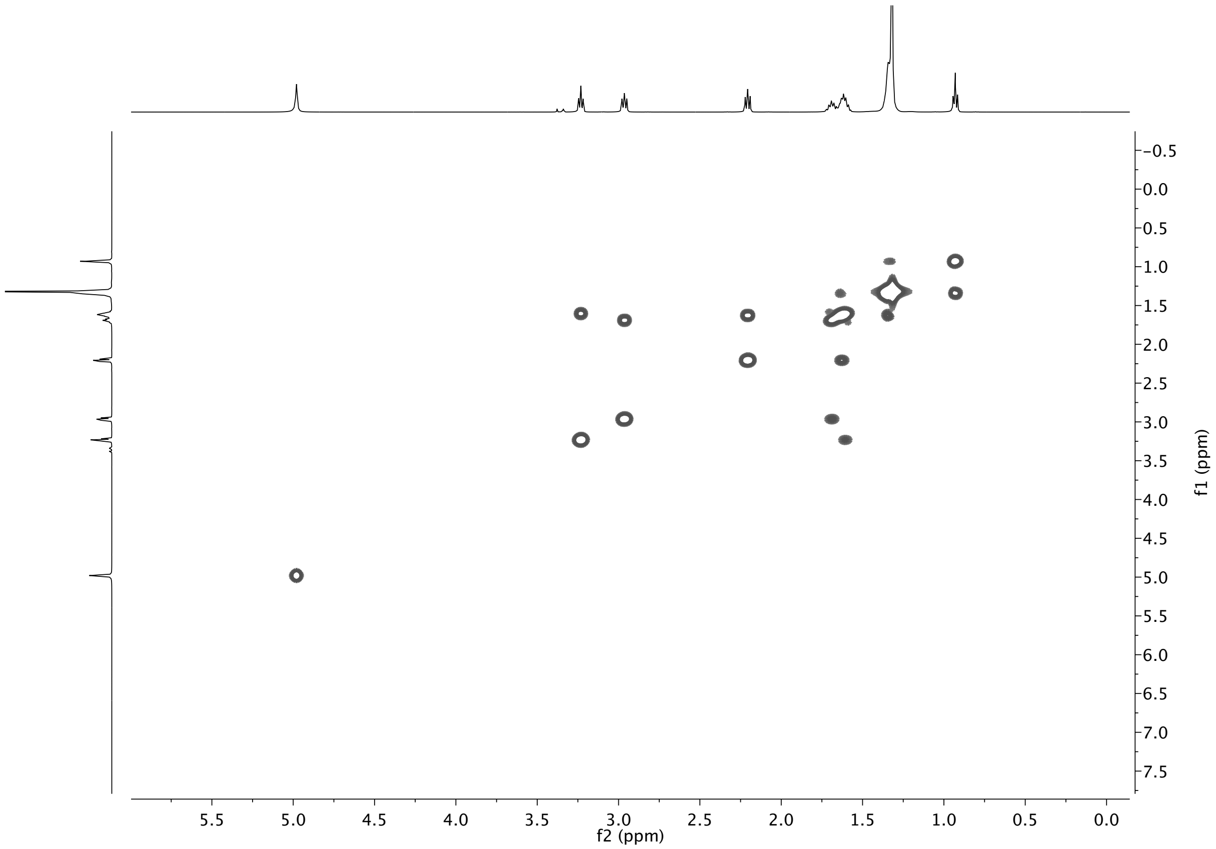


**Figure d4.** COSY spectrum of synthetic **NMP** in MeOD

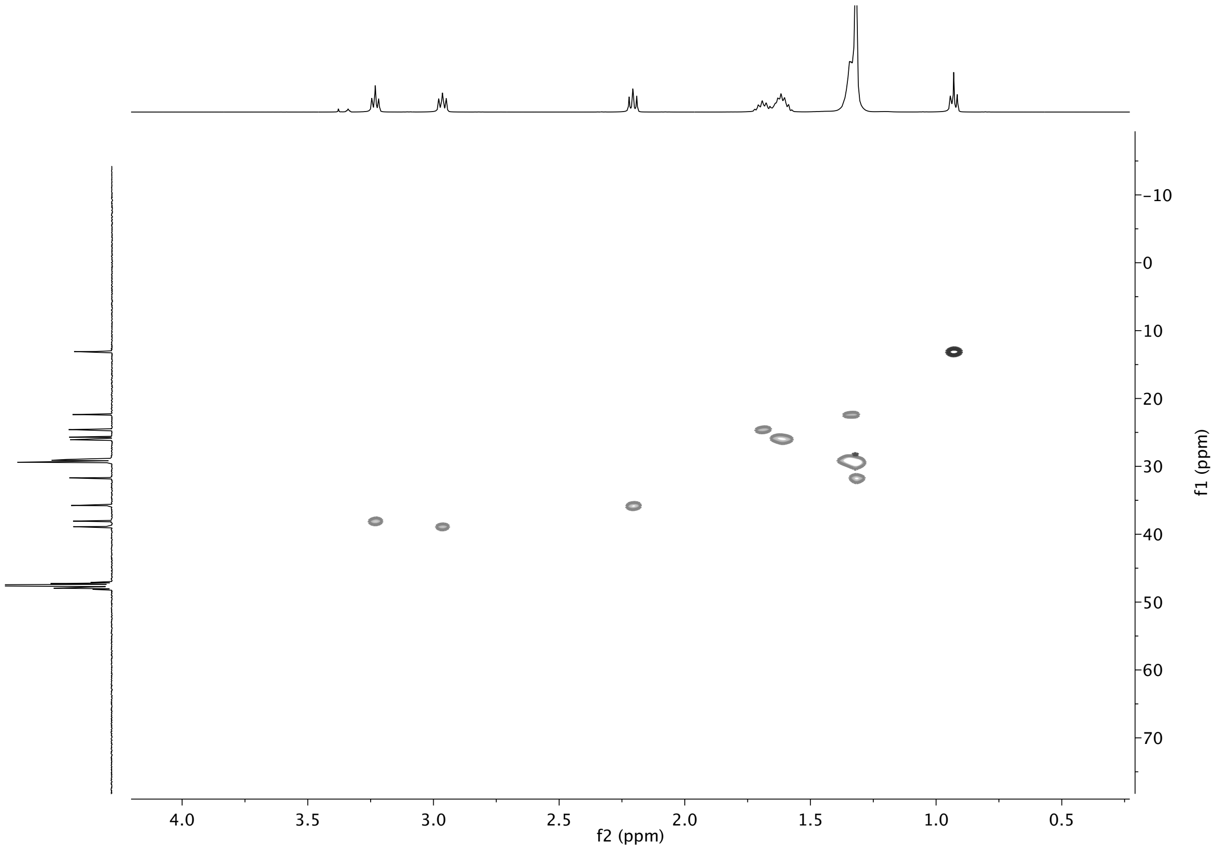


**Figure d5.** HSQC spectrum of synthetic **NMP** in MeOD

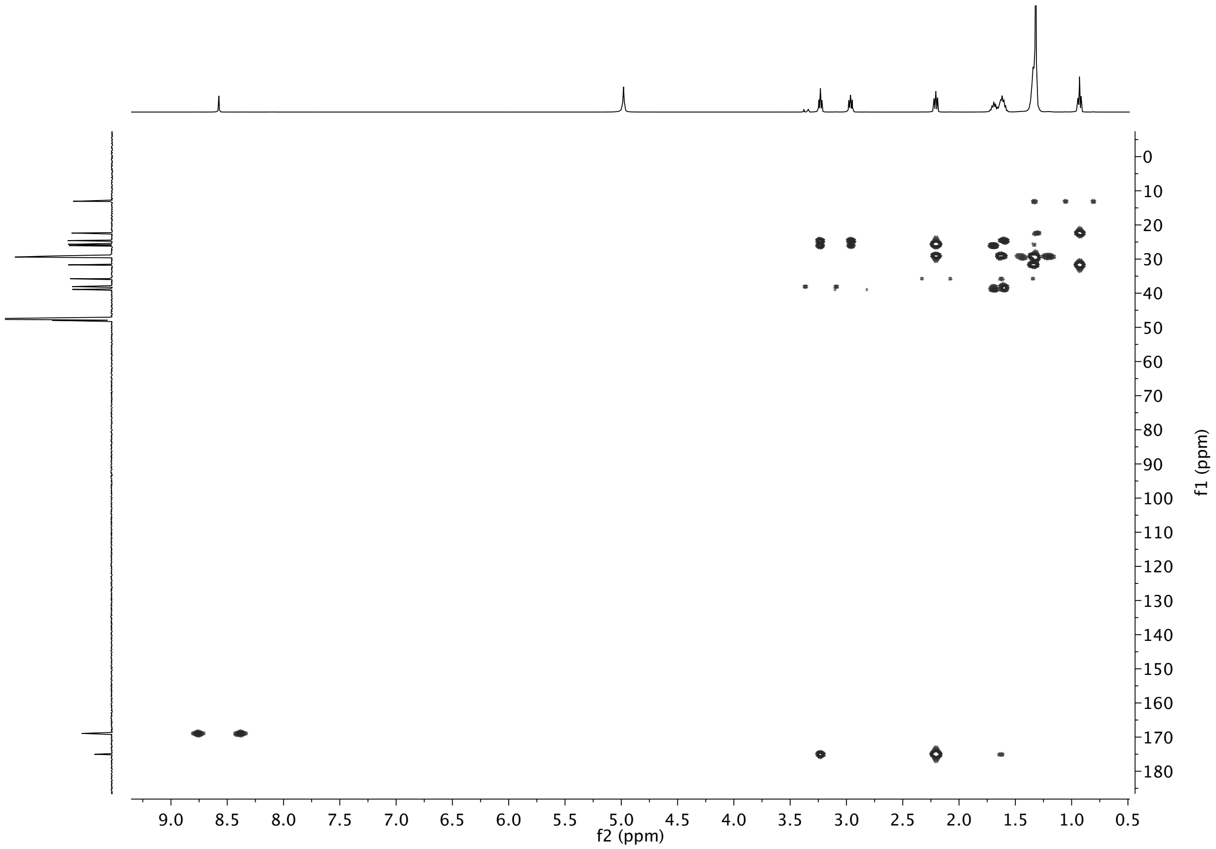


**Figure d6.** HMBC spectrum of synthetic **NMP** in MeOD

**2.2 NMR spectra of synthetic *N*-Acetyl-Palmitoyl-Putrescine (NAPP)**

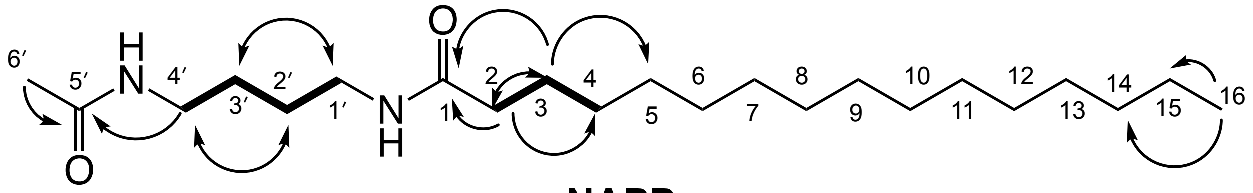


**NAPP**

**Figure e1.** Structure of synthetic **NAPP** and key HMBC ⟶ and COSY ▬ correlations

**Table e.** NMR data for synthetic **NAPP** (500 MHz in MeOH)

| Position | δH, mult (*J* in Hz) | δC | HMBC | COSY |
|---|---|---|---|---|
| **1** | | 174.9 | | |
| **2** | 2.19, t (7.41, 7.59) | 35.8 | 25.7, 29.0,174.9 | H-3 |
| **3** | 1.62, p (5.9, 7.28, 7.0, 6.25) | 25.7 | 29.1, 35.8, 174.9 | H-2, H-4 |
| **4** | 1.32 | 29.0 | | H-3 |
| **5** | 1.32 | 29.1 | | |
| **6** | 1.32 | 28.9 | | |
| **7** | 1.32 | 29.0 | | |
| **8** | 1.32 | 29.3 | | |
| **9** | 1.32 | 29.4 | | |
| **10** | 1.32 | 29.4 | | |
| **11** | 1.32 | 29.4 | | |
| **12** | 1.32 | 29.4 | | |
| **13** | 1.32 | 29.4 | | |
| **14** | 1.32 | 31.7 | | |
| **15** | 1.33, br, m | 22.4 | | H-14 |
| **16** | 0.93, t (6.64, 7.07) | 13.1 | 22.4, 31.7 | H-13 |
| **1′** | 3.2, m | 38.6 | 26.4,174.9 | H-2′ |
| **2′** | 1.5, p (3.36, 3.32, 3.42, 3.27) | 26.3 | 38.7 | H-1′, H-3′ |
| **3′** | 1.5, p (3.36, 3.32, 3.42, 3.27) | 26.4 | 38.6 | H-2′, H-4′ |

| | | | | |
|---|---|---|---|---|
| **4′** | 3.2, m | 38.7 | 26.3,171.8 | H-3′ |
| **5′** | | 171.8 | | |
| **6′** | 1.95, s | 21.1 | 171.8 | |
| **N-H** | 5.84 in CDCl3 | | | |
| **N-H** | 5.69 in CDCl3 | | | |

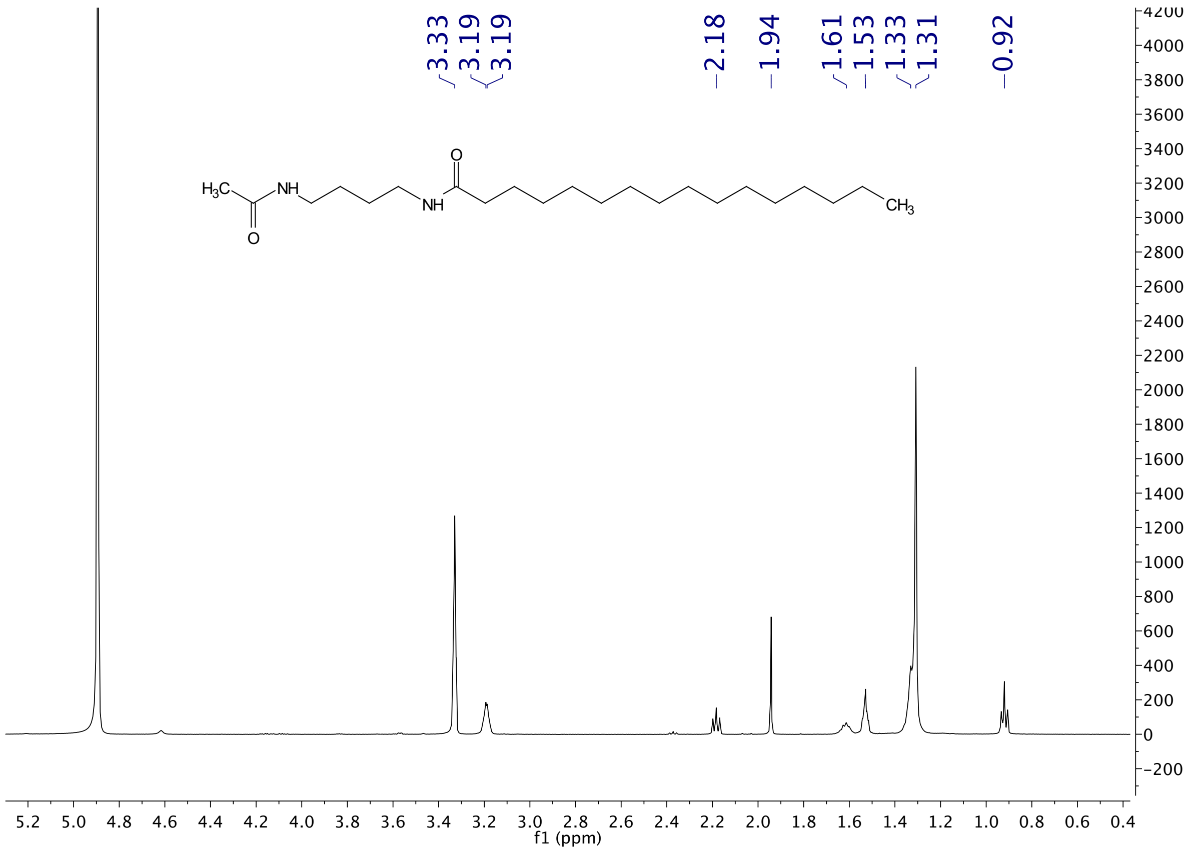


**Figure e2.** [1]H NMR spectrum of synthetic **NAPP** in MeOH

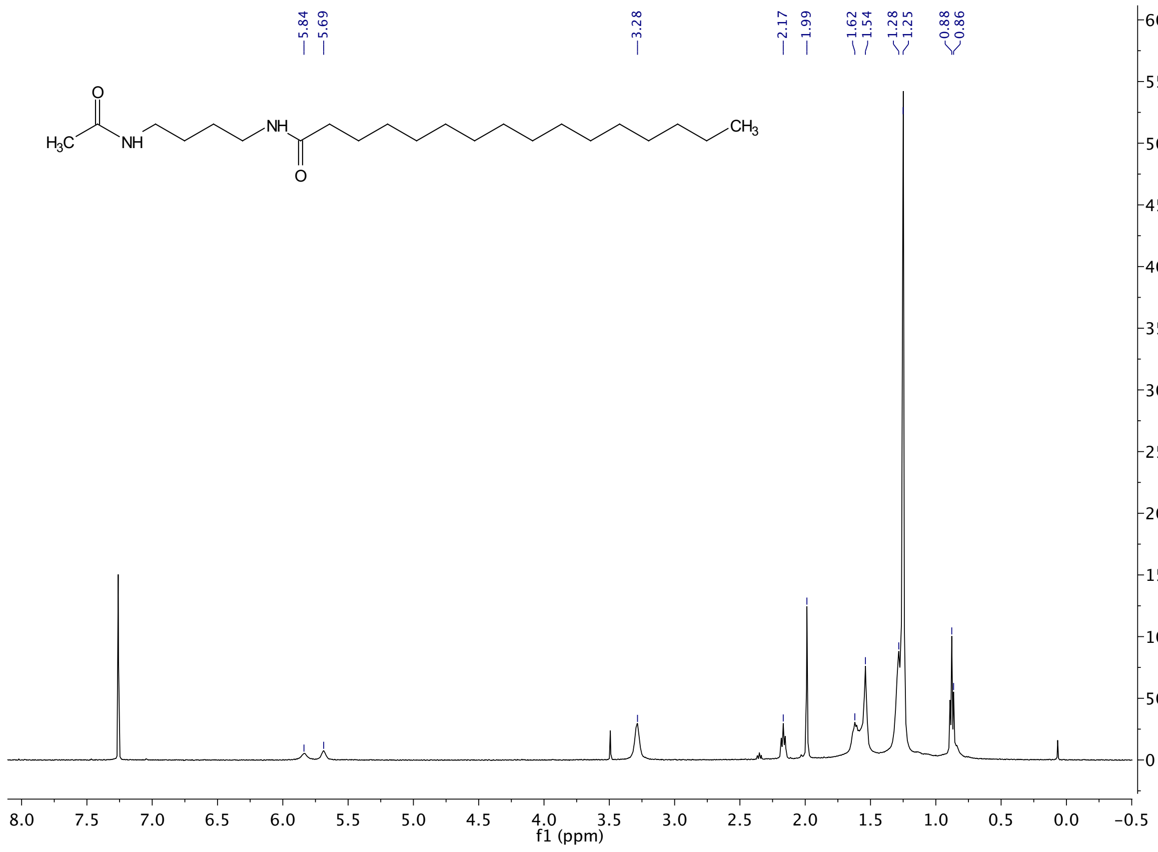


**Figure e3.** [1]H NMR spectrum of synthetic **NAPP** in $CDCl_3$

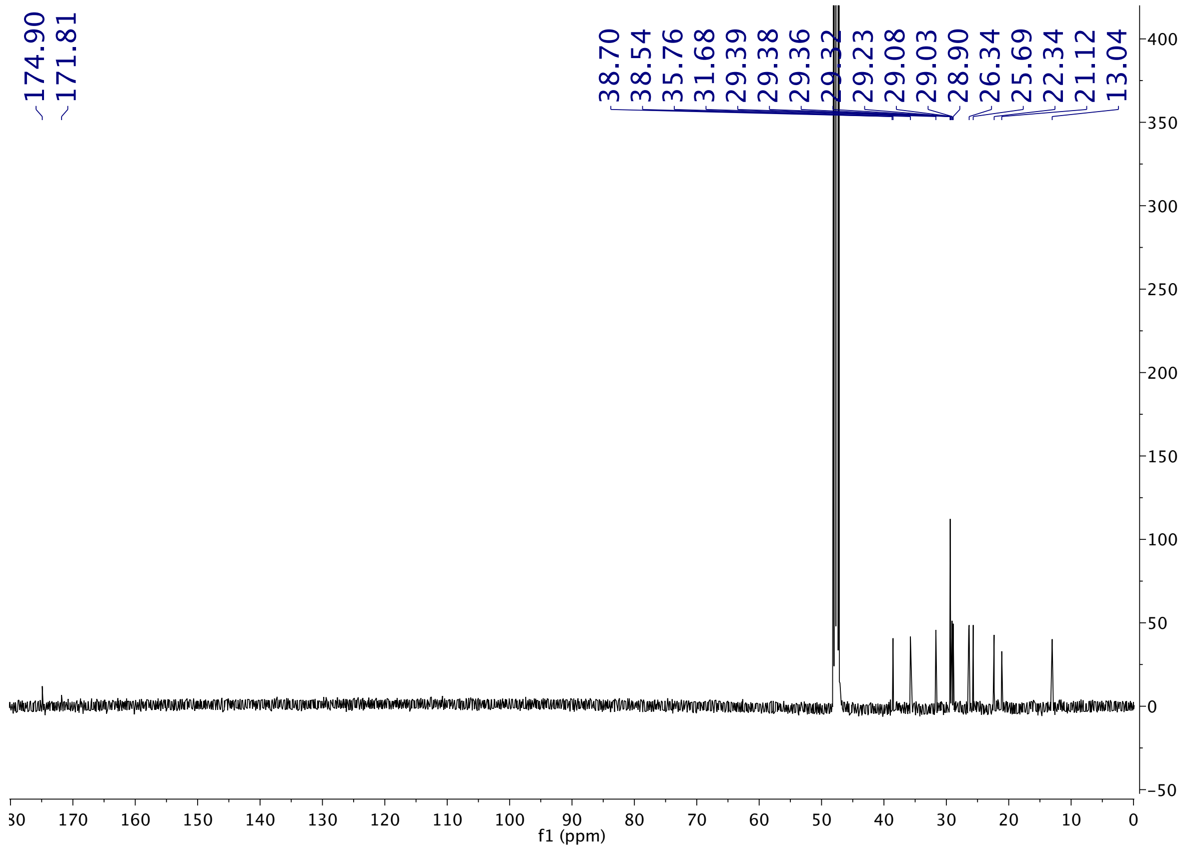


**Figure e4.** $^{13}C$ NMR spectrum of synthetic **NAPP** in MeOD

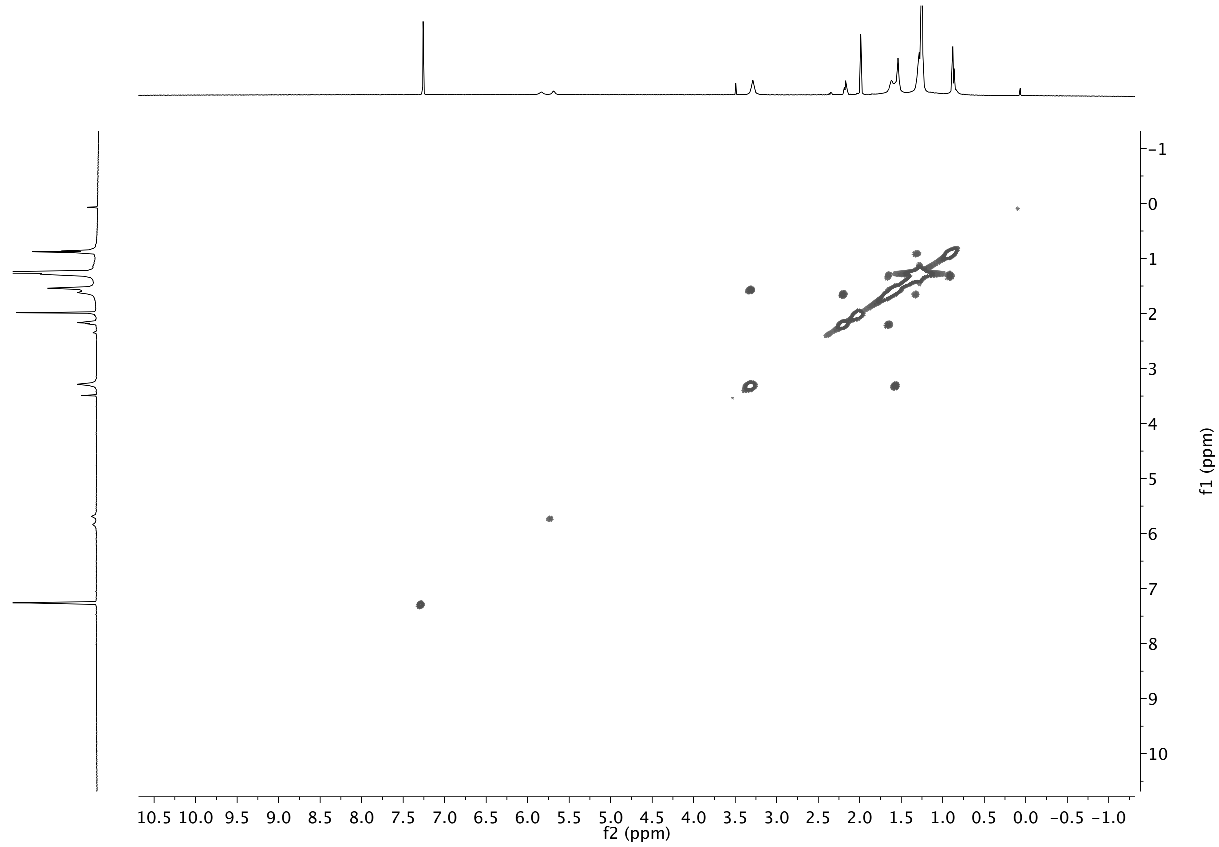


**Figure e5.** COSY spectrum of synthetic **NAPP** in $CDCl_3$

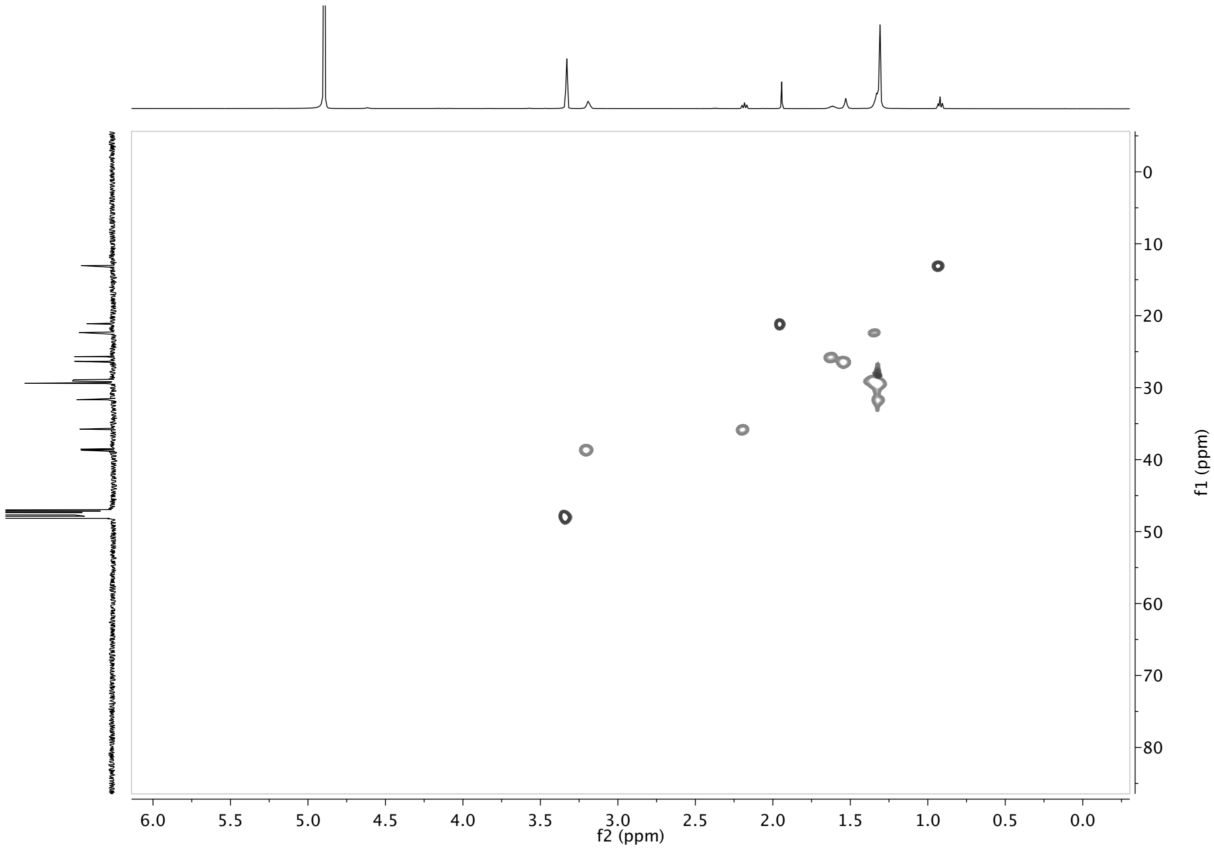


**Figure e6.** HSQC spectrum of synthetic **NAPP** in MeOH

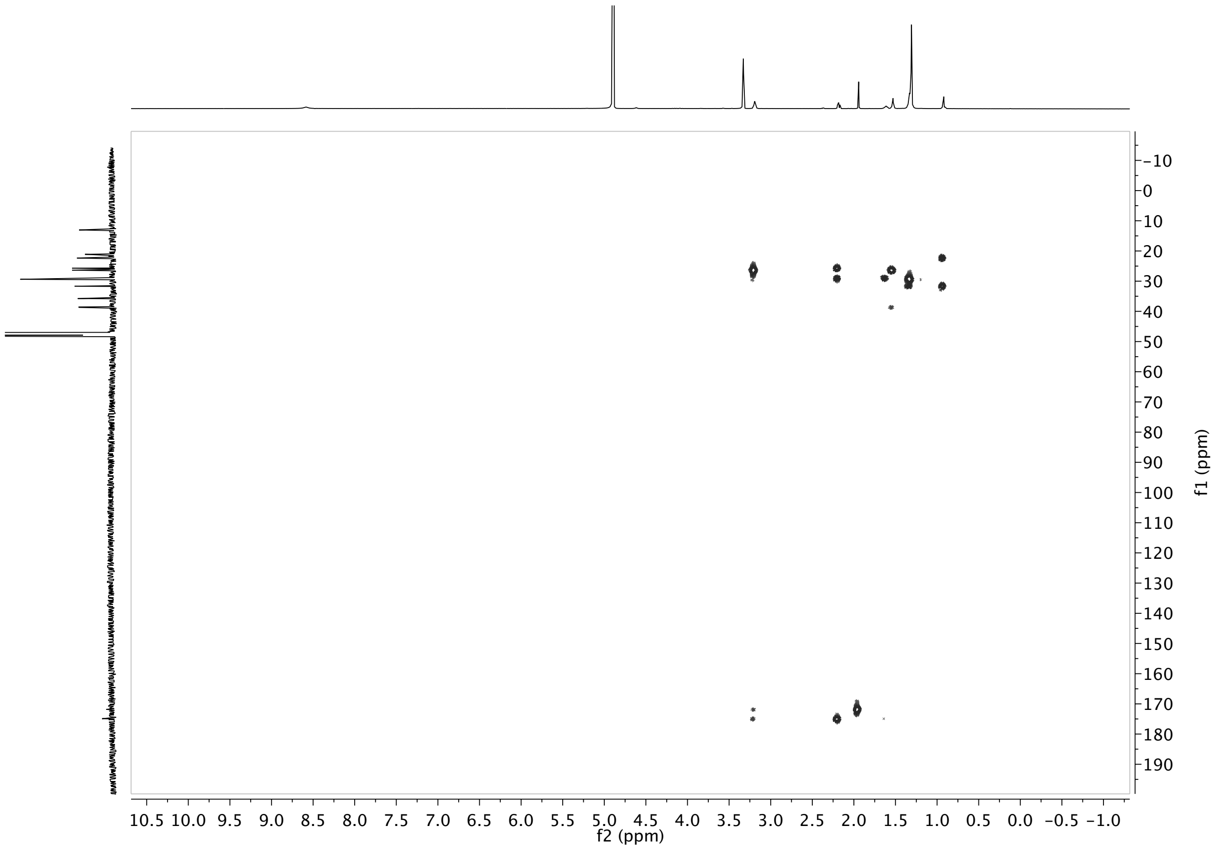


**Figure e7.** HMBC spectrum of synthetic **NAPP** in MeOH

**2.3 NMR spectra of synthetic *N*-Acetyl-Myristoyl-Putrescine (NAMP)**

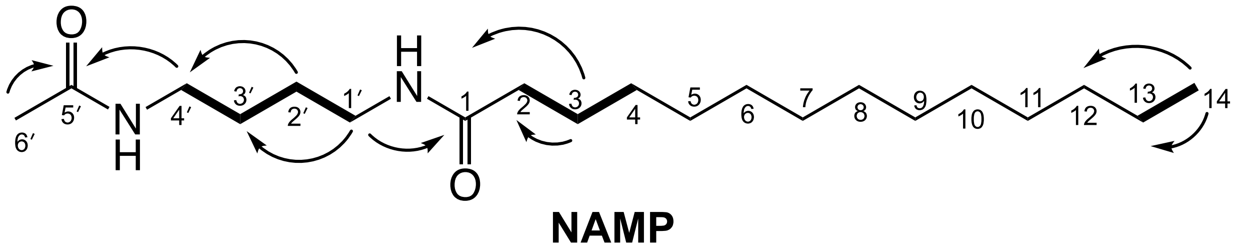


**Figure f1.** Structure of synthetic **NAMP** and key ⟶ and COSY ▬ correlations.

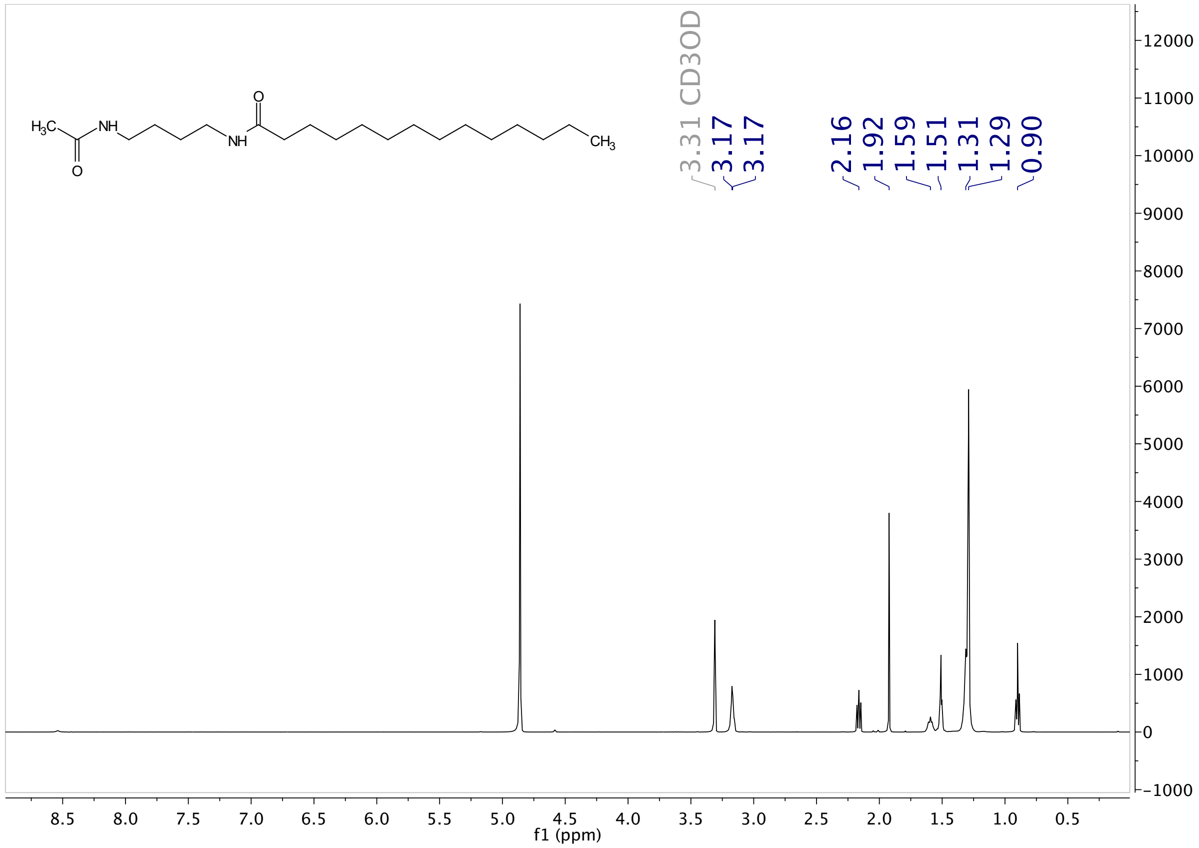


**Figure f2.** [1]H NMR spectrum of synthetic **NAMP** in MeOD

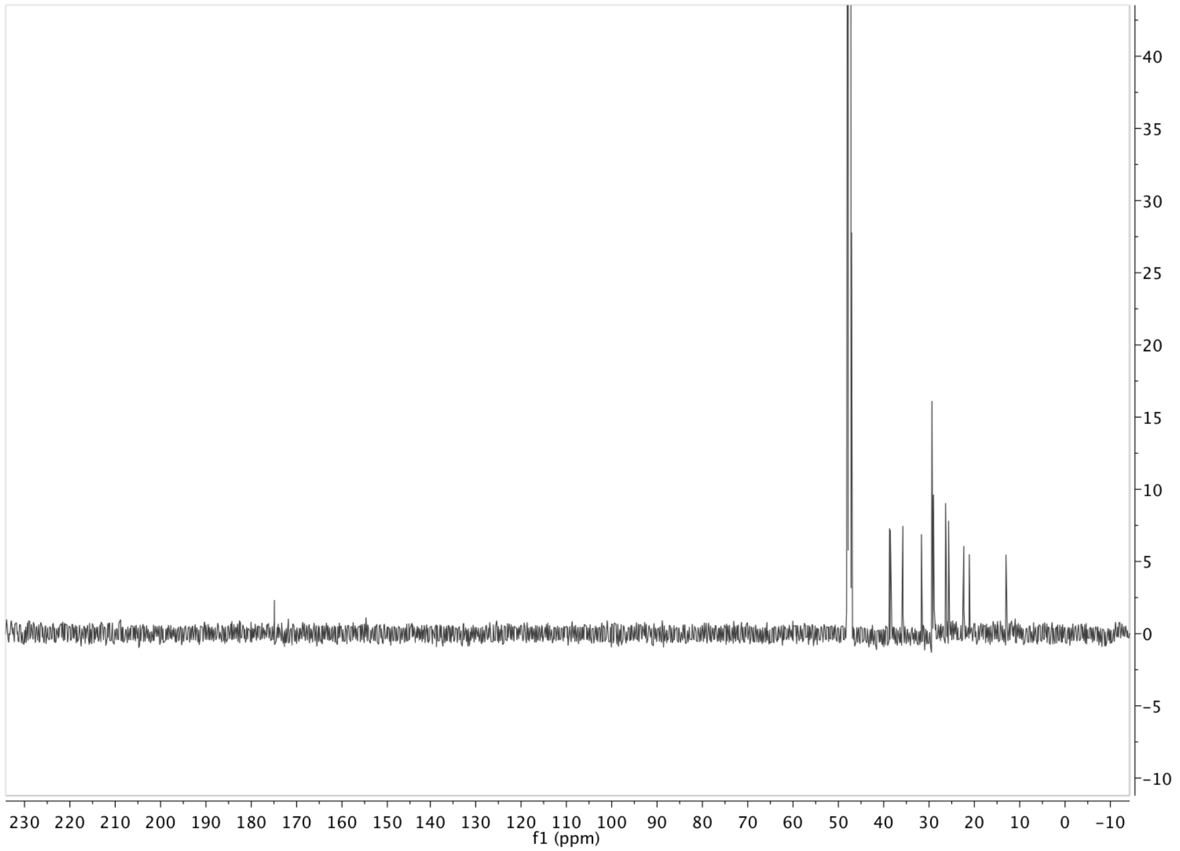


**Figure f3.** $^{13}C$ NMR spectrum of synthetic **NAMP** in MeOD

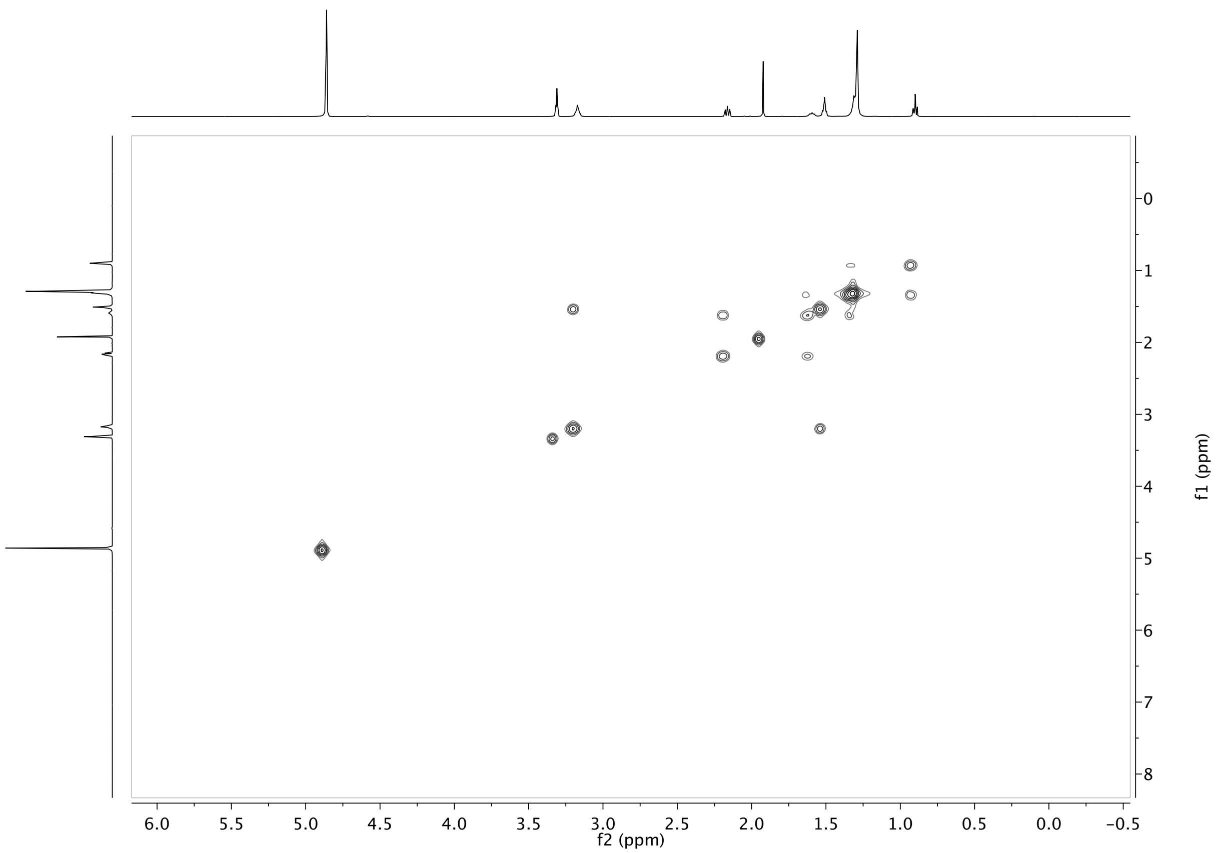


**Figure f4.** COSY NMR spectrum of synthetic **NAMP** in MeOD

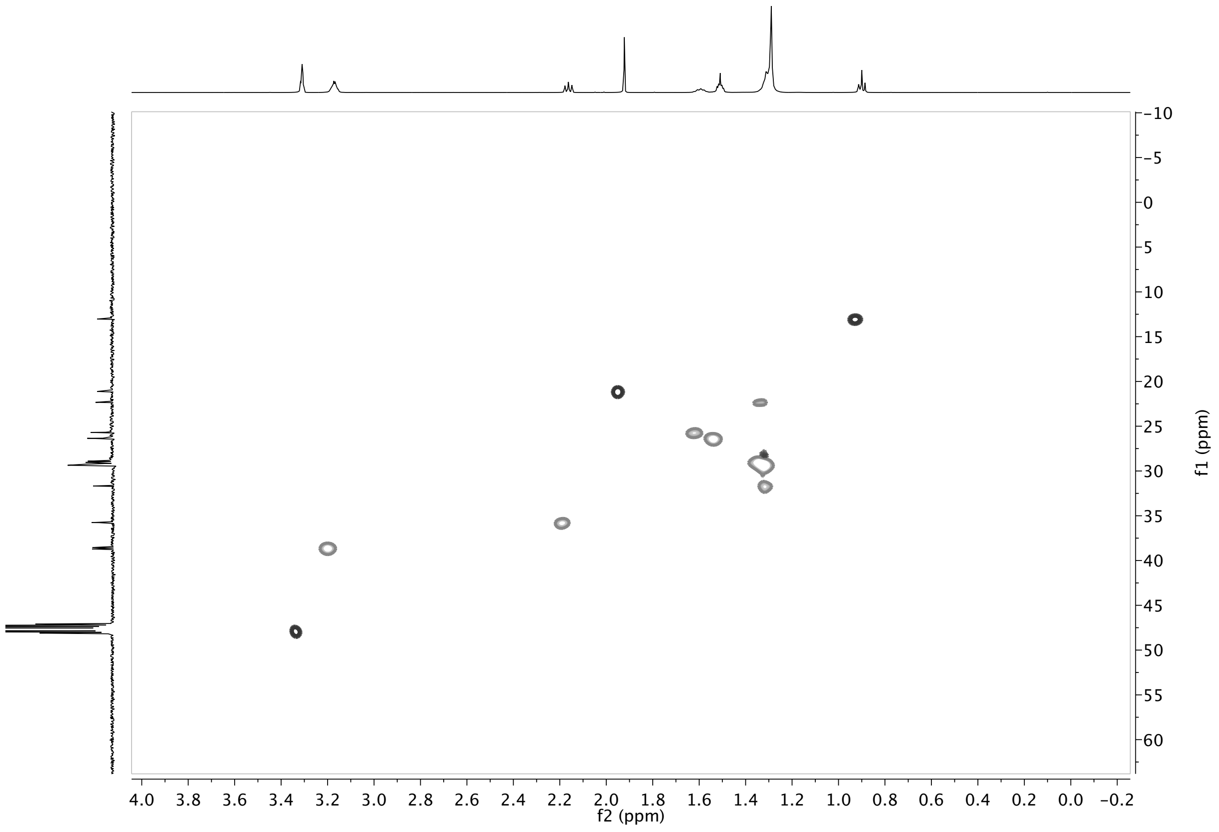


**Figure f5.** HSQC NMR spectrum of synthetic **NAMP** in MeOD

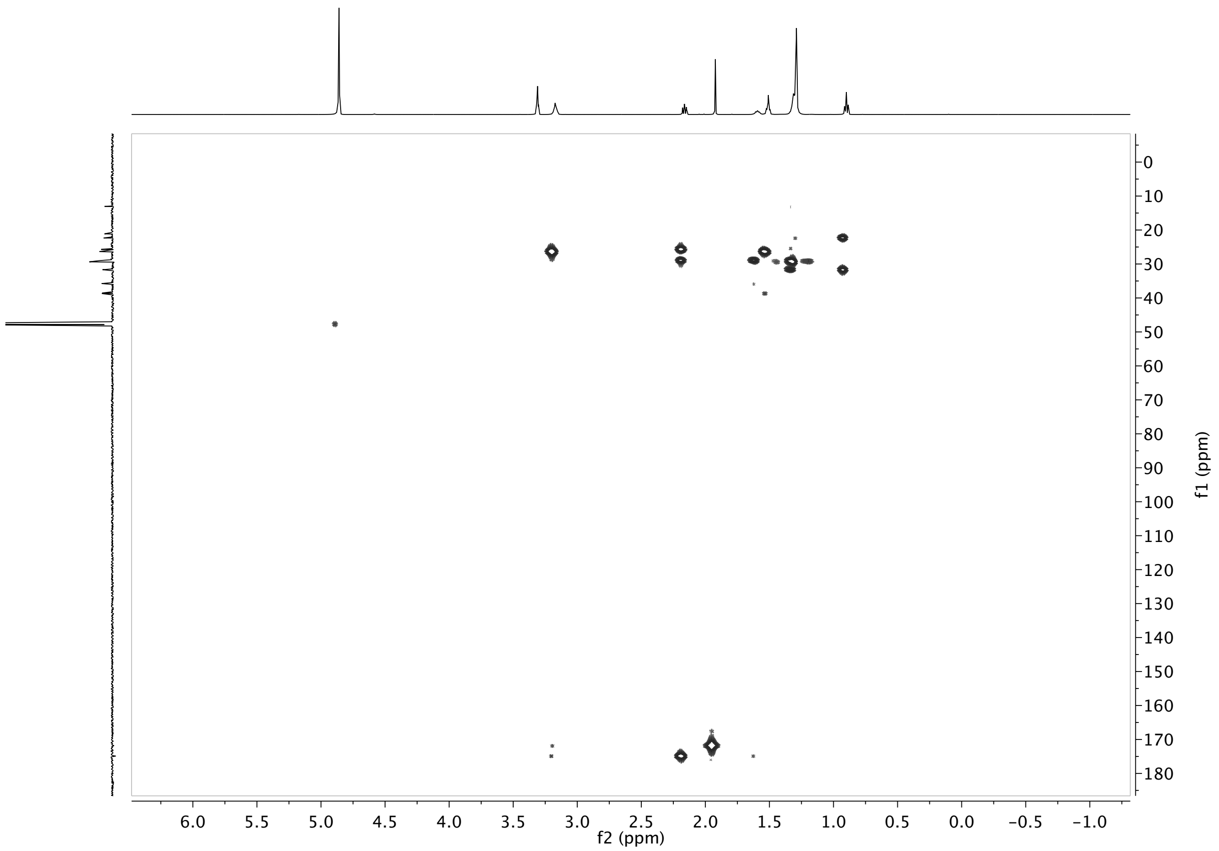


**Figure f6.** HMBC NMR spectrum of synthetic **NAMP** in MeOD

#### 2.4 NMR spectra of synthetic *N*-Myristoyl-L-Lysine

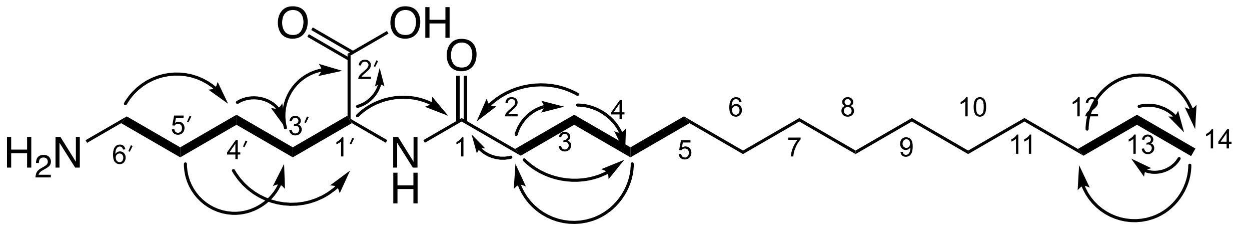


**NML**

**Figure g1.** Structure of synthetic **NML** and key → and COSY ▬ correlations.

**Table f.** NMR data for synthetic **NML** (500 MHz in MeOD)

| Position | δH, mult (*J* in Hz) | δC | HMBC | COSY |
|---|---|---|---|---|
| **1** | | 174.3 | | |
| **2** | 2.23, t (7.57,7.57) | 35.7 | 25.2, 28.3,174.3 | H-3 |
| **3** | 1.61, m | 25.2 | 28.3, 35.7, 174.3 | H-2, H-4 |
| **4** | 1.32 | 28.3 | 25.2, 35.7 | H-3, H-5 |
| **5** | 1.32 | 28.4 | 25.2 | |
| **6** | 1.32 | 28.4 | | |
| **7** | 1.32 | 28.6 | | |
| **8** | 1.32 | 28.7 | | |
| **9** | 1.32 | 28.7 | | |
| **10** | 1.32 | 28.8 | | |
| **11** | 1.32 | 28.8 | | |
| **12** | 1.29 | 31.0 | 12.4, 21.7, 28.8 | H-13 |
| **13** | 1.29, br, m | 21.7 | 12.4, 31.0 | H-14, H-12 |
| **14** | 0.90, t (6.88,6.88) | 12.4 | 21.7, 31.0 | H-13 |
| **1′** | 4.25, m | 53.2 | 21.4, 31.1, 174.3, 176.8 | H-3′ |
| **2′** | | 176.8 | | |
| **3′** | 1.66, 1.82m | 31.1 | 21.4, 25.9, 53.2,176.7 | H-1′, H-4′ |
| **4′** | 1.42, m | 21.4 | 31.1, 38.2, 53.2 | H-3′, H-5′ |
| **5′** | 1.68, m | 25.9 | 21.4, 31.1,38.2 | H-3′, H-4′ |
| **6′** | 2.90, t (7.09,7.09) | 38.2 | 25.9, 21.4 | H-5′ |

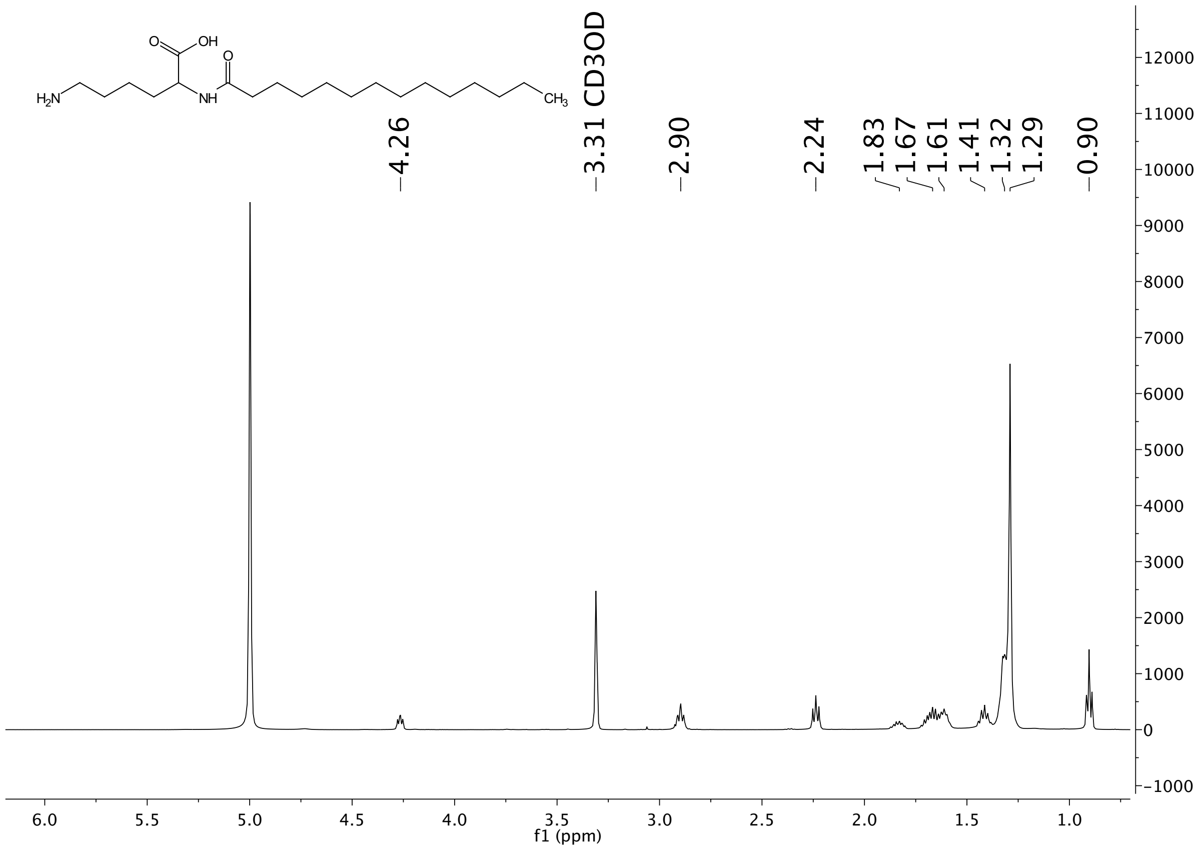


**Figure g2.** $^1$H NMR spectrum of synthetic **NML** in MeOD

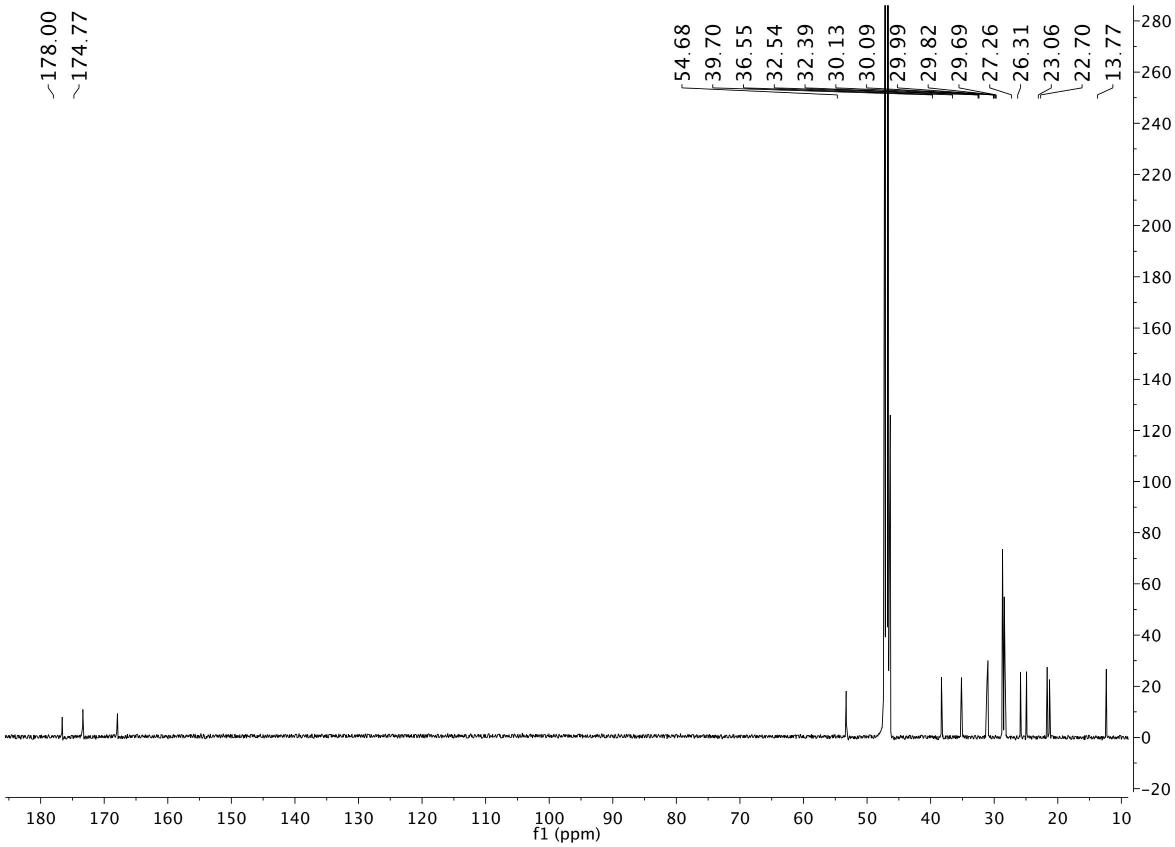


**Figure g3.** $^{13}C$ NMR spectrum of synthetic **NML** in MeOD

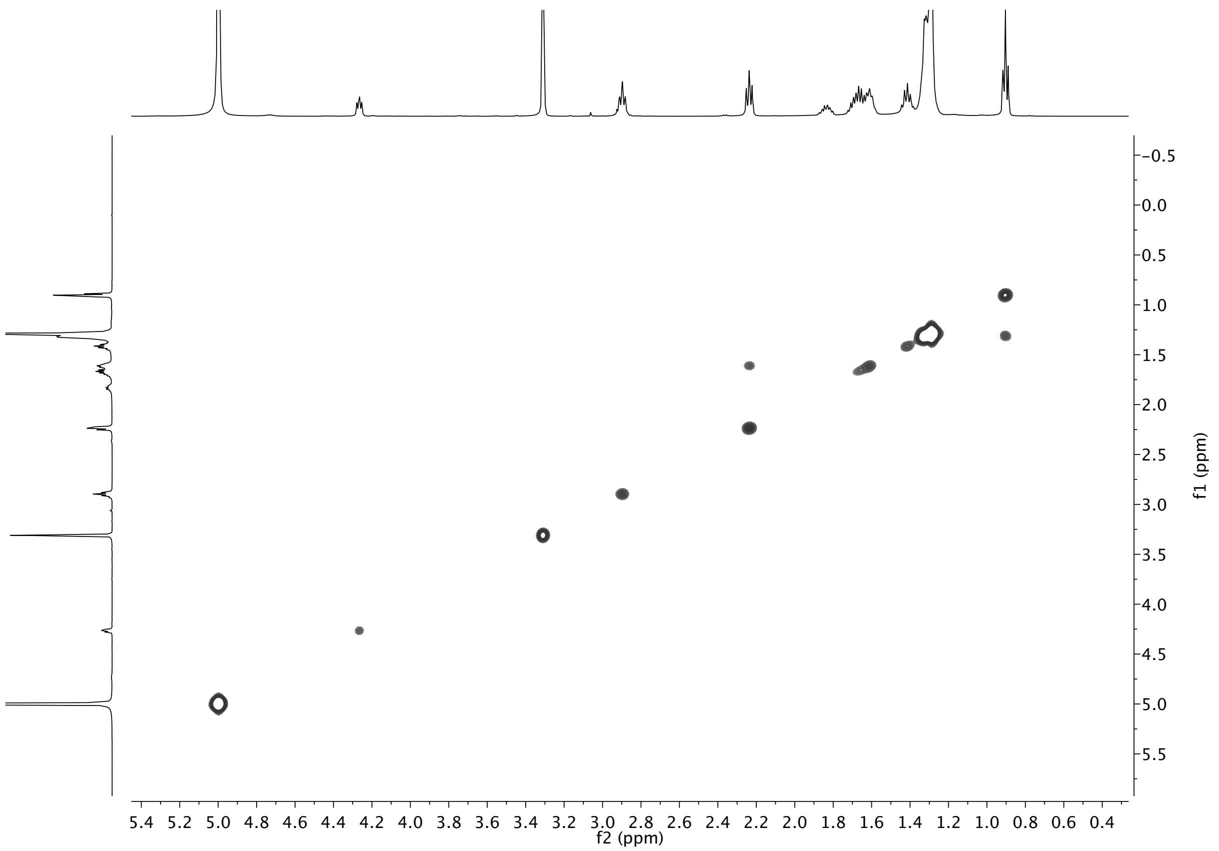


**Figure g4.** COSY NMR spectrum of synthetic **NML** in MeOD

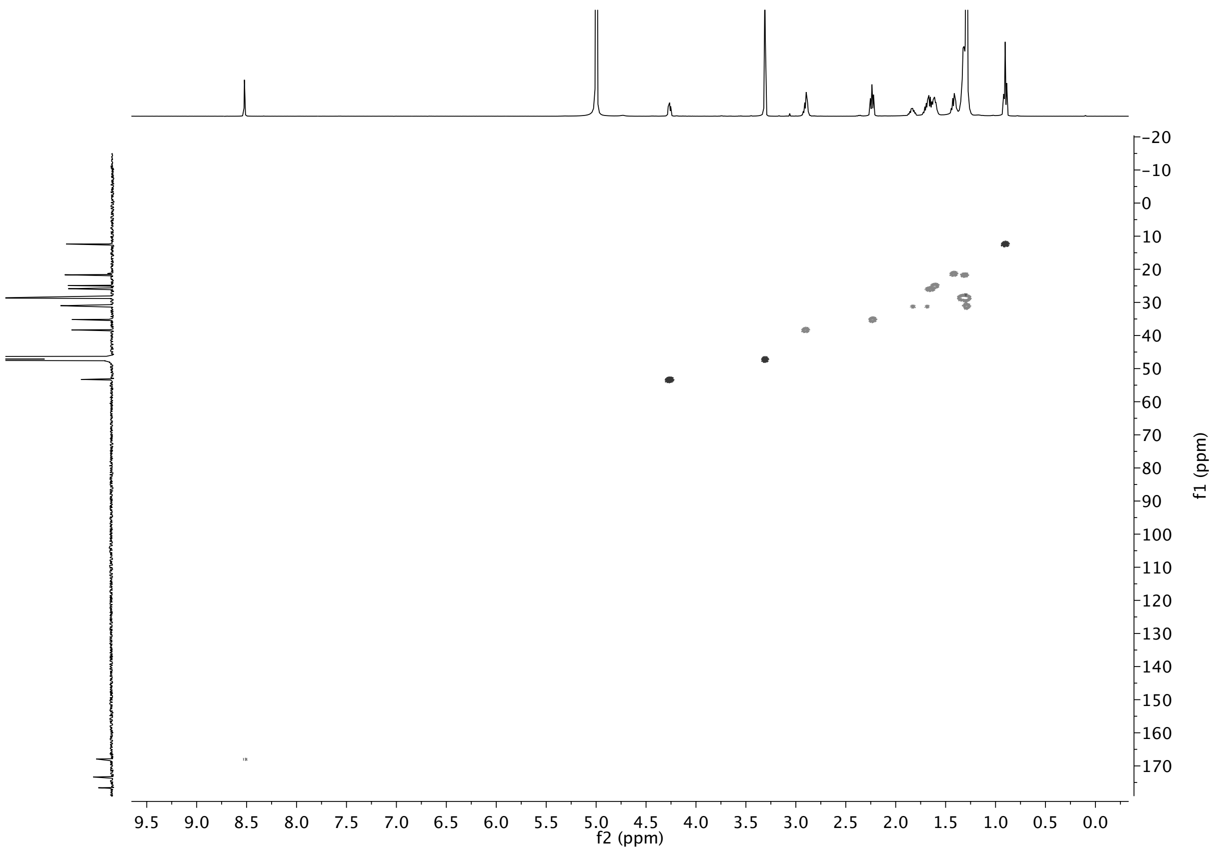


**Figure g5.** HSQC NMR spectrum of synthetic **NML** in MeOD

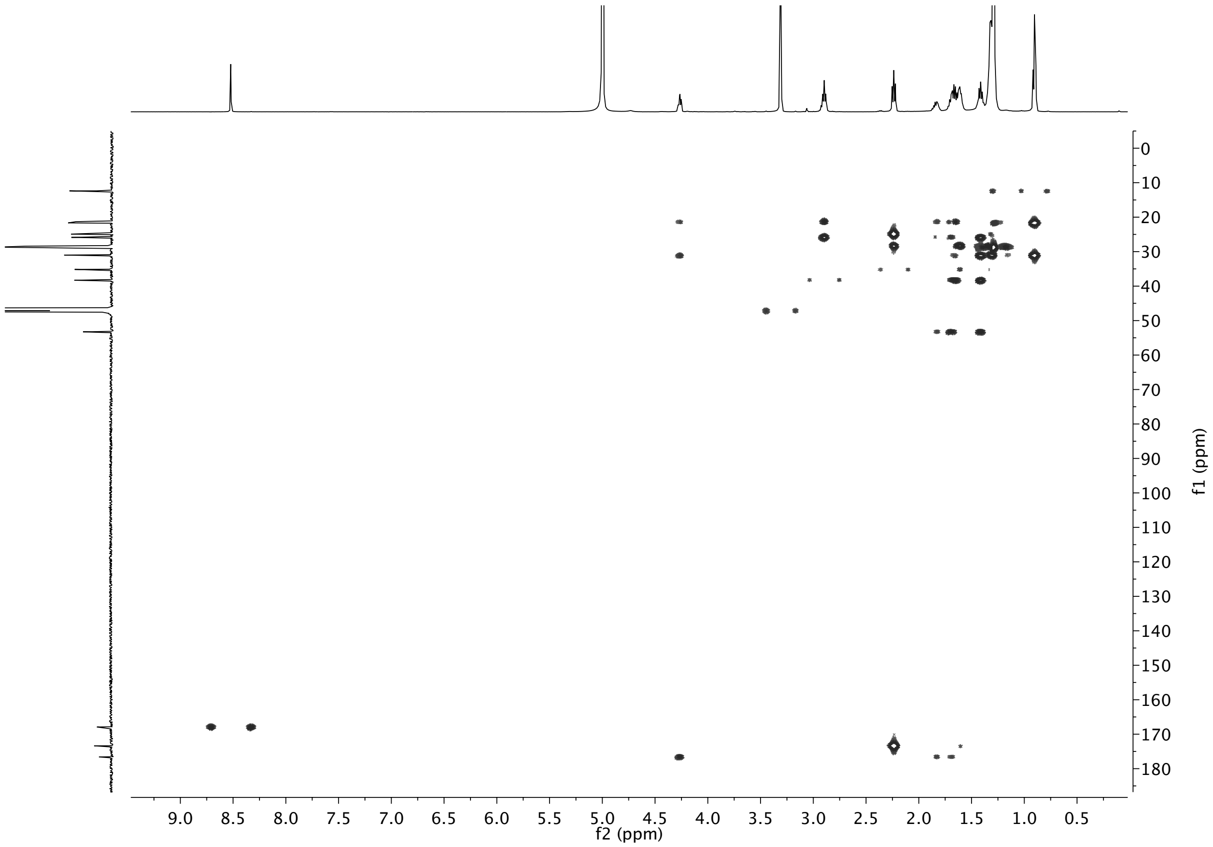


**Figure g6.** HMBC NMR spectrum of synthetic **NML** in MeOD
