## Supplementary material for "A meta-analysis of the gut microbiome in inflammatory bowel disease patients identifies disease-associated small molecules": Methods

### STAR ★METHODS

### KEY RESOURCES TABLE

| Reagent or Resource | Source | Identifier |
|---|---|---|
| **Chemicals** | | |
| **10mM dNTP mix** | Life Technologies | 18427-088 |
| **1,4-Dithiothreitol (DTT)** | Sigma (Roche) | 10197777001 |
| **2-Propanol (HPLC)** | Fisher Scientific | A451-4 |
| **2XYT Broth** | Fisher Scientific | BP9743500 |
| **1,4-Diaminobutane** | Sigma-Aldrich | D13208 |
| **Acetonitrile** | Fisher Scientific | A998-4 |
| **Ammonium Chloride** | Sigma-Aldrich | A4514 |
| **Ampicillin** | Sigma-Aldrich | A1593 |
| **BD Bacto™ Dehydrated Agar** | Fisher Scientific | 214010 |
| **Gibco™ Bacto™ Peptone** | Fisher Scientific | DF0118-17-0 |
| **Beef Extract** | Fisher scientific | S25661A |
| **Brain Heart Infusion** | VWR | 90003-032 |
| **Butyric acid** | Sigma-Aldrich | B103500 |
| **Calcium Chloride** | Sigma-Aldrich | C8106 |
| **Carbenicillin Disodium Salt** | Sigma-Aldrich | C1389 |
| **Chloroform-D** | Cambridge Isotope Laboratories | DLM-29-10 |
| **D-(+)-Glucose** | Sigma-Aldrich | G8270 |
| **D-(+)-Maltose monohydrate** | Sigma-Aldrich | M5895 |
| **D-Cellobiose** | Santa Cruz Biotechnology | 280654 |
| **D-(−)-Fructose** | Sigma (Millipore) | 47740 |
| **Dichloromethane** | Fisher Scientific | AC406920040 |
| **Dextran sulfate sodium salt, colitis grade (36,000 - 50,000)** | MP Biomedical | 0216011090 |
| **Gibco™ DMEM** | Thermo Fisher | 11995073 |
| **Gibco™ DPBS** | Thermo Fisher | 14040133 |
| **Ethyl Acetate (HPLC)** | Fisher Scientific | E195-4 |
| **Fmoc-L-Lys(Boc)-Wang Resin** | Matrix Innovation | FL-200-1201 |
| **Formic acid (LC/MS)** | Fisher Scientific | A117-50 |
| **Fluorescein isothiocyanate–dextran** | Sigma-Aldrich | FD4 |
| **Gifu Anaerobic Medium (GAM) Broth, Modified** | HyServe GmbH | 1005433 |
| **Biotium GelRed Nucleic Acid Gel Stain** | Fisher Scientific | NC9524151 |
| **Hemin** | Sigma-Aldrich | 51280 |

*(continued next page)*

***Continued***

| | | |
|---|---|---|
| **HyClone Iron-Supplemented Calf Serum** | Cytiva | SH30072.04 |
| **IPTG, dioxane-free** | Thermo Scientific | R0393 |
| **Isovaleric acid** | Sigma-Aldrich | 129542 |
| **L-Cysteine** | Fisher Scientific | ICN19464625 |
| **L-Cysteine Hydrochloride Monohydrate** | FUJIFILM Wako Pure Chemical | 033-05272 |
| **Reagent or Resource** | Source | Identifier |
| **LB Broth (Miller)** | Sigma-Aldrich | L3522 |
| **M17 Broth** | Fisher Scientific | OXCM0817B |
| **Magnesium sulfate heptahydrate** | Sigma-Aldrich | 230391 |
| **Meat extract** | Sigma (Millipore) | 70164 |
| **Methanol (HPLC)** | Fisher Scientific | A452-4 |
| **Methanol-$D_4$** | Cambridge Isotope Laboratories | DLM-51 |
| **Methylene Chloride (HPLC)** | Fisher Scientific | D151-1 |
| **Myristoyl chloride** | Fisher Scientific | AC180631000 |
| ***N*-(4-Aminobutyl)acetamide** | Sigma-Aldrich | CDS021099 |
| **N-(4-Aminobutyl)tetradecanamide** | Sigma (Enamine) | ENAH908234CC |
| **N-Lauroyl-L-lysine** | Santa Cruz Biotechnology | 491941 |
| **N-Myristoyl Lysine** | Ambinter | 16228299 |
| ***N,N*-Dimethyl Formamide** | Sigma-Aldrich | D4551-500ml |
| **Palmitoyl chloride** | Fisher Scientific | AC304291000 |
| **Paraformaldehyde Solution, 4% in PBS** | Fisher Scientific | AAJ19943K2 |
| **Piperidine solution** | Sigma-Aldrich | 80645 |
| **Potassium phosphate monobasic** | Sigma-Aldrich | P5655 |
| **Potassium phosphate dibasic** | Sigma-Aldrich | 795496 |
| **Propionic acid** | Sigma-Aldrich | P5561 |
| **Pyridine** | Fisher Scientific | AC610991000 |
| **Radiant™ SYBR Green Master Mix qPCR Kit** | Alkali Scientific | QS1020 |
| **BD Difco™ Reinforced Clostridial Medium** | Fisher Scientific | DF1808-17-3 |
| **Resazurin sodium salt** | Santa Cruz Biotechnology | 62758-13-8 |
| **Sodium bicarbonate** | Sigma-Aldrich | S6014 |
| **Sodium chloride** | Sigma-Aldrich | S7653 |
| **Sodium hydroxide** | Sigma-Aldrich | 795429 |
| **Spectinomycin dihydrochloride pentahydrate** | Sigma-Aldrich | S4014 |
| **Streptomycin sulfate salt** | Sigma-Aldrich | S9137 |
| **BD Difco™ Terrific Broth** | Fisher Scientific | DF0438-17 |

*(continued next page)*

| *Continued* | | |
|---|---|---|
| **Trace Mineral Supplement** | ATTC | MD-TMS |
| **Trifluoroacetic acid** | Sigma-Aldrich | 302031 |
| **Triisopropylsilane** | Sigma-Aldrich | 233781 |
| **BD Bacto™ Tryptic Soy Broth** | Fisher Scientific | DF0370-17-3 |
| **Gibco™ Trypan Blue Solution** | Thermo Scientific | 15250061 |
| **Trypticase Peptone** | VWR | 90000-434 |
| **TWEEN® 80** | Sigma-Aldrich | P1754 |
| **Vitamin $K_1$** | Sigma-Aldrich | 95271 |
| **Vitamin Supplement** | ATTC | MD-VS |
| **Difco™ Yeast Extract** | VWR | 90000-026 |
| **Experimental models: Organisms/Strains/Cell lines** | | |
| **Caco-2 [Caco2] (Human colonic epithelial cells)** | ATCC | HTB-37 |
| ***Clostridium clostridioforme* WAL-7855** | BEI Resources | HM-317 |
| ***Clostridium bolteae* ATCC BAA-613 (WAL 16351, CCUG 46953)** | ATCC | BAA-613 |
| ***Clostridium Sporogenes* ATCC 15579** | ATCC | 15579 |
| **Competent *E. coli* BL21(DE3)** | New England Biolabs (NEB) | C2527I |
| ***E. coli* BL21(DE) harboring pGFPuv- smNRPS-ebf and pYS20-sfp** | This paper | NA |
| ***E. coli* BL21(DE) harboring pUC57- smNRPS-ecf and pYS20-sfp** | This paper | NA |
| ***E. coli* BL21(DE) harboring pGFPuv pYS20-sfp** | This paper | NA |
| ***E. coli* BL21(DE) harboring pUC57 pYS20-sfp** | This paper | NA |
| ***E. coli* Stellar™ Competent Cells** | Takara Bio | 636763 |
| **pUC57 plasmid DNA** | GenScript | SD1176 |
| **pUC57-smNRPS-ecf** | This paper | NA |
| **pGFPuv Vector** | Takara Bio | 632312 |
| **pGFPuv-smNRPS-ebf** | This paper | NA |
| **pYS20-sfp** | This paper | NA |
| **Sequence-Based Reagents** | | |
| **PC016_SM_ebf_ pGFP-UV primers** | Forward: CAGGAAACAGCTATGAAAACGCCAGACTTGACTG<br>Reverse: AGTTGGAATTCATTAAAATCTTTTAATTTTCTTTGACGGCG | Amplification of *ebf* |

(continued next page)

*Continued*

| | | |
|---|---|---|
| **pGFP-IF primers** | Forward: TAATGAATTCCAACTGAGCGCCGG<br>Reverse: CATAGCTGTTTCCTGTGTGAAATTG | Linearization of pGFP-UV |
| **sfp-duet primers** | Forward: AAGGAGATATACATATGAAGATTTACGGAATTTATATG<br>Reverse: TTGAGATCTGCCATATTATAAAAGCTCTTCGTACG | Amplification of *sfp* |
| **ecf-Eco primers** | Forward: TATGAATACTATGACCGTGATG<br>Reverse: ACGGCTGGTAATATACACTTC | Screening for *ecf* |
| **PC033_ebf_seq/PC014_SM_ebf primers** | Forward: TCGCCGGTTATCAGCACC<br>Reverse: GGTCTGATCGGATCCTTAAAATCTTTTAATTTTCTTTGACGGCG | Screening for *ebf* |
| **sfp primers** | Forward: TCATAAAGAAGATGCTCACC<br>Reverse: TAATAAAAGCTCTTCGTACGAG | Screening for *sfp* |
| **β-actin primers** | Forward: AAGTGTGACGTTGACATCCG<br>Reverse: GATCCACATCTGCTGGAAGG | Amplification of *β-actin* |
| **TNF-α primers** | Forward: GCCTCCCTCTCATCAGTTCT<br>Reverse: CACTTGGTGGTTTGCTACGA | Amplification of *TNF-α* |
| **IL-6 primers** | Forward: GAGGATACCACTCCCAACAGACC<br>Reverse: AAGTGCATCATCGTTGTTCATACA | Amplification of *IL-6* |
| **IL-1β primers** | Forward: CAACCAACAAGTGATATTCTCCATG<br>Reverse: GATCCACACTCTCCAGCTGCA | Amplification of *IL-1β* |

*(continued next page)*

| *Continued* | | |
|---|---|---|
| **IL-17A primers** | Forward: TTTAACTCCCTTGGCGCAAAA Reverse: CTTTCCCTCCGCATTGACAC | Amplification of *IL-17A* |
| **CCL2 primers** | Forward: GTTGGCTCAGCCAGATGCA Reverse: AGCCTACTCATTGGGATCATCTTG | Amplification of *CCL2* |
| **IL-12p40 primers** | Forward: AGTGACATGTGGAATGGCGT Reverse: CAGTTCAATGGGCAGGGTCT | Amplification of *IL-12p40* |
| **Software and Algorithms** | | |
| **Adobe Illustrator** | Adobe | NA |
| **Agilent Masshunter Workstation** | Agilent | NA |
| **Agilent Qualitative analysis** | Agilent | NA |
| **antiSMASH 5.1.1** | (Blin et al., 2019) | https://antismash.secondarymetabolites.org |
| **Blast 2.7.1+** | NCBI | https://blast.ncbi.nlm.nih.gov/Blast.cgi |
| **ChemDraw Professional 16.0** | PerkinElmer | NA |
| **SRA Toolkit Release 2.10.0** | NCBI | https://github.com/ncbi/sra-tools |
| **PRINSEQ-lite 0.20.4B** | (Schmieder and Edwards, 2011) | https://prinseq.sourceforge.net/index.html |
| **seqIO_extract** | Davis E | https://github.com/davised/seqIO_extract |
| **Geneious** | Geneious | http://www.geneious.com/ |
| **PRINSEQ-lite v0.20.4B** | (Schmieder and Edwards, 2011) | http://prinseq.sourceforge.net/ |
| **SPAdes v3.11.0** | (Bankevich et al., 2012) | https://github.com/ablab/spades |
| **PhyloPhlAn (release 28 April 2020)** | (Asnicar et al., 2020) | https://github.com/biobakery/phylophlan |
| **R version 4.3.1 (Nickname: Beagle Scouts)** | R Core Team (2023) | https://www.R-project.org |
| **RStudio 2023.3.0.386 (Release name: Cherry Blossom)** | Posit team (2023) | https://www.rstudio.com/ |
| **ggplot2 2.2.1** | Wickham H (2009) | https://cran.r-project.org/web/packages/ggplot2/index.html |

*(continued next page)*

***Continued***

| | | |
|---|---|---|
| **ggpubr 0.1.6** | Kassambara, 2017 | https://cran.r-project.org/web/packages/ggpubr/index.html |
| **ape 5.7-1** | Paradis E, et al. (2023) | https://cran.r-project.org/web/packages/ape/ |
| **Boruta 8.0.0** | Kursa MB and Rudnicki WR (2010) | https://cran.r-project.org/web/packages/Boruta/ |
| **caret 6.0-94** | Kuhn M (2008) | https://cran.r-project.org/web/packages/dabestr/ |
| **dabestr 2023.9.12** | Ho J, et al. (2019) | https://cran.r-project.org/web/packages/dabestr/ |
| **DescTools 0.99.52** | Signorell A, et al. (2023) | https://cran.r-project.org/web/packages/DescTools/ |
| **drc 3.0-1** | (Ritz et al., 2016) | https://cran.r-project.org/web/packages/drc/ |
| **ggbreak 0.1.2** | Shuangbin Xu, Chen M, Feng T, Zhan L, Zhou L, Yu G (2021) | https://cran.r-project.org/web/packages/ggbreak/ |
| **ggpubr 0.6.0** | Kassambara A (2023) | https://cran.r-project.org/web/packages/ggpubr/ |
| **ggrepel 0.9.4** | Slowikowski K (2023) | https://cran.r-project.org/web/packages/ggrepel/ |
| **ggtree 3.10.0** | (Yu et al., 2017) | https://github.com/YuLab-SMU/ggtree |
| **jcolors 0.0.5** | Jared Huling | https://github.com/jaredhuling/jcolors |
| **lvplot 0.2.1** | Wickham H | https://cran.r-project.org/web/packages/lvplot/ |
| **metafor 4.4-0** | Viechtbauer W (2010) | https://cran.r-project.org/web/packages/metafor/ |
| **metaviz 0.3.1** | Kossmeier M, et al. | https://cran.r-project.org/web/packages/metaviz/ |
| **missForest 1.5** | Stekhoven DJ (2022) | https://cran.r-project.org/web/packages/missForest/ |

*(continued next page)*

| *Continued* | | |
|---|---|---|
| **pROC 1.18.5** | Robin X, et al (2011) | https://cran.r-project.org/web/packages/pROC/ |
| **randomForest 4.7-1.1** | Liaw A and Wiener M (2002) | https://cran.r-project.org/web/packages/randomForest/ |
| **reshape2 1.4.4** | Wickham H (2020) | https://cran.r-project.org/web/packages/reshape2/ |
| **ROCR 1.0-11** | Sing T, et al. (2005) | https://cran.r-project.org/web/packages/ROCR/ |
| **rstatix 0.7.2** | Kassambara A (2023) | https://rpkgs.datanovia.com/rstatix/ |
| **scales 1.3.0** | Wickham H | https://cran.r-project.org/web/packages/scales/ |
| **strex 1.6.1** | - | https://cran.r-project.org/web/packages/strex |
| **stringi 1.8.3** | Gagolewski M (2022) | https://cran.r-project.org/web/packages/stringi |
| **tidyverse 1.2.1** | Wickham H (2019) | https://www.tidyverse.org/ |
| **jsonlite 1.8.8** | Ooms J (2014) | https://cran.r-project.org/web/packages/jsonlite |
| **Antibodies Assays** | | |
| **Critical Commercial Assays** | | |
| **E.Z.N.A.® Total RNA Kit I** | Omega Bio-Tek | R6834-01 |
| **High-Capacity RNA-to-cDNA™ Kit** | Thermo Scientific | 4387406 |
| **In-Fusion® HD Cloning System** | TAKARA | 639646 |
| **Quick-DNA Fungal/Bacterial Miniprep Kit** | Zymo Research | D6005 |
| **QIAquick Gel Extraction Kit** | QIAGEN | 28706 |
| **QIAprep Spin Miniprep Kit** | QIAGEN | 27106 |
| **RealTime-Glo™ MT cell viability assay kit** | Promega | G9713 |
| **Zymoclean Large Fragment DNA Recovery** | Zymo Research | D4046 |
| **DNeasy® PowerSoil® Kit** | QIAGEN | 12888 |

*(continued next page)*

| Continued | | |
|---|---|---|
| **Animal Models** | | |
| **Dextran sulfate sodium (DSS)–induced colitis model** | The Jackson Laboratory | JAX 000664 |
| **IL-10–deficient colitis model** | The Germ-Free Facility at the University of Michigan | NA |
| **Others** | | |
| **Agilent 6120 quadrupole mass spectrometer** | Agilent | NA |
| **Agilent 6530 Q-TOF LC/MS equipment** | Agilent | NA |
| **Agilent Poroshell 120 EC-C18 2.6 um (4.6x100mm)** | Agilent | NA |
| **Reservoir-2 Frits, 20ml** | Agilent | 12131017 |
| **Mega BE-C18, 10mg** | Agilent | 12256031 |
| **Anaerobic chamber, Vinyl, Type B** | Coy Laboratory Products | NA |
| **NMR Bruker AVANCE III 500mHz** | Bruker | NA |
| **Tecan Spark® multimode microplate reader** | Tecan | NA |

### CONTACT FOR REAGENT AND RESOURCE SHARING

Further information may be obtained from the Lead Contact Mohamed S. Donia (; address: Department of Molecular Biology, Princeton University, Princeton, New Jersey, 08544, USA).

### EXPERIMENTAL DESIGN

#### 1. *Construction of plasmids for heterologous expression*

##### *1.1. Construction of the ecf expression vector (pUC57-ecf)*

The DNA sequence of *ecf* (resulting from the metagenomic analysis of the MetaHIT cohort) was codon-optimized for expression in *E. coli*, and synthesized as a single DNA fragment fused to the *lac* promoter in pUC57 (GenScript, USA). No changes in the final protein sequence were introduced during optimization. The resulting vector was named pUC57-ecf.

##### *1.2. Construction of the ebf expression vector (pGFPuv-ebf)*

*Clostridium bolteae* ATCC BAA-613 was grown in an anaerobic chamber at 37°C in modified PYG media (according to the DSMZ recipe, DSMZ GmbH) and genomic DNA was extracted using the Quick-DNA Fungal/Bacterial Miniprep Kit (Zymo Research, USA). PC016F_SM_ebf_ pGFP-UV and PC016R_SM_ebf_ pGFP-UV primers were used to amplify the *ebf* BGC from the obtained genomic DNA. In parallel, pGFP-IF-F and pGFP-IF-R primers were used to amplify the 2.5 kbps linear fragment of pGFPuv vector. The amplicons were assembled by using In-Fusion® HD Cloning Kit (Takara Bio, Japan) following the manufacturer protocol, where the final design puts *ebf* under the control of the constitutive *lac* promoter in pGFPuv, replacing *gfp*. The assembly reaction mixture was further purified using Zymo DNA Clean and Concentrator Kit (Zymo Research, USA), and transformed to *E. coli* Stellar cells. The resulting construct was verified using restriction enzyme digestion and Sanger sequencing and named pGFPuv-smNRPS-ebf.

#### 1.3. Construction of an sfp expression vector (pYS20)

The *sfp* gene (accession number CP019663.1) was amplified from *B. subtilis* 168 using primers sfp-duet_F and sfp-duet_R then cloned into NdeI-digested pCDF-Duet-1 (Novagen, USA) using the In-Fusion® HD Cloning Kit (Takara Bio, Japan) following the manufacturer protocol. The final design puts *sfp* under the control of the inducible T7 promoter in pCDF-Duet-1. The assembly reaction was then transformed into *E. coli* Stellar cells (Takara Bio, Japan) following the manufacturer protocol, and the resulting construct was verified using restriction enzyme digestion and Sanger sequencing.

### 2. Transformation of E. coli BL21 with constructed plasmids

#### 2.1. pGFPuv-smNRPS-ebf

Electrocompetent *E. coli* BL21 cells were transformed with pGFPuv-smNRPS-ebf and pYS20 to yield *E. coli* BL21_smNRPS-ebf_sfp. For a control strain, *E. coli* BL21 was transformed with pGFPuv and pYS20 to yield *E. coli* BL21_pGFPuv_sfp. Upon transformation, transformants that are resistant to carbenicillin and streptomycin were inoculated in LB supplemented with 100 ug/mL carbenicillin and streptomycin at 30°C, 200 rpm overnight for plasmid extraction. Plasmids were extracted using QIAprep Miniprep kit (QIAGEN, USA) and further verified by PCR using PC033_ebf_seq_F, PC014R_SM_ebf, sfp_F and sfp_R, and restriction enzyme digestion.

#### 2.2. pUC57-smNRPS-ecf

Similarly, pUC57-smNRPS-ecf and pYS20 were transformed into electrocompetent *E. coli* BL21 and selected on both 100 ug/mL carbenicillin and streptomycin. Transformants were grown in selective LB medium, and plasmids were extracted and further verified by PCR using ecf-E.co_F, ecf-E.co_R, sfp_F and sfp_R, and restriction enzyme digestion.

### 3. Heterologous expression of ecf and ebf in E. coli BL21

A single colony of each expression cell line (*E. coli* BL21_smNRPS-ebf_sfp or *E. coli* BL21_smNRPS-ecf_sfp) or an empty vector control cell line (*E. coli* BL21_pGFPuv_sfp or *E. coli* BL21_pUC57_sfp) was used to inoculate a 2 mL LB broth culture (10 g tryptone, 5 g yeast extract, 10 g NaCl, 100 ug/mL of both carbenicillin and streptomycin), which was incubated at 30°C with shaking at 200 rpm. After overnight cultivation, 1mL of the seed culture was used to inoculate 50 mL of LB medium containing the corresponding antibiotics. This expression culture was shaken at 200 rpm, IPTG was added at O.D. 0.25-0.5 and the expression continued for 72 additional hours. The cultures were then spun down, and bacterial cell pellets and cell-free media were processed separately. For the pellets, spun-down cells were suspended in 20 mL MeOH and sonicated for 20 min. Cell debris were then spun down and collected supernatant was dried under vacuum using a rotary evaporator. Cell-free media were extracted using Diaion HP-20 resin, 2.5 g activated resin were added to the 50 mL supernatant and the mixture was incubated for 2 hours at room temperature at 120 rpm shaking condition. The mixture was put into a column (Empty SPE cartridge, Agilent USA) and the resin was washed by 20mL milli-Q water and eluted with 20 mL MeOH. The organic solution was dried under vacuum using a rotary evaporator.

Dried extracts from pellet and cell-free media extracts were re-dissolved in 1 mL MeOH, and 20 uL of which were analyzed by LC-MS (Agilent 6120 Quadrupole LC/MS) and HR-HPLC-MS analyses (Agilent 6530 Q-TOF LC/MS equipment). HPLC-MS conditions: Poroshell 120 EC-C18 2.7 um 4.6x100 mm column, flow rate 0.6 ml/min, 0.1% formic acid in water (solvent A), 0.1% formic acid in acetonitrile (solvent B), gradient: 1 min, 0.5% B; 1-30 min, 0.5-100% B; 30-35 min, 100% B. HR-HPLC-MS conditions: Poroshell 120 EC-C18 2.7 um 2.1x100 mm column, flow rate 0.25 ml/min, 0.1% formic acid in water (solvent A), 0.1% formic acid in acetonitrile (solvent B), gradient: 1 min, 0.5% B; 1-20 min, 0.5%-100% B; 20-25 min, 100% B.

### 4. *Optimization of heterologous expression by different culturing conditions*

To determine the most optimum culture condition for the highest FAA production, several media and culture conditions were explored. These include four types of media (M9, LB, 2XYT broth and Terrific broth containing 100 µg/mL of both carbenicillin and streptomycin), two aeration conditions (aerobic and anaerobic), three temperatures (20, 25 and 30°C) and two cultivation durations (72 and 120 hours). In all cases, the *ecf* producing strain was expressed and analyzed as mentioned above, and semi-quantitative production yields were assessed by comparing the resulting HPLC-MS chromatograms (**Figure S2**).

### 5. *Large scale expression, FAA isolation, purification, and structural elucidation*

The FAA products were found predominantly in the cell pellets. The best conditions for heterologous expression were determined as: terrific broth medium containing the corresponding antibiotics, 30°C temperature, 120 hours, 200 rpm. 75 L of the *ecf* harboring cell line were cultured (1 L each in baffled Fernbach flasks). Cell pellets were recovered by centrifugation, combined, suspended in 2L MeOH, and sonicated for 1 hour. Cell debris were spun down and collected supernatant was dried under vacuum using a rotary evaporator. The dried material was suspended in ethyl acetate and the organic solvent extract was dried under vacuum using a rotary evaporator. The ethyl acetate extract was fractionated by reverse phase flash column chromatography (Mega Bond Elut-C18 10g, Agilent Technology) using the following mobile phase conditions: solvent A:B (water: acetonitrile with 0.01% formic acid), and a gradient from 100% A to 100% B with 20% increment steps. Fractions containing the FAAs as identified by HPLC-MS were applied to reverse phase HPLC to purify individual FAA molecules using a fraction collector and monitoring by the mass of individual molecules (Agilent 1260, Poroshell 120 EC-C18 2.7 um 4.6x100 mm column, flow rate 0.6 ml/min, 0.1% formic acid in water (solvent A), 0.1% formic acid in acetonitrile (solvent B), gradient: 1 min, 0.5% B; 1-30 min, 0.5%-100% B; 30-35 min, 100% B). This strategy yielded < 300 µg of NAMP and NAPeP, and < 50 µg of the rest.

#### *5.1. Structural elucidation of isolated N-Acetyl-Myristoyl-Putrescine (NAMP)*

The molecular formula predicted by HR-MS for the first compound was $C_{20}H_{40}N_2O_2$ (m/z: $[M+H]^+$ Calculated $C_{20}H_{41}N_2O_2$ 341.3168, Observed 341.3172) (**Data S5, table a**). One- ($^1H$ and $^{13}C$) and two-dimensional (COSY, HSQC, and HMBC) nuclear magnetic resonance (NMR) experiments suggested the presence of a four-carbon spin system and a long-chain fatty acid substructure, comprised of two amide carbon atoms, sixteen methlylene carbon atoms, and two methyl carbon atoms. Based on the carbon chemical shift data, the four-carbon spin system is predicted to be functionalized at both ends with nitrogen atoms. HMBC correlations were observed between the methylene protons H2′ and C1′, and between the methyl protons H14 and C12, as well as between the four methylene protons (H1′ and H4′) and the two carbonyl carbons (C1 and C5’), respectively (**Data S5, Table b and Figures a1-a6**). These correlations established the structure of NAMP as an *N*-acetylated derivative of 4-aminobutamine. Based on the molecular formula and HR-LC-MS/MS fragmentation pattern, the fatty acid chain must be a fully saturated 14-carbon fatty acid. The final structure of NAMP was therefore elucidated to be *N*-Acetyl-Myristoyl-Putrescine (NAMP), and further confirmed by synthesizing a reference standard (**see below**). The 1D and 2D NMR spectra and tandem mass spectra are identical to its synthetic reference standard (**Figure S3** and **Data S5**).

#### *5.2. Structural elucidation of isolated N-Acetyl-Palmitoyl-Putrescine (NAPP)*

The molecular formula predicted by HR-HPLC-MS for the purified compound was $C_{22}H_{46}N_2O_2$ (m/z: $[M+H]^+$ Calculated 369.3481, Observed 369.3480). $^1$H-NMR spectrum (**Data S5, Figure b1, b2**) and HR-HPLC-MS/MS analysis showed a similar pattern to that of NAMP, suggesting a closely related structure varying only in the length (C16 instead of C14) of the fatty acid chain. The proposed structure (*N*-Acetyl-Palmitoyl-Putrescine) was then synthesized (**see below**), and proved to match the $^1$H-NMR, retention time, and HR-HPLC-MS/MS fragmentation pattern of the isolated molecule (**Figure S3** and **Data S5**).

#### *5.3. Structural elucidation of isolated N-Acetyl-Palmitoleoyl-Putrescine (NAPeP)*

The isolated molecule was predicted by HR-MS to have the following molecular formula: $C_{22}H_{46}N_2O_2$ (m/z: $[M+H]^+$ Calculated $C_{22}H_{47}N_2O_2$ 367.3325, Observed 367.3327). Based on the predicted molecular formula for this molecule, the fatty acid side chain was determined to be [C18:1]. One-dimensional ($^1$H and $^{13}$C) spectra were highly similar to NAMP and NAPP. The observed differences were mainly in the presence of two methylylidene carbon atoms (C-9,10; 129.4 ppm) and their protons (5.37 ppm) and the presence of two shielded methylene carbons (C-8,11; 26.7ppm) and their protons (2.07 ppm), which indicated the *Z*-unsaturated fatty acid moiety (**Data S5, Table c and Figures c1-c6**). The position of the double bond is predicted at C-9 based on the position that is most frequently seen in *E. coli* lipids, corresponding to a palmitoleic acid moiety (Mejia et al., 1999). HMBC correlations were observed between H9, H10 and C8, C11 and between olefinic protons to C9 and C10. The 4-carbon spin system with COSY correlations from H8 to H11 was also revealed. The final structure was therefore elucidated to be *N*-Acetyl-Palmitoleoyl-Putrescine (NAPeP).

#### *5.4. Structural elucidation of isolated NMP, NML and NLL*

The molecular formulae for these molecules as deduced by HR-HPLC-MS, are shown in **Data S5, Table a**. Because of the limited amounts of each of these three molecules, even from 75 L of culture (estimated <30 µg of each), structural elucidation relied mainly on HR-LC-MS/MS analysis to propose the most plausible structure, followed by total synthesis and comparison of retention times and HR-HPLC-MS/MS spectra to those of the detected molecules (**Figure S3, Figure S4**, and **Data S5**). The structures were therefore determined to be *N*-Myristoyl-Putrescine (NMP), *N*-Myristoyl-L-Lysine (NML) and *N*-Lauroyl-L-Lysine (NLL). While NMP and NML standards were synthesized here (**see below**), NLL standard was commercially available and purchased from Santa Cruz Biotech. The absolute configurations of NML and NLL were confirmed by comparing their retention times with enantiopure synthetic standards using a chiral column: Agilent 1260, Poroshell 120 Chiral-T column 2.7µm, 4.6x100 mm column, flow rate 0.8 ml/min and the following HPLC conditions (**Figure S4**).

##### 5.4.1. HPLC condition of NML

0.1% formic acid in water (solvent A), 0.1% formic acid in acetonitrile (solvent B), gradient: 1 min, 30% B; 1-30 min, 30%-60% B; 30-31 min, 60%-100% B, 31-36, 100%B).

##### 5.4.2. HPLC condition of NLL

0.1% formic acid in water (solvent A), 0.1% formic acid in acetonitrile (solvent B), gradient: 1 min, 20% B; 1-30 min, 20%-40% B; 30-31 min, 40%-100% B, 31-36, 100%B).

### 6. Chemical synthesis of FAAs

#### 6.1. Chemical synthesis of N-Myristoyl-Putrescine (NMP), N-Acetyl-Myristoyl-Putrescine (NAMP), and N-Acetyl-Palmitoyl-Putrescine NAPP (origin and catalog numbers for all reagents are shown in Key Resources Table).

For the synthesis of NMP, 0.01 mol of diaminobutane was mixed with 0.01 mol of pyridine in 10 mL of DMF and stirred for 10 min at room temperature. After transferring the mixture to an ice bath, 0.01 mol of myristoyl chloride dissolved in 10 mL DMF was added dropwise for 60 min so that the temperature of the reaction mixture does not exceed 5°C. The resulting solution was stirred for 10 h at room temperature and then poured in ice water to remove the pyridine salt (**Figure S5**) (Zhang et al., 2010)**.** The mixture was then filtered using a Buchner funnel, and the white powder obtained by filtration was washed three times with Milli-Q water and dried in a water bath. The washed powder was then dissolved in MeOH, and purified by reverse phase HPLC (Agilent 1260, with the use of a fraction collector, and monitoring by mass). Poroshell 120 EC-C18 2.7 um 4.6x100 mm column, flow rate 0.6 ml/min, 0.01% formic acid in water (solvent A), 0.01% formic acid in acetonitrile (solvent B), gradient: 1 min, 0.5% B; 1-30 min, 0.5%-100% B; 30-35 min, 100% B. For the preparation of NAMP and NAPP, the same strategy for chemical synthesis and purification was employed except for the following modifications: acetyldiamine was used instead of diaminobutane for NAMP, and palmitoyl chloride was used instead of myristoyl chloride for NAPP. Chemical structures of synthetic products were confirmed using HPLC-HR-MS/MS analysis, as well as 1D and 2D NMR (**Data S5, Figures d1-d6, e1-e7, f1-f6 and Tables d and e**).

The retention time, $^1$H-NMR and HPLC-HR-MS/MS spectra of NAMP and NAPP purified from the *ecf* heterologous expression were found to be identical to those of the synthetized standards (**Figure S3**). Retention time and HPLC-HR-MS/MS spectra of NMP from the heterologous expression were also identical to those of the synthetic standard (**Figure S6**).

#### 6.2. Chemical synthesis of N-Myristoyl-L-Lysine (NML)

100 mg Wang resins with preloaded L-lysine was incubated in DMF for 30 min. *N*-Fmoc was removed by treating with 3 mL of piperidine (20% solution in DMF) for 3 times and a duration of 10 min for each treatment, followed by several washes with DMF. Myristoyl chloride (1 equivalent) in DMF was then added and the resin suspension was shaken for 2 h at 25°C

(**Figure S3**) (Cohen et al., 2017). The FAA product was eluted from resin by treatment with trifluoroacetic acid (TFA) supplemented with 2.5% (v/v) water and 2.5% (v/v) triisopropylsilane (TIPS). TFA was removed by lyophilizing and NML was purified by reverse phase HPLC using a fraction collector and monitoring by mass (Agilent HPLC1260, Poroshell 120 EC-C18 2.7 um 4.6x100 mm column, flow rate 0.6 ml/min, 0.01% formic acid in water (solvent A), 0.01% formic acid in acetonitrile (solvent B), gradient: 1 min, 0.5% B; 1-30 min, 0.5%-100% B; 30-35 min, 100% B). The structure of the synthetic product was confirmed using HPLC-HR-MS/MS analysis, as well as 1D and 2D NMR (**Data S5, Figures g1-g6 and Table f**). Retention time and HPLC-HR-MS/MS spectra of NML from the heterologous expression is identical to those of the synthetic standard (**Figure S3** and **S4**).

### 7. *Detection of FAAs from native Clostridium sp. harboring ecf and ebf*

For seed culture preparation, a frozen scoop from the glycerol stock of *Enterocloster bolteae* ATCC BAA-613 or *Enterocloster clostridioformis* WAL-7855 was inoculated in 5 mL pre-reduced modified PYG liquid media (according to the ATCC recipe, ATCC, USA) and grown overnight at 37°C in an anaerobic chamber (70% $N_2$, 25% $CO_2$, 5% $H_2$). 250 µL of seed cultures were inoculated into 100 mL of seven different pre-reduced liquid media (modified GAM, TYG, RCM, LB, BHI, modified PYG, M17, and TSB) and incubated for another 120 hours at 37°C under the same anaerobic conditions. After 120h, the cultures were centrifuged at 3,900 rpm for 30 min, and cell pellets were suspended in 20 mL MeOH and sonicated for 30 min. Cell debris were then spun down and collected supernatant was dried under vacuum using a rotary evaporator. The extracts were resuspended in 500 µL MeOH and 10 µL were analyzed by HPLC-HR-MS/MS (**Figure S6**). Synthetic standards were also analyzed using the same conditions: Agilent 6530 Q-TOF LCMS, Poroshell 120 EC-C18 2.7 um 2.1x100 mm column, flow rate 0.25 ml/min, 0.1% formic acid in water (solvent A), 0.1% formic acid in acetonitrile (solvent B), gradient: 1 min, 0.5% B; 1-20 min, 0.5%-100% B; 20-25 min, 100% B.

### 8. *Mouse strains and colonization*

SPF C57BL/6 mice were housed by the Unit for Laboratory Animal Medicine at the University of Michigan. Female and male mice, age 7–10 weeks, were used in all experiments. All animal studies were conducted in accordance with protocols reviewed and approved by the University of Michigan Institutional Animal Care and Use Committee.
For *E. coli* infection *in vivo*, mice were pre-treated with ampicillin and streptomycin (20 mg/mouse, respectively). One day after antibiotic treatment, mice were colonized with *E. coli* strain expressing *ecf* (*E. coli* BL21_smNRPS-ecf_sfp) or an isogenic strain harboring an empty vector (*E. coli* BL21_pUC57_sfp) ($10^9$ CFU/mouse each). To induce the expression of *ecf*, mice were given drinking water containing 25 mM IPTG during the experimental period. Feces were collected from the mice, and homogenates of feces were cultured on LB agar plates with antibiotics. The number of viable bacteria was estimated by plate counting the CFUs.

#### *8.1. Dextran sulfate sodium (DSS)–induced Colitis Model*

Because mice are treated with IPTG in drinking water, DSS colitis was induced via oral gavage of 200 mg DSS (dissolved in 200 µL PBS; equivalent to 3% DSS ad lib) daily for 6 days (Kitamoto et al., 2020). The animals were monitored for weight loss during the course of experiments. Mice were euthanized on day 7, and colon weight and length were measured. The colon tissues were immediately preserved overnight in 4% paraformaldehyde, processed into paraffin-embedded tissue sections, and stained with H&E for histological assessment. A veterinary pathologist performed a blind evaluation of the histological scores as a previous study (Cooper et al., 1993). Briefly, inflammation and epithelial loss were assessed for severity based on the most severe lesion in each section (0, none; 1, mild; 2, moderate; 3, severe; 4, marked). Lesion extent was assessed as the percent of the section affected (0, 0%; 1, 1%–25%; 2, 26%–50%; 3, 51%–75%; 4, 76%–100%). The extent and severity scores for inflammation and epithelial cell loss were multiplied to give a total score for each parameter (range 0–16). The total scores for each parameter were summed to give a total colitis score (range 0–32).

#### *8.2. IL-10–deficient Colitis Model*

Germ-free IL-10-deficient mice were colonized with either the *E. coli* expressing *ecf* (*E. coli* BL21_smNRPS-ecf_sfp) or the isogenic strain harboring an empty vector control (*E. coli* BL21_pUC57_sfp), and maintained for 4 weeks. Mice were given drinking water containing 25 mM IPTG during the experimental period. Mice were euthanized 4 weeks after colonization, and colon weight and length were measured.

#### *8.3. Intestinal permeability assay*

The intestinal permeability assay was performed using FITC-dextran at the end of the experiment as previously described (Sugihara et al., 2022). Briefly, the food was removed for 4 hours and then 0.6 mg/g body weight of 4 kDa FITC-dextran (FD4, Sigma-Aldrich) was administered by oral gavage. Blood was collected after 4 hours and fluorescence intensity was measured (excitation, 485 nm; emission, 520 nm). FITC-dextran concentrations were determined using a standard curve generated by serial dilution of FITC-dextran.

#### *8.4. Quantitative real-time PCR*

Colonic tissue samples were collected, and RNA was extracted using E.Z.N.A. Total RNA Kit I (Omega Bio-tek). cDNA was synthesized using a High-Capacity RNA-to-cDNA Kit (Thermo Fisher Scientific). qPCR was performed using a Radiant SYBR Green Lo-ROX qPCR Kit (Alkali Scientific). The relative expression of the target genes was calculated using β-actin as a reference.

### *9. Toxicity of fatty acid amides against human colonic epithelial cells*

Caco-2 were cultured in Dulbecco's Modified Eagle Medium (DMEM), supplemented with fetal bovine serum (FBS) and antibiotics (i.e., penicillin and streptomycin) in Nunclon Delta-treated dishes in at 37°C and 5% $CO_2$ until obtaining the required number of cells. On the day of the experiment, the cells were detached using Trypsin-EDTA (0.25%), then washed using Dulbecco's phosphate-buffered saline (DPBS). Cells were resuspended in DMEM and counted using Trypan Blue (Cell viability = 96%). Cells were seeded in cell culture-treated 96-well clear polystyrene microplates at 5,000 cells per well in a 100 µL. In each well, reagents of RealTime-Glo™ MT cell viability assay kit (Promega) were added to monitor cell viability over time. Microplates were incubated at 37°C and 5% $CO_2$. After 24 hours, cells were treated with the FAAs at concentrations 1–32 µM then incubated further at 37°C and 5% $CO_2$. Because the FAAs were dissolved in methanol, final methanol concentration was kept at 1%. Untreated, methanol-treated, and 0.1% Triton-treated cells were included as controls. Final cell viability was measured after 24 hours of incubation to assess the cytotoxicity of the FAAs. Luminescence in relative light unit (RLU) was measured using a Tecan Spark® multimode microplate reader.

### 10. Computational analysis of metagenomic and metatranscriptomic samples

#### 10.1. Pre-processing of metagenomic and metatranscriptomic data

Sequence Read Archive (SRA) data of Illumina reads were downloaded for all datasets using their corresponding NCBI BioProject, then FASTQ was extracted using the SRA Toolkit. A Data Transfer Agreement was established to obtain the raw data of the IBD Plexus, Crohn's & Colitis Foundation initiatives, i.e., SPARC and RISK. Raw reads were filtered using PRINSEQ (Schmieder and Edwards, 2011), with the following parameters: reads were required to have an average quality score of 30, and less than two percent of undetermined (N) bases, otherwise they were discarded; reads were trimmed on both ends to remove bases with less than a quality score of 30. If trimming ended up producing a read of less than half of the initial read length, this read was also discarded(Schmieder and Edwards, 2011). SPAdes was then used to assemble filtered reads of metagenomes for each individual subject (both pairs and singletons), with default parameters (Bankevich et al., 2012).

#### 10.2. Creating a non-redundant representative set of BGCs:

To detect biosynthetic gene clusters (BGCs), we used antiSMASH v5.1.1 with the following parameters: --smcogs --clusterblast” on scaffolds that are 5 Kbps or longer for each sample’s SPAdes assembly. antiSMASH outputs a GenBank file for each annotated BGC (n = 80,665). BGCs were filtered to keep only those with no ambiguous nucleotides (n = 70,078). Using the GenBank file for each BGC, we used Python and Python module Biopython (within seqIO_extract.py) to parse out each identified BGC and its nucleotide sequence to create a master FASTA file of all BGCs.

To assemble a non-redundant set of BGCs, we launched blastn on the master FASTA file of BGCs against itself. Using cutoffs of 90% percent DNA sequence identity and 90% breadth of coverage, BGCs were de-replicated so that truncated, duplicated, or BGCs of shorter length that match BGCs of longer length were removed. The resulting non-redundant set of BGCs included 10,060 BGCs.

To determine the taxonomy of the organisms harboring the identified BGCs, we used blastn to query the nucleic acid sequences of the identified, de-replicated BGCs against all genomic sequences from the NCBI RefSeq Genome and Nucleotide databases (04-29-2022). We ran blastn with the following parameters: “-evalue 0.00001 -max_target_seqs 1000 -perc_identity 80”. Then, using ≥ 90% percent DNA sequence identity and ≥ 50% breadth of coverage, we identified the best hit for each BGC. If multiple hits passed our cutoffs, we filtered first by taking the hit with the highest bit score, then highest percent DNA sequence identity, then highest breadth of coverage, and if there were still ties, we assigned the first hit to appear in the BLAST results to the BGC. If a BGC has no BLAST hits that passed our cutoff, we classified the taxonomy as “Unassigned”.

Chemical classes of the non-redundant set of BGCs were extracted from the GenBank files. For plotting purposes, we reduced the total number of BGC classes and grouped the BGCs based on three major BGC classes: RiPP, NRPS, PKS, and Others. The “RiPP” class membership included: microcin, bacteriocin, lantipeptide, LAP, RaS-RiPP, lassopeptide, bacteriocin-lantipeptide, sactipeptide, thiopeptide, bacteriocin-proteusin, glycocin, microcin-lassopeptide, proteusin, and proteusin-bacteriocin. The "NRPS" class membership included: NRPS, and NRPS-like. The "PKS" class membership included: PKS, T1PKS, T2PKS, T3PKS, PKS-like, transAT-PKS-like, and transAT-PKS. Lastly, the "Other" class membership included: terpene, arylpolyene, resorcinol, phosphonate, oligosaccharide, ectoine, betalactone, butyrolactone, ladderane, siderophore, hserlactone, NAGGN, bacteriocin-arylpolyene, amglyccycl, ladderane-arylpolyene, phenazine, CDPS, hglE-KS, furan, indole, nucleoside, other, and hybrids.

To identify open reading frames (ORFs) from the non-redundant set of BGCs, we parsed the GenBank files and used the "gene_kind" element to annotate all ORFs (n = 199,333). Of all ORFs, 15,147 core biosynthetic genes were identified. To assemble a non-redundant set of CB-ORFs, we used blastn on the FASTA file of all CB-ORFs. Using cutoffs of 90% percent DNA sequence identity and 90% breadth of coverage, we collated 5,157 unique CB-ORFs, which were quantified and used for all statistical analyses (see below). For the purpose of calculating BGC abundance, prevalence, and enrichment statistics (In relation to **Figure 2A**, **2B**, and **2D**), and because a CB-ORF can be shared between different BGCs, we first performed all calculations based on the CB-ORFs as described below, generated a representative set of BGCs based on their CB-ORF content (n = 3,328), and finally used this set to infer BGC statistics (allowing us to avoid counting a CB-ORF (or clustered CB-ORFs) more than once) (**Data S1**).

#### *10.3. Quantifying the abundance of CB-ORFs in metagenomic and metatranscriptomic samples*

To quantify the abundance of core biosynthetic ORFs, we first used Bowtie 2 with the following parameters: "--end-to-end --very-sensitive --score-min L,-0.6,-0.3." to map quality filtered reads from metagenomic and metatranscriptomic samples to the CB-ORFs database. We then parse the Bowtie 2 output to calculate the following metrics: *abundance* in Reads Per Kilobase per Millions of sequenced reads (RPKM), and *breadth of coverage* as the total percentage of the CB-ORF's length that is covered by sample reads. We considered a CB-ORF to be present in a sample if it is covered at any breadth of coverage. This method allowed us to detect CB-ORFs and BGCs in samples where they are at very low abundance.

#### *10.4. Identifying and prioritizing enriched CB-ORFs and BGCs in metagenomic samples using statistics and machine learning algorithm*

We used two-samples proportion z-test followed by Bonferroni correction to compare the prevalence of each CB-ORF in a pairwise manner, i.e., HC versus CD and HC versus UC. We used the following cutoffs: adjusted P cutoff of $\leq 0.01$, a prevalence difference of $\geq 10$ to identify differentially prevalent CB-ORFs. We then inferred BGCs enrichment using the representative set of BGCs (explained above). At the BGC level, enrichment criteria were that (1) a BGC must harbor one or more differentially prevalent CB-ORF(s), and (2) a BGC cannot harbor multiple CB-ORFs enriched in opposite directions (This occurred in only one case for a BGC with two CB-ORFs, one of which was enriched in UC, while the other was enriched in HC) (**Data S1**).

To test the ability of a random forest classifier to predict the disease status of metagenomic samples based on the abundance of CB-ORFs, we split the samples (only one sample per subject) into training and testing sets. The training set did not include any of the testing set samples. The testing set comprised 20% of all samples and were not seen by the classifier, nor contributed unique BGCs (and correspondingly their unique CB-ORFs) that allowed their identification. The model was trained then tested (number of trees = 500). The model was evaluated with Receiver Operating Characteristic (ROC) area under the curve (AUC) value, Precision-Recall (PR) AUC value, and out-of-bag estimate of error rate, as evaluation metrics. To validate the robustness of the model, we used a five-fold cross-validation approach, where we performed five iterations of sampling 20% of the dataset, removing their unique CB-ORFs from all features, then training on the remaining 80% of the samples and testing on the never seen 20% (**Data S1**). To determine which predictors (i.e., CB-ORFs) were most important in driving predictions (i.e., CD vs. HC), we used Boruta's algorithm (Kursa and Rudnicki, 2010), an established feature selection algorithm.

#### *10.5. Calculation of the abundance and enrichment of ebf/ecf BGCs in metagenomic and metatranscriptomic samples*

To calculate the abundance and prevalence of the *ebf and ecf* BGCs in all samples and due to the high sequence similarity between them (~92% identity at the nucleotide level), we quantified these BGCs separately using the same Bowtie 2 pipeline and parameters described above. We used two-samples proportion z-test, followed by Bonferroni correction to determine statistical significance regarding the prevalence of *ebf* and *ecf* together in different groups. If multiple samples existed per subject, the detection of *ebf* or *ecf* in any metagenomic sample from the subject was used as the minimum criterion to deem this subject as positive for *ebf* or *ecf* presence. To determine the statistical significance comparing the abundance of *ebf* and *ecf*, we used Kruskal-Wallis test, followed by Dunn's multiple comparison and adjusted by Bonferroni correction. If multiple samples existed per subject, the subject-based abundance of *ebf* or *ecf* used in this analysis was based on the average abundance across all samples from the same subject. To test the enrichment of *ebf* and *ecf* across countries, we used a meta-analysis random effects model and effect sizes were calculated using a meta-analysis of log ratio of means. Similarly to the previous analysis, if multiple samples existed per subject, the subject-based abundance of *ebf* or *ecf* used in this analysis was based on the average abundance across all samples from the same subject.

To evaluate the expression of *ebf* and *ecf* in samples, we used paired metagenomic and metatranscriptomic samples from the iHMP-IBD dataset and another publicly available cohort (Abu-Ali et al., 2018; Lloyd-Price et al., 2019; Schirmer et al., 2018). We quantified their abundance in samples using the same quantification method (described above). Two-samples proportion z-test, followed by Bonferroni correction was used to determine statistical significance regarding the prevalence of *ebf* and *ecf* in different groups. If multiple paired samples existed per subject, the detection of *ebf* or *ecf* in any metatranscriptomic sample from the subject was used as the minimum criterion to deem this subject as positive for *ebf* or *ecf* expression.

#### *10.6. Annotating previously characterized BGCs using the MIBiG database*

To identify if any of the enriched BGCs matched already characterized BGCs, we leveraged the MIBiG 3.1 database (Terlouw et al., 2023). We downloaded GenBank files of all 2,502 BGCs in the database, then parsed their nucleotide sequences. Next, we used blastn to determine if CB-ORFs from our database matched any of the 2,502 BGCs in the MIBiG 3.1 database using the following cutoffs: percent DNA sequence identity ≥ 90% and ≥ 90% breadth of coverage of the CB-ORF by a MIBiG BGC (**Figure 2E**). Finally, to determine if our set of BGCs match any of the MIBiG 3.1 database BGCs we used blastn with the following cutoffs: percent DNA sequence identity ≥ 90% and ≥ 50% breadth of coverage of the MIBiG 3.1 BGC by one of our BGCs (**Data S1**).

#### *10.7. Building a phylogenetic tree of Clostridia strains*

To build a phylogenetic tree of strain from the class Clostridia with publicly available genomes, we first downloaded all 8,427 genome assemblies of Clostridia strains from the RefSeq database (08-13-2021). We used PhyloPhlAn (release 28 April 2020) (Asnicar et al., 2020) to build the tree based on a set of 400 universal marker genes. We then used tblastn to identify and annotate Clostridia strains harboring an *ebf/ecf* homologous BGC based on the presence of the two CB-ORFs (C and A domains) of *ebf.* We used the following cutoffs for tblastn results: E-value ≤ 0.001, percent sequence identity ≥ 90% and breadth of coverage ≥ 50% of the CB-ORF by the genome. To annotate Clostridia strains harboring previously characterized FAA BGCs, we used "Supplementary Table 3" in Chang et al. to identify the matching strain, as well as the protein accessions to confirm BGC presence in our Clostridia strains using tblastn (Chang et al., 2021).

### DATA AND SOFTWARE AVAILABILITY

Data S1, Data S2, Data S3, and Data S4 contain all the computational analysis outputs using standard tools (described in the Resources Table; and software parameters described in the Methods). Data S5 contains all NMR spectra used for the structural elucidation of the compounds discovered in this study. All datasets used in this study are publicly available and can be accessed through NCBI BioProject database (https://www.ncbi.nlm.nih.gov/bioproject/) using the following accession numbers: PRJNA398089, PRJNA389280, PRJNA487636, PRJNA384246, PRJNA385949, PRJNA321058, SRP057027 (can be accessed through NCBI SRA database https://www.ncbi.nlm.nih.gov/sra/), PRJNA237362, PRJNA685168, PRJNA400072, PRJNA942468, PRJEB5224, PRJEB1220, PRJEB2054, PRJEB35587, PRJNA429990, PRJEB15371, PRJNA532645, PRJNA354235). IBD Plexus datasets can be requested through the Crohn's & Colitis Foundation (https://www.crohnscolitisfoundation.org/research/grants-fellowships/ibd-plexus). Software used in this study is described in the Resource Table. This paper does not report original code.
